# Using reciprocally retained gene families to detect whole-genome multiplications in plants

**DOI:** 10.1101/2025.04.18.649489

**Authors:** Setareh Tasdighian, Cecilia Sensalari, Steven Maere

**Author notes:** **_Corresponding author_**_: E-mail:_. These authors contributed equally to this work (Setareh Tasdighian, Cecilia Sensalari).

## Abstract

Traces of ancient whole-genome multiplications (WGMs) have been observed all over the plant kingdom and have been associated with various evolutionary processes, such as increased evolvability, speciation, adaptation to changing environments, domestication and the origin of evolutionary novelties. However, understanding the impact of WGMs on plant evolution requires accurate detection of WGM events, which is challenging because of rapid signal erosion due to genome rearrangements, sequence divergence and the occurrence of additional large- and small-scale duplications (SSDs). Here, we investigate whether reciprocally retained gene families (RR GFs), i.e. GFs that preferentially expand through WGM and rarely undergo SSDs, can be used as WGM markers. Using stochastic birth-death (BD) modeling of GF gene count data to test for WGM presence or absence, we demonstrate that strongly RR GFs have higher power to detect true WGMs and to reject false WGMs than non-RR GFs. However, none of the RR GFs is a perfect WGM marker on its own, and different GFs perform better in different plant clades, prohibiting the use of a fixed small set of RR GFs as WGM markers across all angiosperms. Instead, we show that using an extended set of RR GFs rather than whole paranomes as input for BD and *K*_S_ distribution modeling approaches leads to improved WGM detection performance.

## 1 Introduction

Traces of ancient whole-genome multiplications (WGMs) have been uncovered in the genomes of species from all eukaryotic kingdoms. In the animal kingdom, a whole-genome duplication (WGD) took place in the ancestor of all vertebrates (Dehal and Boore, 2005), followed by a second WGD in the jawed vertebrate ancestor and a whole-genome triplication (WGT) in the jawless vertebrate ancestor (Marlétaz *et al*., 2024; Nakatani *et al*., 2021), a third WGD in the teleost lineage and a fourth WGD in salmonids (Jaillon *et al*., 2004; Lien *et al*., 2016). Evidence for ancient WGMs has also been found in the genomes of e.g. baker’s yeast (Kellis *et al*., 2004; Wolfe and Shields, 1997), the ciliate *Paramecium tetraurelia* (Aury *et al*., 2006) and the bdelloid rotifer *Adineta vaga* (Flot *et al*., 2013). In the plant kingdom, ancient WGMs and current polyploidy are very common. Ancient WGMs have been hypothesized at the base of the seed plant and angiosperm clades (Jiao *et al*. (2011), but see also Ruprecht *et al*. (2017)), the base of the core eudicots (Jaillon *et al*., 2007; Chanderbali *et al*., 2022) and early in the monocot lineage (Jiao *et al*., 2014; Ming *et al*., 2015), and many plant lineages have undergone additional WGMs in their evolutionary history (Van de Peer *et al*., 2009, 2017; Leebens-Mack *et al*., 2019; Cui *et al*., 2006).

WGMs in plants and other organisms have repeatedly been associated with increased speciation (Tank *et al*., 2015; Schranz *et al*., 2012), adaptation to biotic and abiotic stress factors and environmental instability (Chao *et al*., 2013; Van de Peer *et al*., 2020), adaptation to climate change and survival of catastrophic events (Vanneste *et al*., 2014; Fawcett *et al*., 2009; Sessa, 2019; Cai *et al*., 2019; Lohaus and Van de Peer, 2016; Van de Peer *et al*., 2020) and (elaboration of) evolutionary innovations (Van de Peer *et al*., 2009; De Bodt *et al*., 2005; Soltis and Soltis, 2016; Fawcett *et al*., 2013). Most of these hypotheses are still strongly debated (Van de Peer *et al*., 2009; Soltis and Soltis, 2016; Lohaus and Van de Peer, 2016; Van de Peer *et al*., 2017), not in the least because the occurrence and exact phylogenetic placement of many hypothesized WGMs are still uncertain and many other WGMs likely remain undetected (Ruprecht *et al*., 2017; Tiley *et al*., 2018, 2016; Zwaenepoel *et al*., 2019).

Three main methods are used for WGM inference: detection of syntenic blocks of duplicated genes within and across species that may be WGM remnants (Proost *et al*., 2011; Lyons and Freeling, 2008), detection of peaks in *K*_S_ distributions (i.e. distributions of the number of synonymous substitutions per synonymous site in paralog pairs, used as a proxy for age since duplication) that indicate sudden duplication bursts (Maere *et al*., 2005; Sensalari *et al*., 2021; Lynch and Conery, 2000; Blanc and Wolfe, 2004b; Cui *et al*., 2006; Zwaenepoel and Van de Peer, 2018), and detection of duplication bursts through gene tree-species tree reconciliation (Jiao *et al*., 2011; Rabier *et al*., 2014; Li *et al*., 2015; Thomas *et al*., 2017; Yang *et al*., 2018; McKain *et al*., 2016; Zwaenepoel and Van de Peer, 2019). All three methods have limitations and are challenged by extensive gene loss eroding the WGM signal over evolutionary time. The identification of WGM remnants through synteny analysis is highly dependent on the availability of good genome assemblies, and older events become progressively more challenging to detect because of genome rearrangements, gene loss and duplicate divergence that weaken the syntenic signal. The inference of WGMs through mixture modeling of *K*_S_ distributions suffers from *K*_S_ stochasticity and saturation effects and is prone to overfitting (Vanneste *et al*., 2013; Tiley *et al*., 2018; Zwaenepoel *et al*., 2019). Gene tree-species tree reconciliation methods require choosing an arbitrary threshold for the number of duplications on a branch to make WGM calls, and are sensitive to taxon sampling, uncertainties in the gene and species trees and heterogeneity in the small-scale duplication (SSD) and loss rates across the species tree (Zwaenepoel *et al*., 2019). Both *K*_S_-based and reconciliation-based methods may give misleading results for allopolyploidies (Thomas *et al*., 2017). More recently, stochastic birth-death (BD) models have also been used for WGM inference (Rabier *et al*., 2014; Tiley *et al*., 2016; Zwaenepoel and Van de Peer, 2020). These BD models model the gene count profiles of gene families across a species tree as a function of a small-scale gene duplication rate (birth rate), a gene loss rate (death rate) and the presence of WGMs. Rabier *et al*. (2014) reported that BD modeling of gene count profiles outperformed gene tree–species tree reconciliation for WGM detection, despite using less information. Tiley *et al*. (2016) tested the BD model of Rabier *et al*. (2014) on a set of 19 WGMs in the angiosperms and found that older WGMs and repeated WGMs on the same branch were more difficult to detect. Zwaenepoel and Van de Peer (2020) reported similar findings using a BD modeling approach accounting for uncertainties in gene tree topology and gene tree-species tree reconciliation, and noted that taxon sampling can influence WGM inference results.

*K*_S_-based methods, reconciliation-based methods and and gene count methods generally use whole-species paranomes to infer WGMs. However, not all gene families (GFs) respond in the same way to WGM. Many GFs lose at least part of their duplicates in the turbulent period of genomic shock and cytological rediploidization that often follows WGM (Li *et al*., 2021). So-called ‘single-copy’ GFs, often containing highly conserved genes involved in essential cellular housekeeping functions, generally return to single-copy status after WGMs (De Smet *et al*., 2013). Other GFs are preferentially retained in duplicate after WGM (Maere *et al*., 2005; Papp *et al*., 2003; Seoighe and Gehring, 2004; Blanc and Wolfe, 2004a). Interestingly, many of the GFs exhibiting preferential duplicate retention after WGM also exhibit preferential duplicate loss after SSD, giving rise to a ‘reciprocal retention’ pattern after WGM and SSD (Maere *et al*., 2005; Freeling and Thomas, 2006; Freeling, 2009; Tasdighian *et al*., 2017). This reciprocal retention pattern is thought to be due to dosage balance effects acting on the genes concerned (Maere *et al*., 2005; Birchler and Veitia, 2007, 2012; Freeling and Thomas, 2006; Freeling, 2009; Tasdighian *et al*., 2017). Many regulatory pathways and protein complexes for instance are dosage balance-sensitive, requiring a precise stoichiometric balance of interacting components to function optimally (Papp *et al*., 2003; Veitia, 2002; Veitia *et al*., 2008; Birchler *et al*., 2001). SSD of dosage balance-sensitive genes without their interactors would create an imbalance and hence is selected against. WGM on the other hand preserves dosage balance by duplicating all genes simultaneously, and instead the loss of dosage balance-sensitive gene duplicates after WGM is selected against because it would lead to dosage imbalance (Freeling and Thomas, 2006; Freeling, 2009; Birchler and Veitia, 2007, 2012; Papp *et al*., 2003; Veitia *et al*., 2008; Makino and McLysaght, 2010). Tasdighian *et al*. (2017) investigated which angiosperm GFs exhibit the strongest reciprocal retention signal, and found that strongly reciprocally retained gene families (RR GFs) indeed exhibit several hallmarks of dosage balance sensitivity, such as stronger constraints on sequence, expression and function divergence than weakly RR GFs, and a functional enrichment for processes more likely to be dosage balance-sensitive such as transcriptional regulation, signal transduction and development.

The fact that not all GFs respond in the same way to WGM implies that not all GFs may be equally valuable from a WGM inference perspective. Single-copy gene families for instance are expected to contain very little WGM signal, and also other GFs may contain more noise than signal if they incurred considerable WGM duplicate loss or exhibit substantial SSD activity, or both. RR GFs on the other hand should contain more WGM signal than other GFs. Indeed, by virtue of preferentially retaining WGM duplicates and rarely retaining SSDs, the size of a strongly RR GF in a particular species should reflect its WGM history. In a perfectly RR GF, duplicates by definition exclusively originate through WGM and are retained indefinitely, so that the number of GF members in a given lineage should perfectly reflect the number and type of WGMs in this lineage. Although none of the GFs investigated in Tasdighian *et al*. (2017) were perfectly reciprocally retained, strongly yet imperfectly RR GFs may also have value as WGM markers, alone or in combination. This idea is conceptually similar to the old idea of using the number of *Hox* gene clusters to infer WGMs in animal lineages (Garcia-Fernàndez and Holland, 1994; Schughart *et al*., 1989).

Here, we test whether strongly RR GFs in angiosperms have more power than weakly RR GFs to detect WGMs, and hence could serve as WGM markers. We expand on a previously developed BD modeling approach (Tasdighian *et al*., 2017) to assess the WGM detection power of individual GFs in angiosperms, and find that strongly RR GFs indeed have more power to detect true WGMs and reject false WGMs than weakly RR GFs. However, no single GF detects all WGMs and hence qualifies as a perfect WGM marker, and the performance of any given GF depends on the plant clade(s) being analyzed, precluding the use of a small set of marker GFs to detect WGMs across the entire angiosperm clade. As an alternative, we explore the use of extended sets of RR GFs instead of whole paranomes in the BD modeling approach of Rabier *et al*. (2014) and in *K*_S_ distribution modeling, and we show that this improves WGM detection performance by eliminating the influence of non-RR GFs that contain more noise than WGM signal.

## 2 Results

### 2.1 Detection of true WGMs in a 37-species angiosperm phylogeny

In previous work (Tasdighian *et al*., 2017), we ranked 9178 core GFs in the angiosperms according to their reciprocal retention strength (a proxy for dosage-balance sensitivity). To assess the power of strongly RR GFs for WGM detection in comparison with weakly RR GFs, we focused on the top-100 most RR GFs and the bottom-50 least RR GFs (bottom-50 instead of bottom-100 because the pattern was already clear after 50 GFs and we did not want to waste computational resources on additional ‘uninteresting’ GFs). We assessed the power of each of these GFs to detect each of 20 well-supported WGMs in an angiosperm phylogeny encompassing 37 species (Fig. 1 and Suppl. Tab. S1), given the GFs’ gene count profiles across the phylogeny (Tasdighian *et al*., 2017). For this, we used a birth-death (BD) model that infers, given a tree with embedded WGM events and the size of a given GF for all species at the tree tips, maximum-likelihood (ML) estimates for a parameter *λ* that captures both the rate of gene duplication through SSD and the rate of gene loss after SSD and WGM, and for the GF size at the root of the phylogeny (see Methods and Tasdighian *et al*. (2017)). Strongly RR GFs yield low *λ* estimates on a tree with all bona fide WGMs (and only those) present. For any given GF and WGM, we estimated *λ* and the GF size at the root of the 37-species tree in the presence and absence of the WGM (with the other 19 ‘true’ WGMs considered present), and then used the corresponding model likelihoods to perform two complementary likelihood ratio tests (cLRTs): one in which the null hypothesis (*H*_0_) is that the WGM is present and one in which *H*_0_ states that the WGM is absent (see Methods). These cLRTs can give four possible outcomes: when *H*_0_ = *WGM presence* is not rejected and *H*_0_ = *WGM absence* is rejected with FDR-corrected *p*(= *q*) ≤ 0.10, the WGM is detected; when *H*_0_ = *WGM presence* is rejected and *H*_0_ = *WGM absence* is not rejected, the WGM is rejected; when both *H*_0_ are not rejected, the tests are inconclusive; and when both *H*_0_ are rejected, the model does not fit the data well.

**Figure 1:**
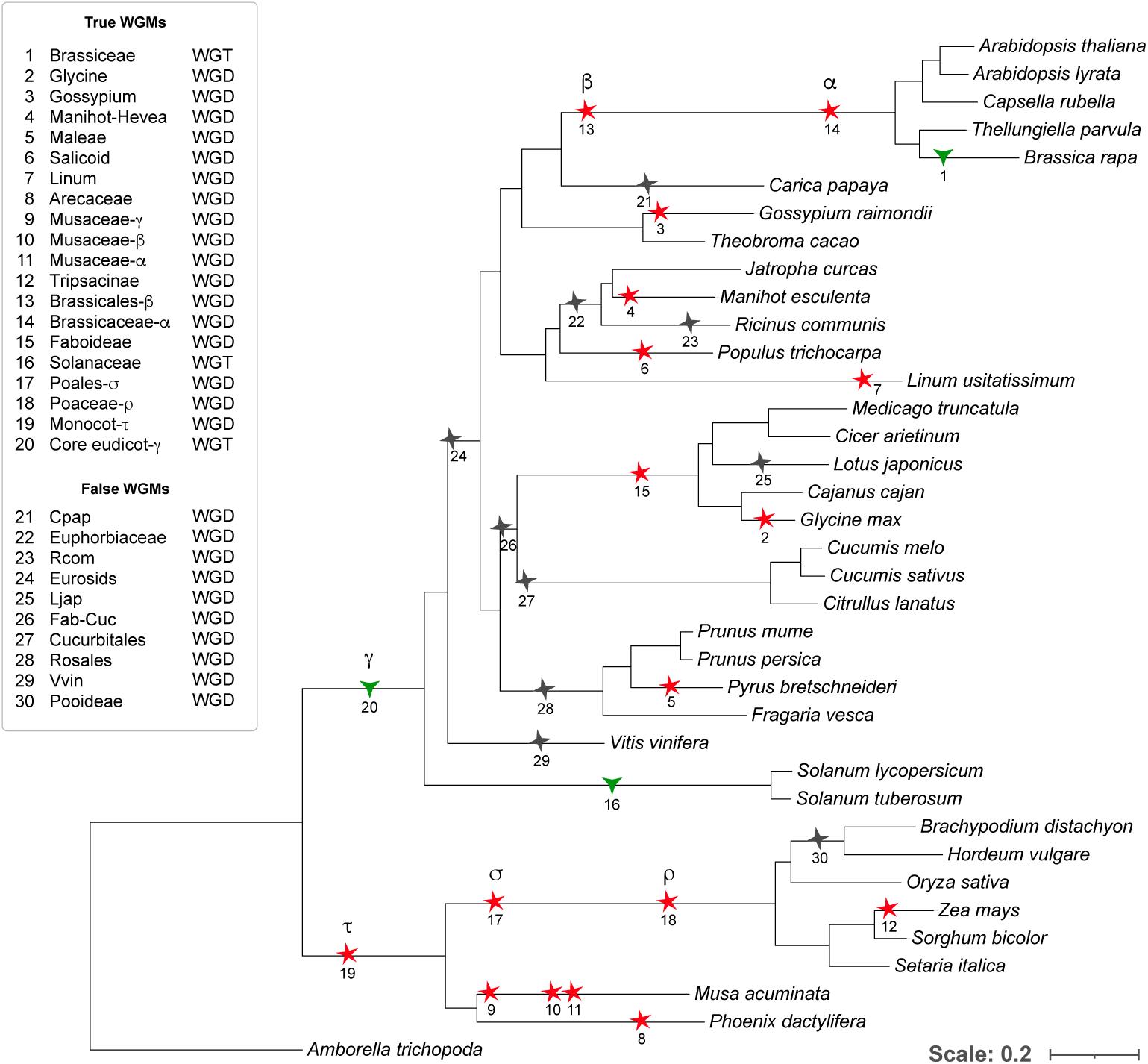
Phylogeny of the 37 angiosperms and 20 WGMs used in this study (Tasdighian *et al*., 2017). ‘True’ WGMs, i.e. WGMs that are well-supported in literature, are marked as red five-pointed stars (WGDs) or green three-pointed stars (WGTs); ‘False’ WGDs, i.e. WGMs without strong literature support, are marked as gray four-pointed stars. The tree was rendered using iTOL (Letunic and Bork, 2021).

The top-100 GFs (estimated optimal *λ* (*λ*\*) in the range [0.3288, 0.4793], Suppl. Data S1A) detected up to 19 of the 20 true WGMs in the tree, while the bottom-50 GFs (*λ*\* range [2.0026, 4.2901]) detected barely any WGMs (Fig. 2 and Suppl. Data S2A, full cLRT results can be found in Tasdighian *et al*. (2025)). 30.5% (610/2000) versus 1.8% (18/1000) of the cLRTs detect a true WGM for the top-100 and bottom-50 GFs, respectively, indicating that strongly RR GFs have higher power to detect true WGMs, as expected. However, the top-100 GFs also reject the presence of true WGMs more often than the bottom-50 GFs (11.55% (231/2000) versus 0.7% (7/1000) of the cLRTs). 97.5% of the cLRTs for the bottom-50 GFs are indecisive, compared to 54.3% of the cLRTs for the top-100 GFs. Interestingly, a limited number (3.65%) of cLRTs for the top-100 GFs reject both the presence and the absence of a WGM, indicating a bad model fit. These results indicate that strongly RR GFs are generally more decisive on the presence or absence of a true WGM than weakly RR GFs. There are however strong differences in performance across the top-100 GFs, with some GFs (GF71, GF72, GF87, GF95) only detecting a single WGM and one GF, GF7, detecting none and rejecting 13 out of 20 WGMs instead, while others (GF0, GF3, GF4 and GF5) each detect at least 15 out of 20 WGMs (Fig. 2 and Suppl. Data S2A). All 12 GFs detecting more than half of the 20 real WGMs are found in the top-30. Top GFs further down the ranking often reject more WGMs than they detect.

**Figure 2:**
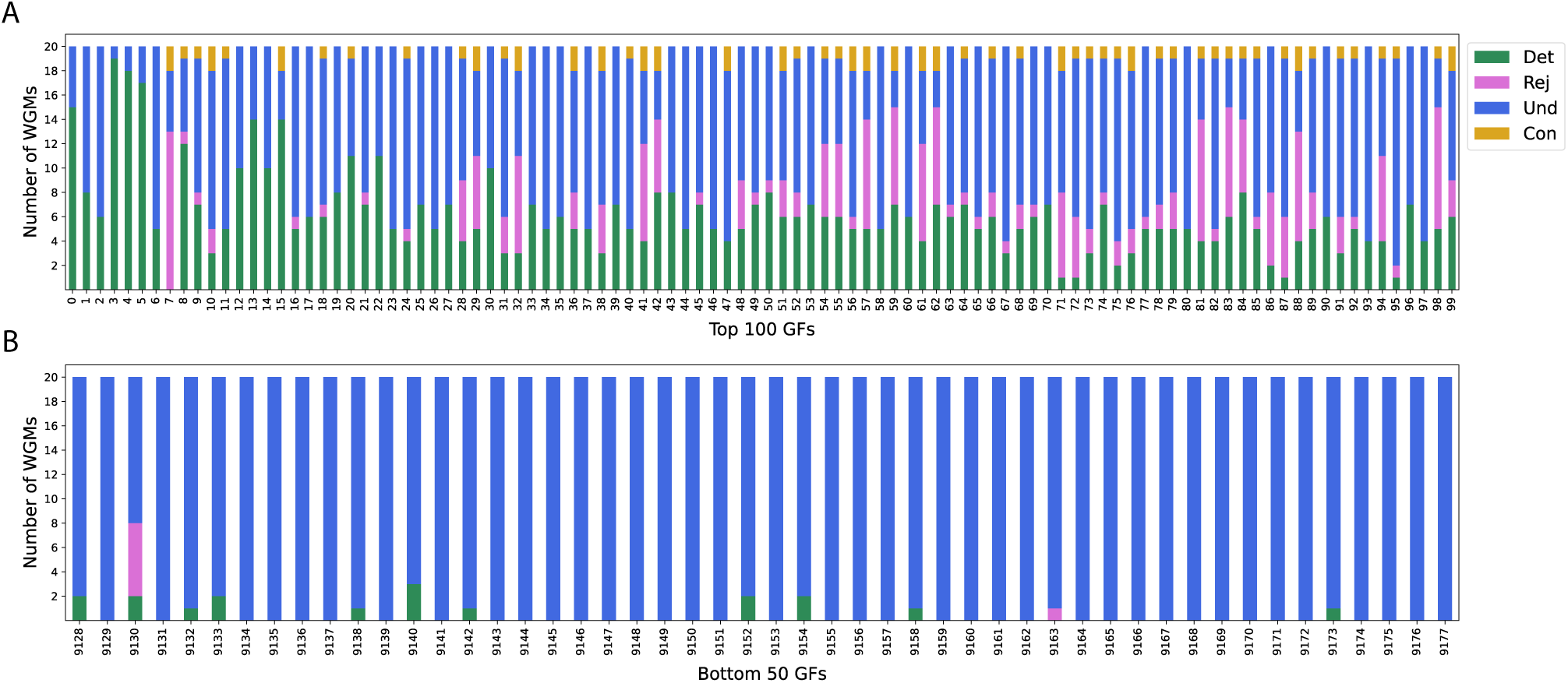
Inference performance of individual gene families belonging to the top-100 GFs (A) and bottom-50 GFs (B). The number of WGMs detected and rejected by a given GF are indicated in green and magenta, respectively. The number of WGMs with undecided cLRT outcome (both *H*_0_ not rejected) is indicated in blue, and the number of WGMs with conflicting cLRT outcomes (both *H*_0_ rejected) in yellow. See Suppl. Data S2A-D.

On the other hand, some WGMs are detected more efficiently than others by the top-100 GFs. All WGMs on the tip branches of the 37-species tree, except the repeated WGMs on the *Musa acuminata* branch, are detected by at least 20 of the top-100 GFs, and six out of 12 tip-branch WGMs are detected by more than half of the top-100 GFs (Fig. 3 and Suppl. Data S2B). The low detection and high rejection of the WGDs on the *M. acuminata* branch is likely because gene loss after repeated WGMs may be higher and the BD model can more easily compensate for the modeled loss of a WGM if there are other WGMs on the same branch (see below).Compared to WGMs on tip branches, WGMs on deeper branches are generally more difficult to detect. In particular, the oldest WGMs in the tree – the *τ* WGD early in the monocot lineage and the *γ* WGT at the base of the core eudicots – are detected by 0 and 4 of the top-100 GFs and rejected by 12 and 37 of the top-100 GFs, respectively, rendering their net detection scores across the top-100 GFs negative (Fig. 4). Surprisingly, also some of the more recent WGMs on internal branches have negative net detection scores, such as the *α* WGD in the Brassicaceae lineage, the *ρ* WGD in the Poaceae and the Solanaceae WGT. For the *α* and *ρ* WGDs, as for the WGDs in the *M. acuminata* lineage, this is likely due to the presence of a second WGD on the same branch (*β* and *σ*, respectively) that may compensate for the loss of the first. Two scenarios can be envisioned. Either the modeled loss of one of several WGMs on the same branch leads to a reduction of *λ*, which would lead to compensatory lower losses after the other WGM(s) on the branch, but would also lead to lower losses after other WGMs and lower SSD gain and loss that may unfavorably impact the modeled likelihood. On the other hand, an increase in *λ* or root size upon modeling the loss of one of several WGMs on the same branch could give the model the opportunity to reach a higher gene count before the remaining WGM(s), which would then be amplified by these WGM(s) to compensate for the removed WGM. The *λ* decrease scenario is preferred by most of the top-100 GFs (Suppl. Fig. S1).

**Figure 3:**
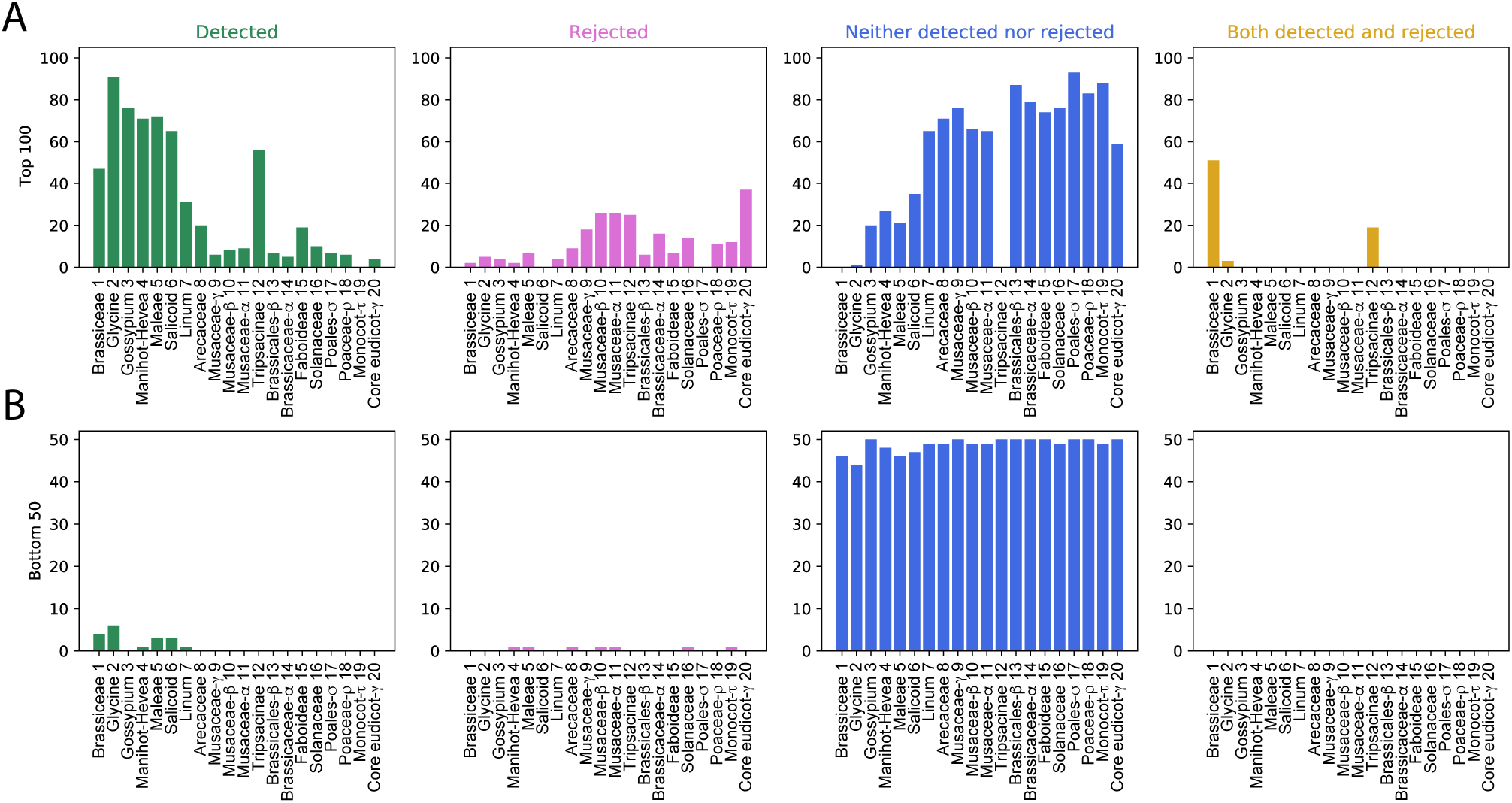
Per-WGM inference performance of the 20 well-supported WGMs by the top-100 GFs (A) and the bottom-50 GFs (B). See Suppl. Data S2A-D.

**Figure 4:**
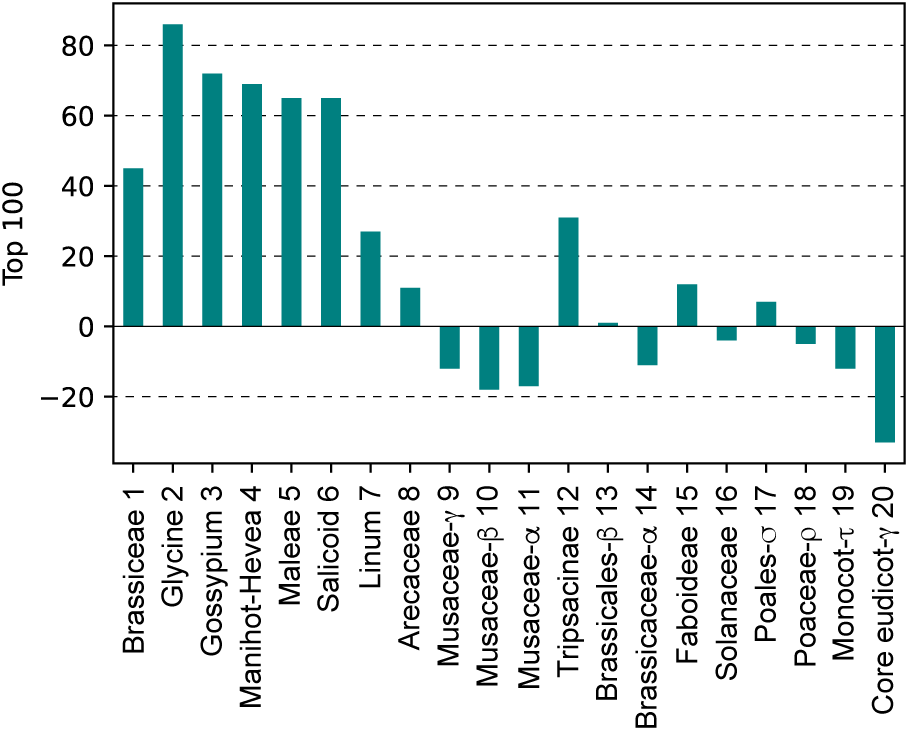
Net detection score for each WGM. The net detection score for a given WGM was calculated as the number of top-100 GFs detecting the WGM minus the number of top-100 GFs rejecting the WGM. Undecided and conflicting cLRT outcomes were ignored as they do not favor either presence or absence of the WGM concerned.

Of the eight internal WGMs (including *τ* and *γ*), only the Faboideae WGD is reasonably well-detected across the top-100 GFs (net detection score 12, (Fig. 4)). The only other internal WGM that is neither very ancient nor part of a set of WGMs on the same branch, the Solanaceae WGT, achieves a marginally negative net detection score of −4 (Fig. 4). Interestingly, the Brassiceae WGT, the only other WGT in the tree except for the ancient *γ* WGT at the base of the eudicots, also features suboptimal detection: while it is detected by 47 top-100 GFs, 51 other top-100 GFs produce conflicting cLRT results (Fig. 3 and Suppl. Data S2B). This suggests that WGTs are more problematic to detect than WGDs using our BD model, possibly because dosage balance effects may be less pronounced after WGT than after WGD (loosing one of three gene copies produces less dosage imbalance than loosing one of two gene copies). Hence, *λ* values that are primarily trained on the (more abundant) WGDs in the tree may be less appropriate for the (less abundant) WGTs. To assess whether increased gene loss after WGT may explain the decreased detection of WGTs in the 37-species tree, we ran cLRTs for the Brassiceae, Solanaceae and *γ* WGTs separately, under the assumption that they were WGDs instead (i.e. the gene count at a given WGT node was doubled instead of tripled in the BD model). However, the corresponding WGD detection scores further decreased (Suppl. Data S2E-G), suggesting that replacement of WGTs by WGDs in our BD model may overcompensate for increased gene loss after WGT.

### 2.2 Rejection of false WGMs in the 37-species angiosperm phylogeny

In addition to testing the power of the top-100 and bottom-50 RR GFs to detect the 20 well-supported WGMs in the 37-species phylogeny, we also tested the power of these GFs to reject 10 ‘false’ WGMs inserted at positions in the tree where no WGMs are known to have occurred (Fig 1, Suppl. Tab. S2 and Suppl. Data S2H-I) (with exception of the ‘false’ WGD in the Cucurbitales lineage that is in fact likely real, see below). LRTs were performed in the same way as for true WGMs, except that in this case the 20 true WGMs are always considered present and *λ* and likelihood estimations occur in the presence or absence of a single false WGM at a time.

As for the true WGMs, the top-100 GFs are much more decisive than the bottom-50 GFs on the presence or absence of a false WGM (Fig. 5). However, whereas many top-100 GFs rejected one or more true WGMs (Fig. 2), only one of the top-100 GFs detects false WGMs: GF7 (ORTHO001651), which rejected 13 of the 20 real WGMs, also detects all 10 false WGMs. Interestingly, two ‘false’ WGMs are rejected less frequently than the others by the top-100 GFs, although they are not detected either (except by GF7): a WGD in the *Carica papaya* lineage (33 rejections and 66 undecided gene families, Suppl. Data S2H) and a WGD in the Cucurbitales lineage (51 rejections and 48 undecided gene families). There is no support in literature for a WGD in the *C. papaya* lineage, other than the *γ* event shared by all core eudicots (see also Suppl. Fig. S2). On the other hand, several recent studies did find evidence for a Cucurbitaceae-specific WGD (Guo *et al*., 2020; Ma *et al*., 2022), with one study generalizing it to a Cucurbitales-specific WGD (Wang *et al*., 2022a). A substitution rate-adjusted *K*_S_ distribution analysis using *ksrates* (Sensalari *et al*., 2021) supports the Cucurbitales-specific scenario (Suppl. Fig. S3). The less pronounced peak signature in the Cucurbitaceae species from the 37-species tree (cucumber, melon and watermelon), compared to the stronger signal present in *Datisca glomerata* from the Cucurbitaceae sister family Datiscaceae, suggests that there may have been more duplicate gene loss in the Cucurbitaceae, which may help explain why this WGM is not detected in our cLRT analyses.

**Figure 5:**
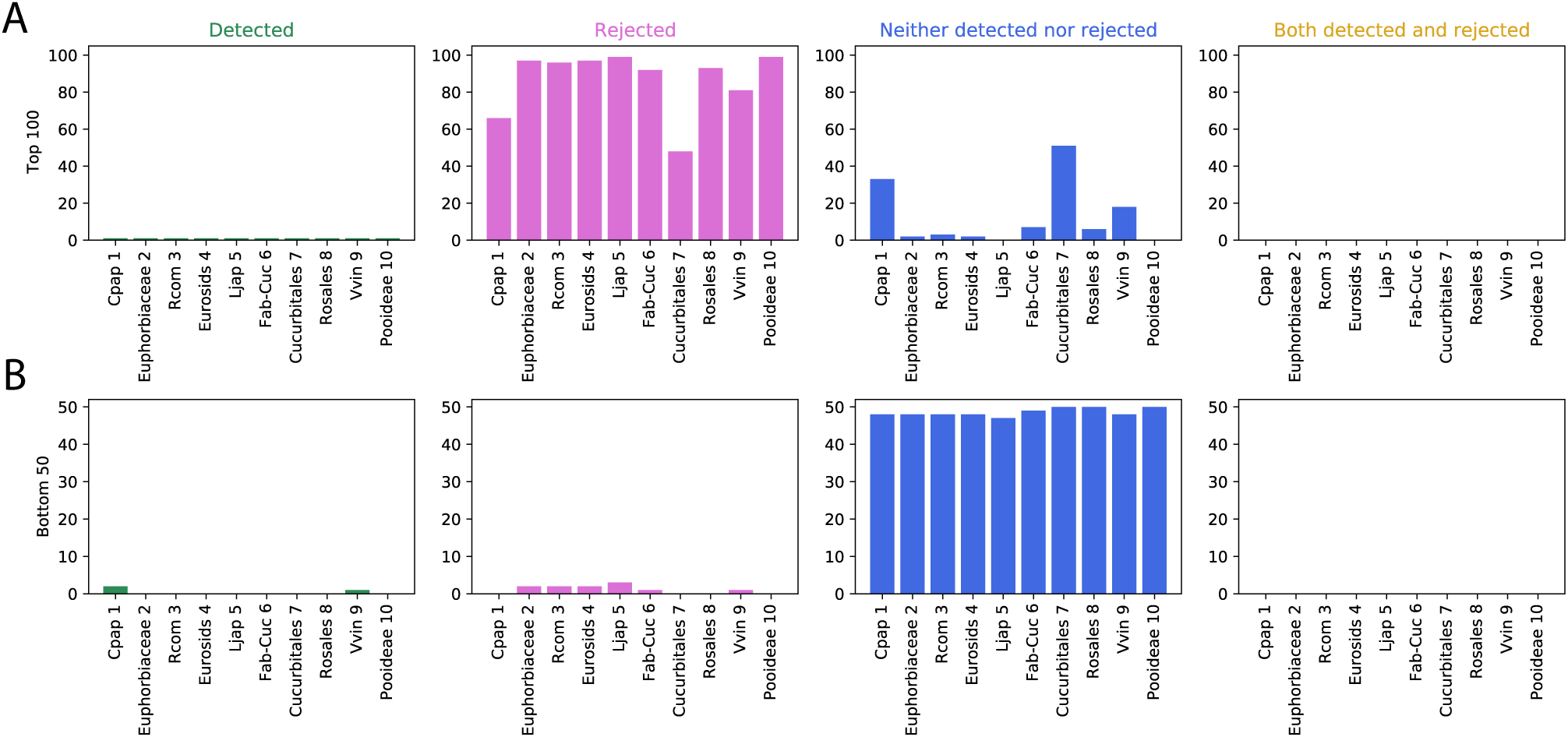
Inference performance on the 10 unsupported WGMs of the top 100 gene families (A) and the bottom 50 gene families (B). The false WGM names on the x-axis are abbreviated, full names reflecting the names of the branches on which the WGMs were modeled can be found in Suppl. Tab. S2. See also Suppl. Data S2H-I.

If the Cucurbitales WGD is indeed a true WGD, as the available evidence indicates, this means there was one true WGD missing in the 37-species phylogeny originally used to rank the GFs in terms of reciprocal retention strength (Tasdighian *et al*., 2017). Although tests by Tasdighian *et al*. (2017) suggested that the *λ*-based GF ranking is fairly robust against removal of a single WGM, the top-100 RR GF set we use here may still deviate from the actual 100 most strongly retained GFs. In order to assess the impact of the Cucurbitales WGD on the *λ* ranking, we re-ranked all 9178 GFs based on the *λ* estimates obtained under the assumption that the Cucurbitales WGD is real (Suppl. Data S1B). The original and reassessed rankings feature a Spearman rank-order correlation coefficient of 0.994 (*p <* 1E-307), with 72 GFs overlapping between the two top-100 lists. These results indicate that leaving out the Cucurbitales WGD does not have a major impact on the inferred reciprocal retention strength of GFs. Given the computational cost of running cLRTs on thousands of GFs, we therefore did not rerun our analyses including the Cucurbitales WGD as a true WGD.

### 2.3 Identification of best-performing gene families

#### 2.3.1 Best-performing GFs in the top-100

16 of the top-100 GFs were found to detect at least 8 out of 20 true WGMs, while rejecting maximum 1 true WGM and producing conflicting cLRT results for maximum 2 WGMs (Suppl. Data S2J). All of these GFs are ranked in the top 50, and the four GFs that detect 15 to 19 WGMs and reject none are all part of the top 5. Next to ‘easy’ tip-branch WGMs that are detected by many of the top-100 GFs, these 16 GFs frequently also detect one or more of the older or repeated WGMs that are more challenging to detect (see full cLRT results in Tasdighian *et al*. (2025)). The eudicot *γ* WGT was detected by 3 of the 16 GFs (ORTHO000593 1, ORTHO002539 and ORTHO000493), the *α*-*β* WGD sequence in the Brassicaceae was detected by 4/16 GFs (ORTHO002539, ORTHO000273 1, ORTHO002657 and ORTHO000887 1), the sequence of three Musa WGDs was detected by 6/16 GFs (ORTHO000593 1, ORTHO002539, ORTHO000273 1, ORTHO001948, ORTHO000493 and ORTHO002651), and the *ρ*-*σ* WGD sequence in the grasses was detected by 6/16 GFs (ORTHO000593 1, ORTHO002539, ORTHO000273 1, ORTHO001948, ORTHO002657 and ORTHO000887 1). The only true WGM not detected by the top-scoring GF (ORTHO002539) is the *τ* WGD in the monocot lineage.

#### 2.3.2 Selection of well-performing GFs beyond the top-100

Due to the substantial computational resources needed for the simulation of distributions of LRT scores under both null hypotheses for the cLRTs (see Methods), we initially focused on the top-100 GFs in the list of GFs ranked by increasing birth-death parameter *λ*, under the rationale that highly RR GFs (low *λ*) should be better WGM markers. Indeed, in the top-100 GFs, *λ* was found to correlate negatively with the number of true WGMs detected (Pearson’s *r* = −0.578, two-tailed *p* = 3.16E-10 (Suppl. Fig. S4A).

Another potential proxy for good WGM marker performance is a strongly negative likelihood ratio (LR) under the null hypothesis that a real WGM is present. Although this LR value in itself doesn’t tell whether or not *H*_0_ is statistically rejected, it can still act as a raw indicator of power and it is far more easily computed than the corresponding p-value (because no background distribution of LR under *H*_0_ needs to be simulated). For the top-100 GFs, the average LR across the 20 real WGMs (with *H*_0_ = *WGM presence*) is more strongly negatively correlated with the number of detected WGMs than the *λ* estimate (Pearson’s *r* = −0.726, two-tailed *p* = 1.27E-17, Suppl. Fig. S4B). We therefore used this average LR as a proxy to scout potentially promising GFs beyond the top-100 GFs, up to rank 2000. We estimated an average LR upper threshold for promising higher-ranked GFs by linearly regressing the average LR for the top-100 GFs against the number of detected WGMs and selecting the expected average LR when 8/20 true WGMs are detected. This resulted in an average LR upper threshold of −0.3472 (Suppl. Fig. S4B). For 13 GFs beyond the top-100 with an average LR ≤ −0.3472 (Suppl. Fig. S5), full cLRT simulations were performed as for the top-100 and bottom-50 GFs (Suppl. Data S2K-L). Five out of these 13 GFs, with ranks 112, 128, 440, 1027 and 1794, detected at least 8 true WGMs (maximum 9) and rarely rejected true WGMs or yielded conflicting outcomes (Suppl. Data S2K). Their detection power appears however to be limited to recent WGMs (Suppl. Data S2L).

### 2.4 Application on independent datasets

The results presented thus far on the WGMs in the 37-species tree of Fig. 1 suggest that strongly RR GFs have higher power to detect WGMs than non-RR GFs. However, it is not guaranteed that the top-performing GFs identified on this 37-species tree will also perform well on plant clades that are not represented in this tree. Indeed, although the 37-species tree covers a wide range of angiosperms and the *λ* values inferred for the top-RR GFs using the 37-species data have a low standard error (see Suppl. Methods), plant clades not represented in the tree may still exhibit different birth-death dynamics (and hence reciprocal retention characteristics) for a given GF than the clades in the 37-species tree.

To assess the broader applicability of prospective marker GFs identified on the 37-species tree, we assessed the performance of the top-100 GFs and the five lower-ranked GFs that detect ≥ 8 true WGMs in the 37-species dataset on three angiosperm clades not included in this dataset and featuring independent, well-supported WGMs, namely the Fagales, Lamiales and Campanulids. To this end, new orthogroups were constructed from the original 37 species’ genomes supplemented by each of these clades in turn, and the resulting orthogroups were matched to the original orthogroups to identify versions of the prospective marker GFs incorporating genes from the extra clade (see Methods andSuppl. Data S3). Note that due to the inclusion of extra species (and possibly also the use of DIAMOND as a homology search algorithm rather than BLAST as used originally on the 37-species dataset), these GFs often do not have exactly the same gene content for the 37 species as the original GFs. After having reconstructed the top-100 GFs and the five additional lower-ranked GFs, we found that the expanded GFs are missing some genes that were originally included and/or possess some extra genes of the original 37 species. For Campanulids (and Fagales, Lamiales, respectively), we found that 12.38% (11.43%, 12.38%) of the reconstructed GFs aren’t missing any genes for the original 37 species, while 41.90% (42.86%, 43.81%) miss up to 5% of the original genes and 35.24% (36.19%, 33.33%) miss between 5% and 10% of the original genes; 10.48% (9.52%, 10.48%) of the reconstructed GFs miss more than 10% of the original genes (Suppl. Fig. S6). 56.19% (53.33%, 52.38%) of the reconstructed GFs do not contain any extra genes for the original 37 species, while 26.67% (25.71%, 29.52%) contain up to 5% extra genes compared to the size of the original GF and 9.52% (10.48%, 9.52%) contain between 5% and 10% extra genes; 7.62% (10.48% and 8.57%) of the reconstructed GFs contain more than 10% extra 37-species genes.

cLRTs were performed for each of the WGMs in each of the independent clades using as input the clade’s inferred gene count profiles for the top-100 GFs plus the 5 selected lower-ranked GFs and a phylogenetic tree including next to the clade’s species also *Amborella trichopoda* and either *Elaeis guineensis* (for Campanulids and Fagales) or *Phoenix dactylifera* (for Lamiales) as outgroups (omitting the other species in the 37-species tree), with the WGMs of interest positioned on the branches (see Methods). The WGM inference results on the new clades are generally poor (see Fig. S7 and full cLRT results in Tasdighian *et al*. (2025)). Only one of the WGMs in the independent clades, the WGD in the Juglandaceae lineage of the Fagales clade, was detected more often than rejected (8 GFs vs. 6 GFs) by the top-100+5 GFs. In the Lamiales clade, the Oleaceae WGT and the Oleeae WGD were each detected by only one of the top-100+5 GFs, while the Pedaliaceae-Phrymaceae WGD was not detected by any GF. In the Campanulids analysis, none of the clade-specific WGMs was detected by any of the top-100+5 GFs. In all three clade analyses, the vast majority of cLRTs was uninformative.

There are a number of potential reasons why WGM inference on the independent clades using the top-100+5 RR GFs inferred from the 37-species dataset may not work well. First, in both the Campanulids and the Lamiales clades, the WGM density is rather high, with 4 and 3 WGMs included in the 4-species Campanulids and Lamiales clades, respectively. In particular the occurrence of two WGMs on the same branch (Apioideae) in the Campanulids clade likely affects inference performance, as in the 37-species dataset, but the occurrence of several other WGMs in close succession may also be problematic (WGD in the Helianthus lineage after WGT in the Asteraceae lineage, and WGD in the Oleeae lineage after WGT in the Oleaceae lineage). Additionally, three out of the seven WGMs in the Campanulids and the Lamiales clades are WGTs, which proved to be more difficult to detect also in the 37-species dataset. On the other hand, this does not explain the poor performance of the top-100+5 GFs in the Lamiales to detect the WGD in the most recent common ancestor of *Erythrante guttata* and *Sesamum indicum* (Pedaliaceae-Phrymaceae WGD), or the borderline detection of the Juglandaceae WGD in the Fagales clade.

Another possibility is that the independent clades are too small to allow accurate *λ* inference and put too few restrictions on *λ* estimates, allowing these estimates to easily accommodate any simulated WGM loss, resulting in a loss of power (Tasdighian *et al*., 2017). We investigated this by incorporating the independent clades in the 37-species tree and running the cLRT pipeline on the extended trees for the 21 best potential marker GFs (16 GFs from the top-100 and the five lower-ranked GFs that detected ≥ 8 true WGMs in the 37-species dataset). Incorporating the clades in a larger tree should have a restrictive effect on the flexibility of *λ* estimates and increase power. Indeed, all WGMs in the added clades were detected by at least some marker GFs in this analysis (Fig. 6 and Suppl. Fig. S8, full cLRT results in Tasdighian *et al*. (2025)), and the Juglandaceae, Helianthus and Pedaliaceae-Phrymaceae WGDs reach net positive detection scores among the 21 marker GFs. The WGTs and the repeated WGMs on the Apioideae branch still have negative net detection scores however, as does the Oleeae WGD. Including any of the independent clades in the tree also affects the performance of the marker GFs on the 20 WGMs included in the original 37-species dataset (Fig. 6).

**Figure 6:**
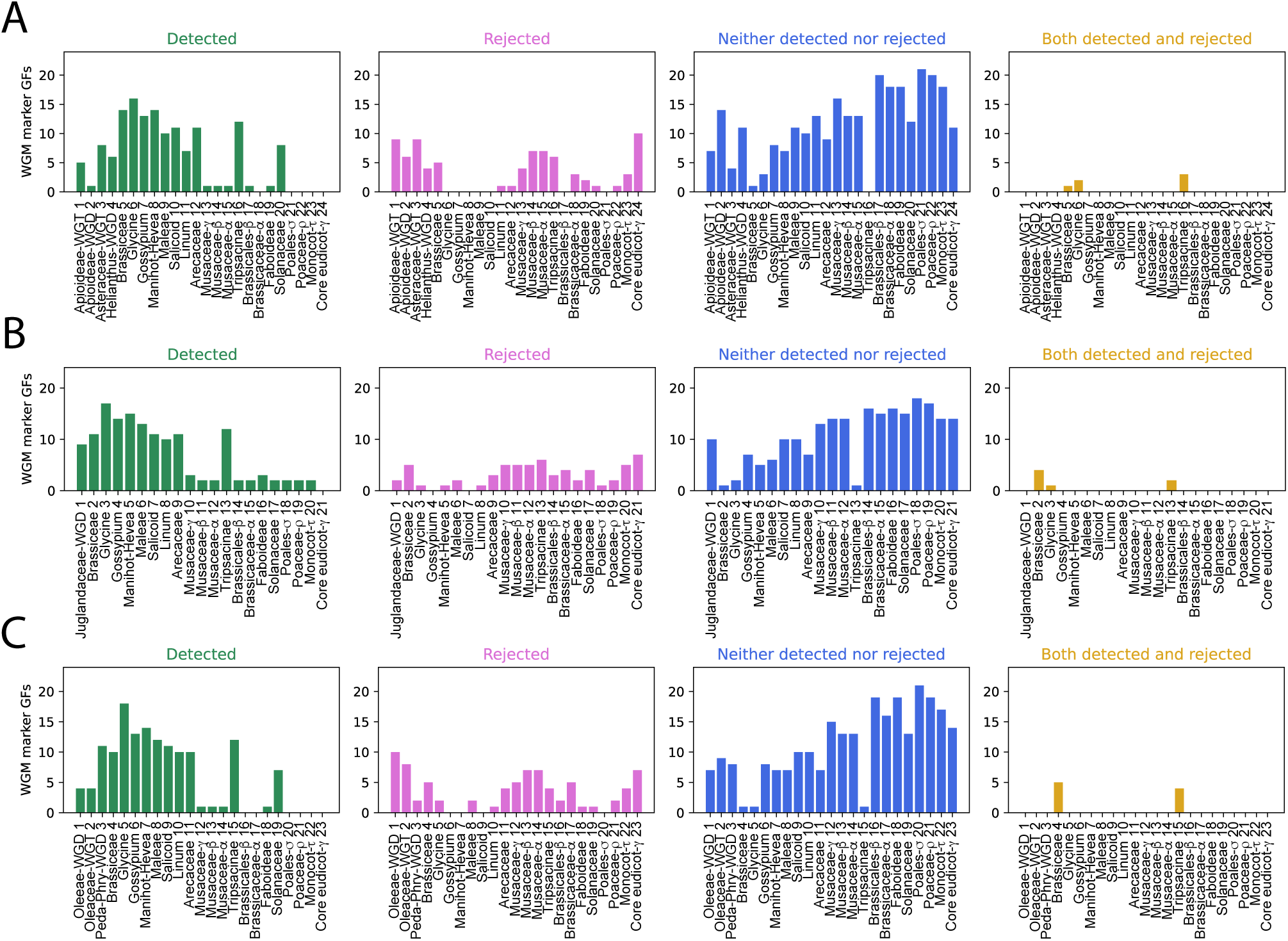
Per-WGM inference performance of the 21 marker GFs on the three extended datasets, encompassing the original 37 species plus Campanulids (A), Fagales (B) or Lamiales (C).

Although three out of seven WGMs in the independent clades, and three out of four WGDs without other WGMs on the same branch, reach net positive detection scores on the extended trees when using the 21 potential WGM markers inferred from the 37-species dataset, their net detection scores do not reach the level of the best-detected WGMs in the 37-species tree (Fig. 6). This suggests that the best marker GFs for the 37-species tree may not necessarily be the best marker GFs for other plant clades. To assess this, we re-ranked the 9178 core-angiosperm gene families according to the *λ* estimates derived from the Fagales dataset (Suppl. Data S1C) and evaluated the performance of the top-100 GFs in this new ranking to infer the WGMs in the Fagales clade. The same procedure was followed for the Lamiales clade (Suppl. Data S1D). The resulting WGM inferences (Suppl. Fig. S9) are indeed better than those made using the original top-100 GFs (Suppl. Fig. 346 S7), with substantial detection scores and low rejection scores for all WGMs despite the low power caused by the small size of the input trees. Spearman’s rank correlations between the Fagales and Lamiales-based GF rankings and the original 37-species-based GF ranking are 0.0763 (*p* =2.489E-13) and 0.169 (*p* =9.408E-60), respectively, and only 5 GFs overlap between the old and new top-100 sets in both cases. These observations indicate that the reciprocal retention-based GF rankings are substantially different across clades (although still significantly correlated). As the reciprocal retention characteristics and WGM inference power of specific GFs vary across clades, no single GF can be considered a good marker GF for all WGMs in all plant lineages.

In conclusion, we have shown that strongly RR GFs have more power to detect true WGMs than non-RR GFs, but the power of any given RR GF is dependent on the plant clade investigated and on the context (other clades in the input dataset). No single RR GF can detect all WGMs under all circumstances. Using a set of marker GFs instead of a single GF and running analyses on a large-enough phylogeny increases WGM inference power in clades not used to select the marker GFs, but small sets of marker GFs selected on one dataset are generally still suboptimal to detect WGMs in other datasets. Furthermore, whereas recent WGDs without other WGMs nearby are generally well-detectable, WGTs, repeated WGMs on the same branch and very old WGMs remain more difficult to detect by our BD model, even on the phylogeny used to select the marker GFs.

### 2.5 Use of reciprocal retention with *WGDgc*

The results described above show that the use of a single RR GF or a small set of RR GFs as WGM markers in our cLRT setup leads to suboptimal WGM inference results, in particular on clades not used to select the marker GFs. As an alternative, we assessed the power of larger sets of RR GFs to detect WGMs. As computationally costly cLRTs need to be performed per GF in our BD model, we instead used *WGDgc*, a previously developed BD framework for WGM detection (Rabier *et al*., 2014), to test the power of larger sets of RR GFs. *WGDgc* is a gene count-based BD model similar to ours, but designed to be run on large sets of GFs at a time and incorporating next to separate gene birth and death rate parameters (as opposed to a single birth-death rate parameter *λ* in our model) also an immediate WGM duplicate retention rate parameter *q*. The latter parameter facilitates a computationally much less costly LRT strategy for WGM testing (Rabier *et al*., 2014). Tiley *et al*. (2016) previously used *WGDgc* for WGM inference on a land plant dataset incorporating 19 WGMs, of which 14 WGMs overlap with our dataset. Using 7564 conserved gene families, they detected 9 out of 19 WGMs, including 7 of the 14 WGMs in common with our dataset: the ‘salicoid’ and Faboideae WGDs, the Glycine WGD, the oldest WGD in the Musaceae, the *σ* WGD in the Poales, the *β* WGD in the Brassicales, and the Solanaceae WGT). The seven other WGMs represented in our dataset (the *ρ* WGD in the Poaceae, the youngest and middle Musaceae WGDs, the Arecaceae WGD, the *τ* WGD in the monocots, the *α* WGD in the Brassicaceae and the *γ* WGT in the core eudicots) were not detected by Tiley *et al*. (2016) when using the full land plant dataset, although four of them (*ρ*, *τ*, Brassicaceae-*α* and *γ*) could be detected in follow-up analyses on smaller 4-taxon trees.

Executing *WGDgc* on our 37-species dataset using the gene counts of the top-100 GFs (original ranking obtained on 37-species tree) detected 14 out of the 20 WGMs at FDR=0.01 (see Tab. 1, Suppl. Data S4 and full WGDgc results in Tasdighian *et al*. (2025)). The *ρ*, *σ* and *τ* WGDs, the Brassicaceae-*α* and Brassicales-*β* WGDs and the Arecaceae WGD were not detected. Interestingly, more true WGMs were detected in the 37-species dataset by *WGDgc* (14/20) at FDR=0.01 than when using the majority vote cLRT strategy on the top-100 GFs at FDR=0.1 (12/20 WGMs detected). Indeed, next to the *ρ*, *τ* and Brassicaceae-*α* WGMs undetected by both methods, the cLRTs also yielded net negative detection scores for the *γ* and Solanaceae WGTs and all three WGDs in the Musaceae, while they detected the Brassicales-*β* WGD that was not detected by *WGDgc*. Given that the same data is used by both methods, these WGM detection differences are due to differences in methodology, such as the inclusion of additional BD parameters in *WGDgc* (see Methods), the use of a single LRT test across all GFs in *WGDgc* rather than complementary LRTs per GF, and concomitant differences in multiple testing correction.

**Table 1:**
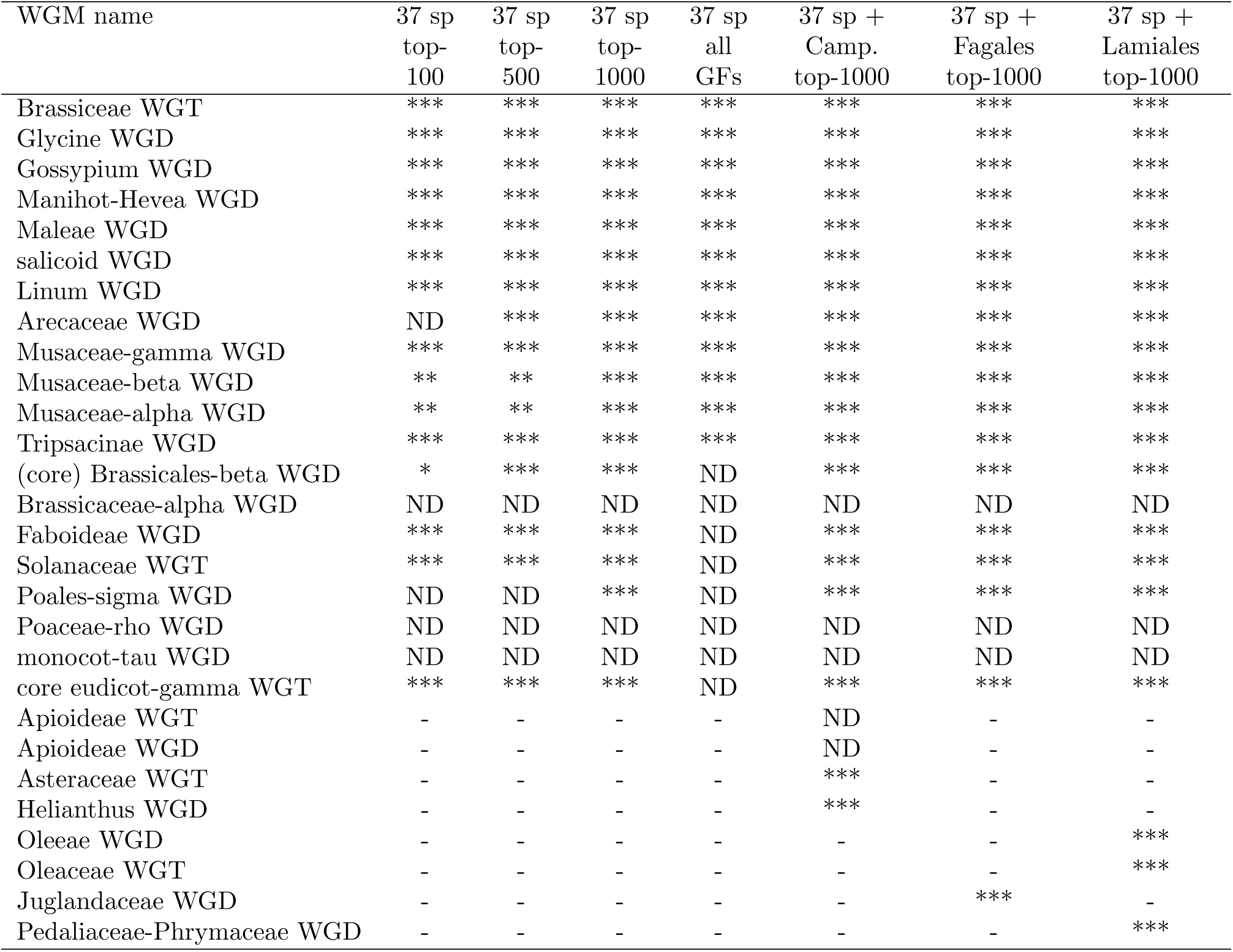
Summary of *WGDgc* WGM inference results. Each row and column summarizes the results for a given WGM and *WGDgc* analysis, respectively. Cell values give the lowest FDR threshold at which a WGM is detected in a given analysis. Full results are gven in Data S4A-U. ND: not detected at FDR=0.05; *: detected at FDR=0.05; **: detected at FDR=0.01; ***: detected at FDR=0.001.

We subsequently ran *WGDgc* on the 37-species dataset using larger sets of RR GFs as input (Tab. 1 and Suppl. Data S4). Using the top-500 RR GFs, *WGDgc* was able to detect 16 out of 20 WGMs at FDR=0.01, with only *ρ*, *σ*, *τ* and the Brassicaceae-*α* WGD escaping detection. Using the top-1000 RR GFs, *WGDgc* detected 17 out of the 20 WGMs at FDR=0.01, leaving only *ρ*, *τ* and the Brassicaceae-*α* WGD undetected. Using larger sets of RR GFs hence improves the WGM detection performance of *WGDgc*. However, when using *WGDgc* on the complete set of 9178 GFs, only 12 out of 20 WGMs are detected at FDR=0.01, with very old WGMs (*γ*, *tau*), several instances of multiple WGMs on the same branch (*ρ* and *σ*, Brassicaceae-*α* and Brassicales-*β*) and two more recent WGMs (Solanaceae WGT and Faboideae WGD) escaping detection (Tab. 1). This indicates that the gene count profiles of less reciprocally retained, lower-ranked GFs contain more noise than signal from the WGM inference perspective and hence interfere with correct WGM inference. Lowering the FDR threshold to 0.001 doesn’t affect the results for the complete 9178 GF set and for the top-1000 GFs, but leads to additional non-detection of the youngest two WGDs in the Musaceae for the top-500 and top-100 GFs (Tab. 1).

Next, we performed *WGDgc* analyses on the three extended datasets including, next to the original 37 angiosperms, species from the Fagales, Lamiales or Campanulids (Tab. 1). For these analyses, we used the corresponding extended top-1000 RR GFs with the same ranks as inferred on the 37-species phylogeny. At FDR=0.01 and 0.001, these analyses produced identical inference results for the 20 WGMs in the 37 species tree as the 37-species *WGDgc* analysis using the top-1000 GFs. Additionally, they detected the Juglandaceae WGD in the Fagales, the Asteraceae WGT and the Helianthus WGD in the Campanulids and all three WGMs in the Lamiales. Hence, of all ‘independent’ WGMs in the investigated clades, only the repeated WGMs in the Apioideae lineage of the Campanulids were not detected. These results show that jointly, the top-1000 RR GFs inferred from the 37-species tree are also able to detect the majority of WGM events in ‘independent’ clades not used to obtain the reciprocal retention ranking.

### 2.6 Use of reciprocal retention in *K* _S_ distributions

Next to testing the WGM inference power of RR GFs in *WGDgc*, we also investigated whether using only RR GFs rather than the entire paranome is beneficial when inferring WGMs from *K*_S_ distributions. In this approach, WGMs are identified as peaks in a distribution of the number of synonymous substitutions per synonymous site (*K*_S_) between paralogous gene pairs, which is used as a proxy for duplication age (Lynch and Conery, 2000). Two sets of genes are routinely used to build *K*_S_ distributions: the entire paranome or a set of ‘anchor pairs’, i.e. duplicated genes located in intragenomic syntenic blocks that are likely remnants of larger-scale (e.g. whole-genome or whole-chromosome) duplications. The focus on anchor pairs mitigates the small-scale duplication (SSD) background that is often prominently present in whole-paranome *K*_S_ distributions and may interfere with WGM inference (Sensalari *et al*., 2021; Tiley *et al*., 2018). Here, we assess whether RR GFs, by virtue of being duplicated preferentially through WGM, can be used to the same effect.

We used the *K*_S_ distribution-based WGM inference tool *ksrates* (Sensalari *et al*., 2021) to infer WGMs in three species (one each in the previously analyzed Fagales, Lamiales and Campanulids clades) using the top-2000 RR GFs (to the extent available, see Methods) inferred from the 37 species and mapped to the corresponding clades, as done previously for the clade-specific cLRT tests. The results of WGM inference for *Olea europaea* var. *sylvestris* (wild olive tree) using *ksrates* on the whole paranome, the set of anchor pairs and the top-2000 RR GFs (see Methods) are shown in Fig. 7. A sizeable SSD background component is visible in the whole-paranome exponential-lognormal mixture model. This SSD background component is substantially reduced in the RR GF lognormal mixture model, and absent in the model based on anchor pair clustering. The Oleeae WGD is picked up and correctly positioned in all three analyses at *K*_S_∼ 0.25, with the smallest component width (i.e. smallest positional uncertainty) in the RR GF analysis. The older Oleaceae WGT is picked up in the anchor pair and top-2000 RR GFs analyses at *K*_S_∼ 0.75, with the corresponding mixture component modes overlapping with the range of *K*_S_ estimates of the split between *O. europaea* and *E. guttata*/*S. indicum*. In the whole-paranome analysis, the Oleaceae WGT is not well-resolved, with two broad and shallow components (b and c) peaking around the split between *O. europaea* and *E. guttata*/*S. indicum*. Both literature (Wang *et al*., 2022b) and *K*_S_ distribution analysis (Suppl. Fig. S15) indicate that the WGT occurred in *O. europaea* lineage after its divergence from *E. guttata* and *S. indicum*. The Oleaceae WGT component and the underlying *K*_S_ peak are substantially narrower in the RR GF analysis than in the two other analyses, leading to a more pronounced separation of the Oleeae WGD and the Oleaceae WGT in the RR GF model. The *γ* WGT in the core eudicots is picked up (although very faintly) in the RR GF model, but not in the whole-paranome and anchor pair analyses. Qualitatively similar results are obtained on *Helianthus annuus* (sunflower, Suppl. Fig. S10) and *Corylus avellana* (common hazel, Suppl. Fig. S11). Most WGMs are picked up by all methods (except for the *γ* WGT in the *H. annuus* lineage, which is missing in the anchor pair analysis and misplaced in the RR GF analysis), the SSD background is reduced in the RR GF analysis and nearly absent in the anchor pair analysis, and the WGM component widths are generally smallest in the RR GF analysis, leading to better distinction between nearby WGMs.

**Figure 7:**
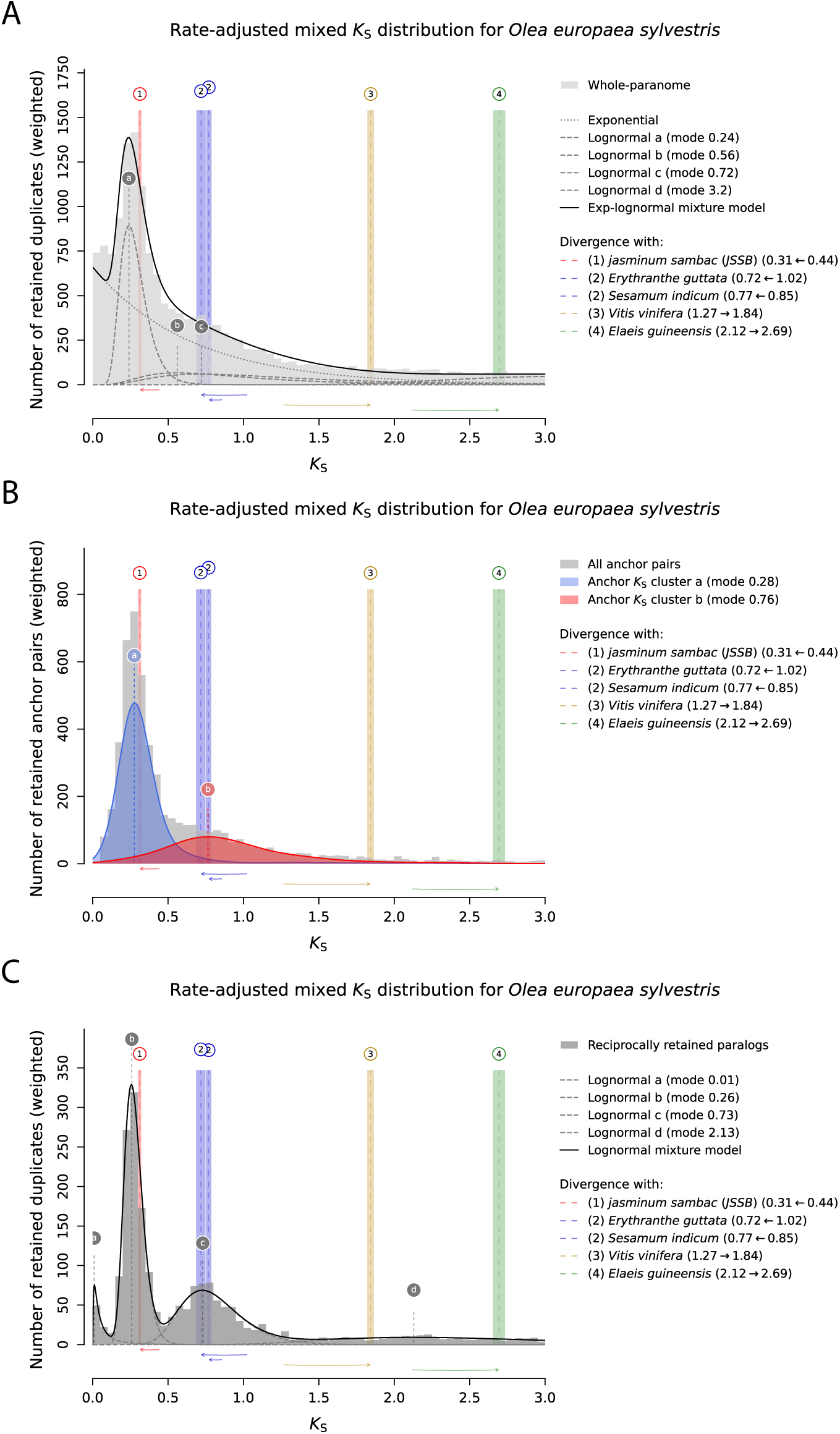
Comparison between whole paranome (A), anchor pair (B) and RR GF (C) *K*_S_ distributions for wild olive tree, generated by *ksrates*. WGM peaks were inferred by means of exponential-lognormal mixture modeling (A), anchor pair clustering (B) and lognormal mixture modeling (C). The modes of these WGM peaks are shown as vertical lines labeled with letters. Vertical lines labeled with numbers indicate the modes of ortholog *K*_S_ distributions, which mark the divergence between wild olive tree and other species. Colored boxes range from one standard deviation (SD) below to one SD above the mean mode estimate. The *K*_S_ ages of ortholog *K*_S_ modes have been adjusted on the basis of substitution rate differences between species, as illustrated by the horizontal arrows above the x-axis.

We additionally evaluated the use of the top-2000 RR GFs in the grass species *Brachypodium distachyon* (Suppl. Fig. S12). Both anchor pairs and RR GFs recover the more recent *ρ* WGD and picked up an older signal possibly covering both the *σ* and *τ* WGDs. The latter signal is more pronounced and better separated from the *ρ* WGD in the RR GF analysis than in the anchor pair analysis. The whole-paranome model recovers *ρ* (component a) and contains components that may correspond to *σ* (component b) and *τ* (component c), but the size (area) of these components seems unrealistically big compared to the size of the more recent *ρ* component, as components are expected to become smaller with age due to duplicate gene loss.

In summary, our results show that using RR GFs in *K*_S_ distribution-based WGM inference is a valid alternative to using whole paranomes or anchor pairs, with better ability to distinguish overlapping WGM peaks than either alternative and potentially more power than anchor pairs for older WGMs. An added advantage of using RR GFs over anchor pairs is that identification of syntenic blocks (which requires a fully assembled genome) is not necessary, so that RR GF analyses may also be performed based on transcriptome data.

## 3 Discussion

Ever since the first eukaryote whole-genome sequences became available (Goffeau *et al*., 1996), researchers have been looking for signals of ancient WGMs in these sequences (Wolfe and Shields, 1997). Various techniques have been developed over the years for this purpose, and newly sequenced genomes (in particular plant genomes) are now routinely analyzed with one or more of these techniques in search of WGM signals. None of the currently available methods is perfect however, and their results are sometimes open to interpretation, causing several WGM claims reported in literature to be disputed (Ruprecht *et al*., 2017; Zwaenepoel *et al*., 2019; Nakatani and McLysaght, 2019). Moreover, different methods sometimes give different results when ran on the same dataset, and all methods are sensitive to the amount and quality of the data analyzed. When using gene tree-species tree reconciliation and gene count-based methods for instance, the taxon sampling density in the input phylogeny may affect the WGM inferences obtained (McKibben *et al*., 2024).

Small-scale duplications are a major source of noise and dubious WGM inferences when using reconciliation-, gene count- and *K*_S_-based methods (Zwaenepoel *et al*., 2019; Sensalari *et al*., 2021; Tiley *et al*., 2018). Synteny-based WGM detection is comparatively immune to such SSD effects by virtue of focusing on syntenic blocks of duplicated genes remaining from (mostly) larger-scale duplication events. Similarly, only using the anchor pairs from a synteny analysis in *K*_S_-based methods, instead of the whole paranome, substantially reduces the impact of SSDs on WGM inference results. In both cases however, high-quality genome assemblies are required to infer syntenic regions, and finding syntenic WGM remnants gets progressively more difficult for older WGMs.

In this study, we tried to eliminate the effect of SSDs on WGM inference in an alternative way, by focusing on strongly RR GFs that preferentially retain duplicates after WGM and rarely duplicate through SSD. We assessed whether these GFs could be used as WGM markers, using a probabilistic framework in which the gene count of a GF of interest is modeled across a species tree as a function of a birth-death parameter *λ*, a GF size at the root of the tree and putative WGMs positioned on the branches. We find that strongly RR GFs have indeed more power to detect true WGMs and reject false WGMs than non-RR GFs. However, while the best-performing GF (GF3) detected 19/20 well-supported WGMs in a 37-species tree, only 12 of the top-100 RR GFs could detect half of these WGMs or more. While recent single WGMs on short terminal branches were generally detected by more than half of the top-100 RR GFs (with the exception of the Brassiceae WGT), single WGMs on longer terminal branches (Linum and Arecaceae WGMs) and longer internal branches (Faboideae and Solanaceae WGMs) were only detected by a minority of GFs. Given that a longer branch gives the model more opportunity to compensate for the omission of a real WGM through increased SSD, it makes sense that detecting WGMs on longer branches is more difficult. Similarly, ancient WGMs such as the *γ* and *τ* events are more challenging to detect because their omission in the model can be compensated for over a longer time scale. Detecting WGMs on branches that also harbor other WGMs (Musaceae, Brassicales and Poales lineages) proved as challenging as detecting ancient events, likely because decreased gene loss after one WGM can compensate for the omission of another one in the model. Tiley *et al*. (2016), using a BD modeling approach similar to ours on gene count data for thousands of plant GFs simultaneously, similarly found that ancient WGMs and repeated WGMs on the same branch were hard to detect. Some of these issues may be remedied through the inclusion of additional taxa in the phylogeny to break up long branches or a series of WGMs on the same branch.

Our analysis of individual RR GFs identified several promising WGM marker GF candidates among the top-100 RR GFs. However, given that the GFs’ reciprocal retention strength was inferred from the same 37-species dataset as used for the WGM inference tests, our selection of 100 GFs to be tested may be biased toward GFs with more power to specifically detect the 20 WGMs in this 37-species dataset rather than angiosperm WGMs in general. In other words, even though the selected GFs can confidently be categorized as strongly reciprocally retained in angiosperms given the taxonomic breadth of the 37-species input phylogeny used (Tasdighian *et al*., 2017), the best-performing GFs on the 37-species dataset are not necessarily the best marker GFs for other WGMs than the ones included in this dataset. When testing the performance of the top-100+5 GFs identified from the 37-species dataset on independent plant clades (Fagales, Lamiales and Campanulids), we found that the candidate marker GFs performed very poorly on detecting WGMs in these clades, and that other GFs exhibited better performance. One potential explanation for the lack of power of the top-100+5 GFs to detect WGMs in smaller independent clades is that their optimal *λ* values are insufficiently constrained (i.e. kept low) by other plant taxa, allowing the model to adapt more flexibly to the presence or absence of a WGM. In support of this hypothesis, the detection performance of candidate marker GFs on the small clades’ WGMs improved when the clades were embedded in the larger 37-species phylogeny. However, in general, the tested GFs were still less performant in detecting newly added WGMs than in detecting the WGMs originally in the 37-species tree. This indicates that the identified candidate marker GFs are indeed biased to some extent toward the original tree, and that no single GF is an optimal WGM marker across all angiosperm clades.

One option to remedy the shortcomings of single candidate marker GFs is to aggregate the results of a number of marker GFs and call WGM presence or absence based on the detection-rejection balance across all markers. However, given that all candidate markers likely have clades in which they perform better or worse one would have to use (a lot) more marker GFs than the 21 identified based on the 37-species phylogeny. As the methodology we developed to model the BD dynamics of individual GFs is very compute-intensive (in particular for generating the background probability distributions for the cLRTs), we used *WGDgc*, a previously developed BD modeling framework, to model multiple GFs simultaneously (Rabier *et al*., 2014; Tiley *et al*., 2016). In comparison to our model, the *WGDgc* model includes extra parameters *q* capturing duplicate retention after a WGM (1-*q* can be considered a rate of near-immediate gene loss after WGM). Incorporation of this parameter *q* with values between 0 and 1 in the model (whereas *q* can only be 0 and 1 in our model, when a WGM is absent or present in the model, respectively) allows likelihood ratio tests to be performed for WGM detection without the need for constructing background LRT distributions, making the *WGDgc* formalism much less compute-intensive. On the other hand, one birth and death rate is estimated for all GFs simultaneously in *WGDgc*, whereas each GF has its own BD rate parameter in our model. However, when modeling GFs with similar BD rates, as should be the case when using a set of RR GFs with low BD rates as input, this should not be a drawback.

Using the top-1000 RR GFs as predictors, *WGDgc* detected all but 3 of the 20 *bona fide* WGMs in the 37-species tree at FDR=0.001. Similarly, all WGMs in the Lamiales, Fagales and Campanulids clades were detected when analyzing extended phylogenies, except for the repeated WGMs in the Apioideae lineage. In contrast, when using all 9178 GFs as predictors, 8 of the 20 WGMs in the 37-species tree were not detected, indicating that inclusion of non-RR GFs adds noise rather than signal for WGM detection. On the other hand, restricting the GF set to the top-500 or top-100 RR GFs led to lower WGM detection performance than when using the top-1000, indicating that the set of RR GF predictors in *WGDgc* may also not be taken too small.

Similarly, using the top-2000 RR GFs rather than the whole paranome to construct *K*_S_ distributions (Fig. 7) yielded *K*_S_ distributions with a lower SSD background and narrower WGM peaks, and hence an improved WGM resolution (in particular for overlapping WGM peaks). The WGM peaks in the RR GF *K*_S_ distributions were even found to be narrower than those in anchor pair *K*_S_ distributions, indicating that anchor pairs belonging to non-RR GFs, by virtue of being less subject to dosage balance effects and the associated sequence divergence constraints (Tasdighian *et al*., 2017), may contribute to peak widening in *K*_S_ distributions.

In summary, our results show that RR GFs carry more WGM signal than non-RR GFs. Although any given RR GF has more power to detect WGMs in some plant clades than in others, and although no single RR GF can detect all WGMs in all clades, a sufficiently large set of RR GFs jointly has more power to detect true WGMs than the entire paranome. We therefore advocate the use of sets of RR GFs in both count-based and *K*_S_ distribution-based tools for WGM inference. To facilitate the latter, a pipeline to retrieve RR GFs in angiosperm clades has been implemented in *ksrates* v2.0 (https://github.com/VIB-PSB/ksrates). Restricting gene tree-species tree reconciliation analyses to RR GFs may similarly improve WGM inference performance, but this was not tested in this study.

## 4 Materials and Methods

### 4.1 Gene count profiles, phylogenetic trees and WGM placement

#### 4.1.1 37-species angiosperm dataset

For identifying potential WGM marker GFs, we initially reused the angiosperm dataset used in Tasdighian *et al*. (2017), including a 37-species phylogenetic tree with 20 well-supported WGMs positioned on the branches (Fig. 1 and Suppl. Tab. S1) and the associated gene count profiles of 9178 core angiosperm gene families from Li *et al*. (2016). For assessing the power of GFs to reject false WGMs, 10 additional WGMs were arbitrarily positioned on branches lacking support for the occurrence of WGMs at the start of this study (Suppl. Tab. S2).

#### 4.1.2 Additional clade datasets

To assess the power of candidate marker GFs retrieved from the 37-species dataset on other plant clades, we compiled data on three angiosperm clades not represented in the 37-species tree, namely the Campanulids, Fagales and Lamiales. Representative species from these clades were selected based on the availability of high-quality sequenced genomes, gene annotations and structural annotation (Suppl. Tab. S3). The number of species in each clade was intentionally kept small (maximum 6 species) in order to reduce the computational resources needed for birth-death modeling and cLRTs (see below). Signatures of WGMs other than those included in the 37-species dataset were previously found in the genomes of several of the selected species (Suppl. Tab. S4).

The presence and phylogenetic positioning of these WGMs was confirmed through *K*_S_ distribution analysis using *ksrates* (Sensalari *et al*., 2021). For any given species in a given clade, *ksrates* was used to compute an anchor pair *K*_S_ distribution of the species, plus the *K*_S_ distributions of ortholog pairs between the species of interest and the other species in the clade of interest, as well as *Elaeis guineensis* and *Vitis vinifera* (these species were included to frame the *K*_S_ range in which the eudicot *γ* WGT occurred). *ksrates* was then used to perform lognormal mixture modeling or clustering of the anchor pair *K*_S_ values, and to display the resulting WGM components together with substitution rate-adjusted mode estimates for the ortholog *K*_S_ distributions, using *Amborella trichopoda* as the outgroup for rate adjustment, in order to assess the phylogenetic positioning of the inferred WGMs (see Suppl. Figs. S13, S14, S15 and S16).

To obtain the phylogenetic tree for a given clade (Suppl. Data S5), we first performed homology searches using DIAMOND v2.0.14 (Buchfink *et al*., 2021) on the translated protein-coding sequences of the clade’s species and selected outgroups (*A. trichopoda* for all three clades, *Elaeis guineensis* for Fagales, and *Phoenix dactylifera* for Campanulids and Lamiales); option *–max-target-seqs* was set to 750 in order to report sufficient homologous pairs. Gene families were defined through orthogroup clustering with OrthoMCL v1.4 (Li *et al*., 2003) using inflation factor 1.5. 100 single-copy GFs were randomly selected from the total number of single-copy GFs available (2100 for Campanulids, 795 for Fagales and 2365 for Lamiales), and an aminoacidic multiple sequence alignment (MSA) was generated for each using MUSCLE v3.8.31 (Edgar, 2004). The MSAs were trimmed and back-translated using trimAl v1.4.rev15 (Capella-Gutiérrez *et al*., 2009) with the *strictplus* and *splitbystopcodon* options. For each clade, the 100 single-copy GF MSAs were concatenated, and the concatenated MSAs (with a length of 125,250 nucleotides for Campanulids, 395,385 for Fagales and 114,141 for Lamiales) were used to build phylogenetic trees with IQ-TREE2 (Minh *et al*., 2020), employing the MGK codon substitution model, codon frequencies estimated using the F3X4 model, four gamma rate categories (G4) to account for rate heterogeneity among sites and ultrafast bootstrap approximation (-B 1000). The resulting trees were rooted on *A. trichopoda* (see Suppl. Data S5 and Suppl. Fig. S17).

WGMs were placed on the clade trees by dating them in terms of the average number of substitutions per codon (t) between anchor pairs (Suppl. Data S5). Species-specific anchor pair t distributions were built with *ksrates* (Sensalari *et al*., 2021) using t values estimated by codeml (PAML v4.9j, Yang (2007)) using default parameters. Subsequently, lognormal mixture models were fit to the t distributions and WGM ages in terms of t were inferred from the corresponding component modes in the *ksrates* model output. WGMs were then positioned on the clade trees by tracing back a distance equal to half the t age along the lineage of the species concerned starting from its leaf node. In cases where multiple species shared a WGM, resulting in different WGM position estimates, the distance of these estimates to the root of the phylogeny was averaged and this average distance was used to position the WGM, as in Tasdighian *et al*. (2017). The consensus WGM positions in the three independent clades are consistent with literature (Suppl. Fig. S17).

For generating the clade gene count profiles of the 9178 core angiosperm gene families for which the WGM marker potential was assessed, we first used DIAMOND v2.0.14 (Buchfink *et al*., 2021) to assess homology relationships among translated protein-coding sequences of both the species of the clade of interest and the original 37 species (using the same genome versions as used in Li *et al*. (2016) for the latter); option *–max-target-seqs* was set to 750 in order to report sufficient homologous pairs. Compared to the approach used in Li *et al*. (2016) for the 37-species dataset, we opted to use DIAMOND instead of BLAST due to its significantly faster run time. The resulting homology table was provided to OrthoMCL v1.4 (Li *et al*., 2003) for orthogroup clustering with inflation factor 1.5. Subsequently, the OrthoMCL orthogroups were mapped to the original 9178 GFs based on overlap in gene content for the original 37 species (note that multiple orthogroups can overlap with one original GF). A filtering step was used to avoid inflating the original gene counts for the 37 species with genes belonging to other angiosperm GFs: if the overlap between a new orthogroup and an original GF included a large number of extra 37-species genes (≥ 15 genes for Campanulids and Fagales and ≥ 16 genes for Lamiales), the match was discarded. Additionally, in a few cases (two in the top-100+5: original GFs ORTHO001338 and ORTHO000847, for all three clades analyzed), a new orthogroup with 14 extra genes was discarded because it overlapped poorly with the original GF (≤ 3 common genes). If multiple new orthogroups mapped to an original GF after filtering, they were merged to create the new GF. For the clade-specific cLRT analyses (see further), only the extended GF counts for the species in the clade of interest were considered (Suppl. Data S3). A small percentage of the 9178 reconstructed GFs didn’t have any gene in the species belonging to specific clade datasets (including outgroups): 179 in the Campanulids clade (2 within the top-100+5 GFs, i.e. ORTHO000580 1 and ORTHO003707), 157 in the Fagales clade (1 within the top-100+5 GFs, i.e. ORTHO000806 1) and 164 in the Lamiales clade (3 within the top-100+5 GFs, i.e. ORTHO002268, ORTHO000806 1 and ORTHO003707).

#### 4.1.3 Extended datasets including the original 37 species and one of the additional clades

For generating the extended datasets (original 37 species plus either Campanulids, Fagales or Lamiales species), the same overall workflow was followed as for the additional clade datasets described in the previous section. To keep the computational cost low, we introduced only three Fagales species in the dataset extended with Fagales (*Carya illinoinensis*, *Corylus avellana* and *Myrica rubra*), instead of the six used in the independent clade analysis. A small percentage of the reconstructed GFs didn’t have any gene in specific extended datasets: 105 in the Campanulids-extended dataset (1 within the top-100+5), 90 in the Fagales-extended dataset (none within the top-100+5) and 93 in the Lamiales-extended dataset (1 within the top-100+5), but none of these GFs belonged to the set of 21 potential marker GFs that was tested on the extended datasets.

For generating the phylogenetic tree of an extended dataset incorporating the original 37 species and a given additional clade, the OrthoMCL results used for generating the GF count profiles of the clade concerned (see previous section) were reused to identify single-copy gene families in the corresponding extended species set. However, only 2, 5 and 4 orthogroups were found to be strictly single-copy for the 37-species set extended with Campanulids, Fagales and Lamiales species, respectively. We therefore searched for orthogroups with at least one member gene per species, but allowing up to 3 extra paralogs. We obtained 96, 116 and 98 such orthogroups for the extended species sets when including Campanulids, Fagales and Lamiales, respectively. These orthogroups were then pruned to single-copy orthogroups by retaining for each species only the paralog with the most orthologous hits in the orthogroup. Based on these pseudo-single-copy orthogroups, phylogenetic trees were constructed for each of the extended species sets as described above for the additional clade datasets (Suppl. Data S5). The resulting extended trees featured the same topology for the original 37 species as the original 37-species tree; Campanulids and Lamiales were placed as sister groups to the Solanaceae in the extended trees (*Solanum tuberosum* and *Solanumlycopersicum*) (Suppl. Fig. S18 and S19), in agreement with the AGP IV classification (Stevens, 2017), while Fagales was placed as sister to the Rosaceae (Suppl. Fig. S20), although AGP IV currently supports a sister relationship with Cucurbitales.

The placement of the Fagales, Campanulids or Lamiales WGMs on the extended tree followed the same approach described earlier for the additional clade datasets (see previous section and Suppl. Data S5); for the placement of the 20 WGMs of the original dataset, we reused the averaged t ages of the WGMs as computed by Tasdighian *et al*. (2017), in order to keep the WGM positioning as similar as possible (see Suppl. Methods 1.2).

Due to small branch length differences between the extended trees and the original 37 species tree, some WGMs were positioned on the wrong branch in the extended trees when using the averaged t ages of Tasdighian *et al*. (2017). The Brassicales-*β* WGD was erroneously positioned before the split between the Brassicaceae and *C. papaya*, and was moved to the beginning of the Brassicaceae branch. The Juglandaceae WGD in the extended tree with Fagales was positioned just before the split between *C. illinoinensis* and *M. rubra*, rather than being specific to Juglandaceae, and hence was moved to the base of *C. illinoinensis* branch. Similarly, the *Musaceae*-*γ* WGD was wrongly positioned just before the split between *M. acuminata* and *P. dactyliphera*, and was moved to the base of *M. acuminata* branch. The *Z. mays* WGD was wrongly positioned just before the split between *Z. mays* and *Sorghum bicolor*, and was moved to the base of *Z. mays* branch. Finally, the *γ* WGT was wrongly positioned as shared only among eurosids (i.e. excluding *V. vinifera*), and was moved to the eudicot stem branch.

The gene family counts used for the extended datasets were the same as the counts obtained for the corresponding additional clade datasets (see previous section), but this time retaining also the counts for the original 37 species (Suppl. Data S3).

### 4.2 WGM inference through complementary Likelihood Ratio Tests

#### 4.2.1 Birth-death model of gene family size evolution

The birth-death (BD) model described in Tasdighian *et al*. (2017) was used to model the evolution of gene family sizes across a phylogenetic tree. Briefly, given a species tree, a set of WGMs positioned on the tree and the gene counts of a gene family of interest at the tips of the tree, this BD model computes the log-likelihood of the observed gene count profile given a unified birth-death rate parameter *λ* and the gene family size *r* at the root of the phylogeny. This log-likelihood is maximized over *λ* and *r* through cutting-plane optimization (Marchand *et al*., 2002). In Tasdighian *et al*. (2017), mainly the maximum likelihood estimate of *λ* (*λ*\*) was of interest as a measure for the reciprocal retention strength of the gene family being studied. Here, the quantity of interest is rather the maximum log-likelihood itself (*logLk*\*), as a measure of how well a given arrangement of WGMs on the tree is supported by the observed gene count profiles.

#### 4.2.2 Complementary likelihood ratio tests

For each gene family *j* and for each true WGM *i*, we calculated the maximum log-likelihood scores of the observed gene count profile under two models: a model in which all true WGMs, including WGM *i*, are present, and a model in which WGM *i* is absent. The resulting log-likelihood scores were used in likelihood ratio tests (LRTs). However, as the two models being compared are not nested, it is impossible to establish which one should be taken as the null hypothesis (*H*_0_) for the LRT. This matters because for non-nested models, the likelihood ratio under *H*_0_ is not asymptotically chi-squared distributed and its distribution may differ depending on whether the presence or absence of WGM *i* is taken to be *H*_0_, potentially leading to situations in which LRTs using one or the other model as *H*_0_ produce conflicting results (Lewis *et al*., 2011). We therefore ran complementary LRTs (cLRTs) in which each of the competing hypotheses is taken in turn as *H*_0_. The two cLRTs produce four possible outcomes (Tab. 2): the two tests can reach a consensus for detection or rejection of WGM *i* by GF *j*, they can both not reject their *H*_0_ (in which case the test results are indecisive and no presence or absence call can be made for WGM *i* based on GF *j*), or they can both reject their *H*_0_ (conflicting results indicating that neither model is appropriate).

**Table 2:** The four possible outcomes of a complementary likelihood ration test (cLRT).

| LRT<sub>1</sub> | LRT<sub>2</sub> | Inference |
| --- | --- | --- |
| $H_0 = \text{WGM presence}$ | $H_0 = \text{WGM absence}$ | |
| $H_1 = \text{WGM absence}$ | $H_1 = \text{WGM presence}$ | |
| $H_0$ not rejected | $H_0$ rejected | WGM detected by GF |
| $H_0$ rejected | $H_0$ not rejected | WGM rejected by GF |
| $H_0$ not rejected | $H_0$ not rejected | Undecided GF |
| $H_0$ rejected | $H_0$ rejected | Conflicting GF |

For each of the two cLRTs, the likelihood ratio test statistic for the observed gene count profile (“observed LR”) is calculated as in Eq. 1.

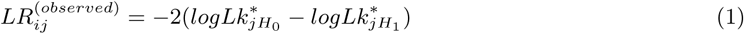

with *H*_0_ and *H*_1_ being the presence and absence of WGM *i*, respectively, in one of the cLRTs, and vice versa in the other cLRT. Given the non-nested nature of the models tested, a standard closed-form solution for the distribution of LR under *H*_0_ that can be used to analytically compute *p*-values for both cLRTs is not available (unlike for the BD models used in Rabier *et al*. (2014) and Tiley *et al*. (2016)). Instead, as proposed in Williams (1970), we approximated the background distributions using model simulations under *H*_0_ (taking each of the competing hypotheses as *H*_0_ in turn).

For any given gene family *j* and WGM *i*, 1000 gene count profiles *ζ* were generated from either *H*_0_ model through Markov Chain Monte Carlo (MCMC), using the optimal 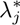 and 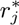 obtained from the *H*_0_ BD model for the observed gene count profile of GF *j*. The BD model was then fit to each simulated profile *ζ* under *H*_0_ and *H*_1_, resulting in two maximum log-likelihoods 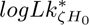 and 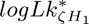 and their LR statistic (“simulated LR”) (Eq. 2):

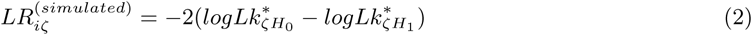

Together, the 1000 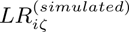 for a given *H*_0_ approximate the LR distribution under this *H*_0_. For each of the competing *H*_0_, an approximate p-value for the observed gene count profile *j* being generated under *H*_0_ was then computed as (Ojala and Garriga, 2010; Lewis *et al*., 2011):

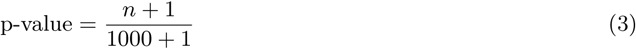

with *n* the fraction of the 1000 simulated LRs that is lower than the observed LR score. This procedure was repeated for every true WGM *i* and GF *j*. For every GF *j*, the list of *p*-values obtained for all WGMs under a given null hypothesis was corrected for multiple testing using the Benjamini-Hochberg correction (Benjamini and Hochberg, 1995). A given *H*_0_ for a given GF and WGM was rejected when its FDR-adjusted p-value was smaller than 0.1.

The workflow described above for true WGMs was also applied for false WGMs. Here, the models compared were a model in which all true WGMs and the false WGM of interest are present, and a model in which the true WGMs are present and the false WGM is absent. Model code is available at https://github.com/VIB-PSB/BirthDeathModel.

### 4.3 WGM inference using *WGDgc*

To test the power of large sets of RR GFs at a time, we used the alternative gene count-based BD model implemented in *WGDgc* v1.3 Rabier *et al*. (2014) in R version 4.1.3. *WGDgc* analyses were performed on the 37-species dataset for all 9178 GFs and for the top-100, top-500 and top-1000 RR GFs. *WGDgc* analyses on the extended datasets were performed for the top-1000 RR GFs only. A small percentage of the top-1000 reconstructed GFs used for the *WGDgc* analyses didn’t have any gene in the species belonging to the clade dataset (including outgroups) and were hence omitted: 18 in the Campanulids-extended dataset, 13 in the Fagales-extended dataset and 11 in the Lamiales-extended dataset. For each dataset, we ran command *MLEGeneCount* once including all WGMs in the tree (*H*_1_) and subsequently multiple times removing each single WGM at a time (*H*_0_); *geomMean* was always set to default 1.5 and, when analyzing all 9178 GFs, *mMax* (the maximum number of surviving lineages at the root) was set to 33, i.e. the size of the largest gene count, to reduce the computational requirements. After calculating the likelihood ratio (LR) between the two fitted models, we retrieved the p-value assuming that the asymptotic distribution for the LR statistic is a 50:50 mixture of a point mass at *q* = 0 and a *χ*-square distribution with 1 degree of freedom (DOF) as described in Rabier *et al*. (2014). We finally performed multiple testing correction through the Benjamini-Hochberg procedure at various thresholds (0.05, 0.01 and 0.001).

### 4.4 WGM inference using *K*_S_ distributions

Whole-paranome and anchor pair *K*_S_ distribution generation for species of interest and subsequent WGM inference on these distributions was done using *ksrates* v1.1.4 (Sensalari *et al*., 2021). *K*_S_ distributions of RR GFs were constructed and analyzed using *ksrates* v2.0 (https://github.com/VIB-PSB/ksrates). The construction of RR GF *K*_S_ distributions in *ksrates* v2.0 follows the same steps for the generation of gene count profiles as outlined above for the extended datasets. Adaptations include 1) extending the original 37 angiosperm species by adding only a single species of interest, and 2) retaining only the genes of this species of interest for *K*_S_ estimation after GF reconstruction. Moreover, the orthogroup clustering step is performed with *OrthoMCLight*, a faster implementation of OrthoMCL v1.4 that optimizes the BLAST table reading step and supports multi-threading, developed to meet the performance requirements of the reciprocal retention pipeline (https://github.com/VIB-PSB/OrthoMCLight). The *ksrates* v2.0 pipeline for WGM inference based on RR GFs is designed to reconstruct the top-2000 RR GFs of the 37-species *λ* ranking, rather than the top-100 as in the cLRTs or the top-1000 as for the *WGDgc* analyses, to ensure sufficient *K*_S_ data while still limiting the introduction of SSD signal from higher-ranked, non-RR GFs. Several factors contribute to *K*_S_ data scarcity: top GFs lacking genes from the species of interest cannot be reconstructed; top GFs containing only a single gene from the species of interest cannot provide *K*_S_ values, since at least a duplicate pair is required; and RR GFs often contain fewer members than the average GF used in whole-paranome *K*_S_ distributions. The effective number of reconstructed top-2000 GFs was 1306 for *O. europaea* var. *sylvestris* (Fig. 7), 1386 for *H. annuus* (Suppl. Fig. S10), 593 for *C. avellana* (Suppl. Fig. S11) and 871 for *B. distachyon* (Suppl. Fig. S12).

Detection of WGM peaks with *ksrates* was performed using exponential-lognormal mixture modeling and lognormal-only mixture modeling for whole-paranome *K*_S_ distributions, anchor pair *K*_S_ clustering and lognormal mixture modeling for anchor pair *K*_S_ distributions, and lognormal mixture modeling for RR GF *K*_S_ distributions. For the (exponential-)lognormal mixture modeling, models with an increasing number of components were fitted, and the model with the lowest Bayesian Information Criterion (BIC) score was selected. For anchor pair *K*_S_ clustering, the number of components was automatically calculated based on the segmental duplication level detected by the i-ADHoRe 3.0 tool (Proost *et al*., 2011) incorporated in *ksrates*.

For the whole-paranome *K*_S_ distributions, exponential-lognormal mixture modeling was most often preferred to lognormal-only mixture modeling based on the BIC score, as exponential-lognormal mixture modeling better accounts for the small-scale duplication (SSD) background. For the whole-paranome *K*_S_ distribution of *B. distachyon*, the lognormal-only mixture model had a slightly lower BIC score (BIC=26045) than the exponential-lognormal mixture model (BIC=29057), but we selected the exponential-lognormal mixture model (Suppl. Fig. S12) because it modeled the SSD background more realistically. For the anchor pair *K*_S_ distributions, anchor pair *K*_S_ clustering was generally preferred over lognormal mixture modeling as the clustering procedure also incorporates synteny information from WGM-derived duplicated regions (Sensalari *et al*., 2021). Exceptions include *D. carota* and *C. sativum* in Suppl. Fig. S13, because in both cases the two Apioideae-specific WGMs were modeled by a single cluster when using anchor pair *K*_S_ clustering.

## 5 Supplementary Material

**Supplementary material.pdf**: Supplementary methods, figures and tables

**Data S1.xlsx**: Gene family rankings in terms of reciprocal retention strength based on different datasets

**Data S2.xlsx**: cLRT results on the 37-species dataset

**Data S3.xlsx**: Gene IDs and gene family counts for the 37-species dataset, extended datasets and independent clade datasets

**Data S4.xlsx**: WGM inference results from the WGDgc analyses

**Data S5.xlsx**: Phylogenetic trees and WGM positioning for the analyzed datasets

Code for the birth-death model and cLRTs is available at https://github.com/VIB-PSB/BirthDeathModel. *ksrates* v2.0 software and code is available at https://github.com/VIB-PSB/ksrates. *OrthoMCLight* is available at https://github.com/VIB-PSB/OrthoMCLight. All input and output files for the cLRT and WGDgc analyses can be found in the following zenodo archive: https://www.doi.org/10.5281/zenodo. 15225340.

## 6 Author contributions

S.M. designed and supervised the study. S.T. designed and implemented the first version of the cLRT modeling framework, which was extended by S.M. and C.S. All authors performed simulations and analyzed data. S.T. wrote an initial draft describing the cLRT framework and the first simulation results. C.S. and S.M. wrote the final manuscript incorporating additional results and analyses.

## 7 Acknowledgments

This work was supported by a grant from the Research Foundation Flanders (FWO grant G.0088.12). We acknowledge Michiel Van Bel for implementation of OrthoMCLight, Kevin De Pelseneer for CLI implementation of the cLRT workflows and Stijn Hawinkel for advice on calculating standard errors on the *λ* estimates. The computational resources (Stevin Supercomputer Infrastructure) and services used in this work were provided by the VSC (Flemish Supercomputer Center), funded by Ghent University, FWO and the Flemish Government – department EWI. We thank Daniel Cruz, Michael Van de Voorde and Stijn Hawinkel for help with the HPC simulations, and Bert Droesbeke for technical assistance with *ksrates* distribution.

## Notes

https://www.doi.org/10.5281/zenodo.15225340

https://github.com/VIB-PSB/BirthDeathModel

https://github.com/VIB-PSB/ksrates

https://github.com/VIB-PSB/OrthoMCLight

### Competing Interest Statement

The authors have declared no competing interest.

