## Supplementary Methods, Figures and Tables for "Using reciprocally retained gene families to detect whole-genome multiplications in plants"

April 17, 2025

### 1 Supplementary Methods

#### 1.1 Comparison between original and extended GFs

To assess the GF reconstruction quality, we computed the Euclidean distance between the original and the reconstructed gene count profiles for each of the 21 marker GFs across the 37 species (Figure M1). In any extended dataset, the gene count Euclidean distance generally ranges between 0 and 4, with a maximum of 7 for a few GFs. To assess the effect of these deviations in gene counts on $\lambda$ estimation, we computed the maximum-likelihood estimates for the $\lambda$ parameters of the 21 marker GFs on the phylogenetic trees and gene count profiles of the extended datasets (see Methods) excluding the additional clades. When using the new trees and gene count profiles, the $\lambda^*$ deviations from the values obtained on the original 37-species dataset are generally fairly small and correlate with the deviations in the gene count profiles, as expected (Figure M1).

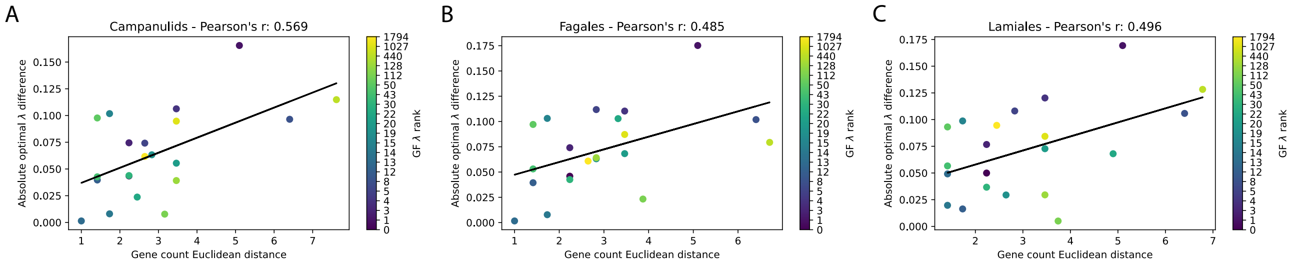


Figure M1: Scatter plots showing the absolute difference between the optimal $\lambda$ computed for the 21 candidate marker GFs on the original 37 species dataset and on an extended dataset (ignoring the added clade) as a function of the Euclidean distance between the corresponding gene count profiles of the marker GFs. (A) Extended dataset for Campanulids. (B) Extended dataset for Fagales. (C) Extended dataset for Lamiales. The Pearson correlation coefficients given on top of the panels indicate a moderate positive correlation between $\lambda$ differences and gene count profile distances. Black lines are linear regression lines. The dots representing the marker GFs are color-coded according to their rank, see the legend on the right of each panel.

#### 1.2 Placement of WGMs on extended trees based on averaged t distances calculated for original 37-species tree

For positioning the 20 WGMs of the 37-species dataset on a phylogenetic tree constructed for an extended set of species (37 species plus an additional clade), we reused the averaged t distances reported in Tasdighian *et al.* (2017). For any given WGM, this averaged t distance is the average of the t distances to the root (of the original 37-species tree) of the WGM positions obtained using the WGM t ages inferred from the anchor pair t distributions for different species that share the WGM (see Methods). The original WGM t ages from Tasdighian *et al.* (2017) were not stored permanently and could hence not be reused for positioning the WGMs on the extended trees, but here we prove that the averaged t distances can be used to the same effect (a solution that is computationally much less costly than recalculating the WGM t ages). Let $R_o\overline{W}_o$ be the averaged t distance for a given WGM, i.e. the distance from the root $R_o$ of the original tree $o$ to the averaged position $\overline{W}_o$ of the WGM (see Suppl. Fig. M2), based on positions $W_{os}$ calculated from the WGM ages inferred for the different species $s = 1 \ldots l$ that share the WGM. Let $L_{os}$ be the position on the original tree of the leaf node for species $s$. $W_{os}L_{os}$ is then half of the (unknown) WGM t age inferred from species $s$ (see Methods). Furthermore, let $\overline{W}_oL_{os}$ be the distance from the consensus position of the WGM to the leaf node for species $s$ in the original tree $o$, which is easily computed given the known averaged t distance of the WGM and the known original phylogeny:

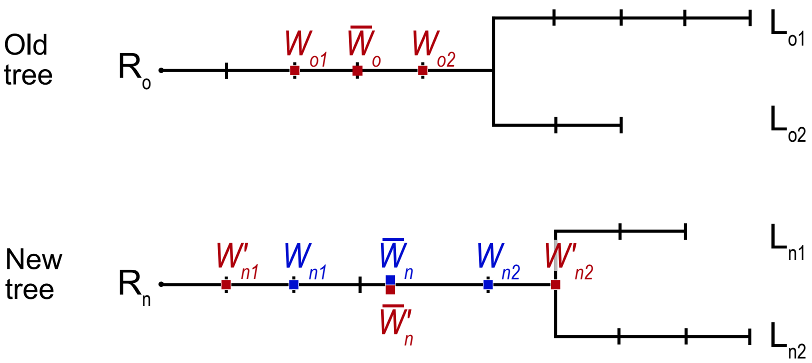


Figure M2: Example illustrating the equivalence of using the averaged t distance from an original tree or species-specific WGM t ages to position a WGM on a new tree. The nomenclature of the nodes is described in the text. Red font node labels illustrate WGM positions obtained using species-specific WGM t ages ($2 \cdot W_{o1}L_{o1} = 2 \cdot W_{n1}L_{n1}$ for species 1 and $2 \cdot W_{o2}L_{o2} = 2 \cdot W_{n2}L_{n2}$ for species 2). Blue font node labels illustrate WGM positions on the new tree $n$ obtained when using the averaged t distance $R_o\overline{W}_o$ from the original tree $o$. Both methods lead to the same consensus position on the new tree ($\overline{W}_n = \overline{W}'_n$). Root node: R; leaf node: L; WGM position: $W$.

$$\overline{W}_o L_{os} = R_o L_{os} - R_o \overline{W}_o \tag{1}$$

with $R_o L_{os}$ the distance from the root of tree $o$ to leaf node $s$.

For the new tree $n$, similar naming conventions hold. For simplicity, we assume in the following that the WGM is shared by only two species, but the reasoning is valid also for more species. Using the (unknown) WGM t ages $2 \cdot W_{o1}L_{o1}$ and $2 \cdot W_{o2}L_{o2}$, the consensus distance $R_n\overline{W}'_n$ from the WGM to the root $R_n$ would be calculated as (Suppl. Fig. M2 in red) :

$$R_n \overline{W}'_n = \frac{R_n W'_{n1} + R_n W'_{n2}}{2} \tag{2}$$

with

$$\begin{aligned} R_n W'_{n1} &= R_n L_{n1} - W_{o1} L_{o1} \\ R_n W'_{n2} &= R_n L_{n2} - W_{o2} L_{o2} \end{aligned} \tag{3}$$

Using the distances $\overline{W}_o L_{o1}$ and $\overline{W}_o L_{o2}$ from the consensus position of the WGM to the leaf nodes for species 1 and 2 in the original tree $o$, as calculated from the available averaged t distance $R_o\overline{W}_o$ and the original tree $o$ using equation 1, the consensus distance $R_n\overline{W}_n$ of the WGM from the root was calculated as :

$$R_n \overline{W}_n = \frac{R_n W_{n1} + R_n W_{n2}}{2} \tag{4}$$

with

$$\begin{aligned} R_n W_{n1} &= R_n L_{n1} - \overline{W}_o L_{o1} \\ R_n W_{n2} &= R_n L_{n2} - \overline{W}_o L_{o2} \end{aligned} \tag{5}$$

i.e., from each species leaf node $s$, the species' lineage was traced backwards and the WGM was placed at a distance equal to $\overline{W}_o L_{os}$. Due to the fact that leaves have different terminal branch lengths, this step produces multiple species-specific WGM positions ($W_{ns}$, see Suppl. Fig. M2 in blue) that act as a replacement for the positions that are normally inferred from anchor pair-derived WGM t ages.

The following proves that the distances $R_n\overline{W}_n$ derived from the averaged t distance and $R_n\overline{W}'_n$ derived from the species-specific WGM t ages are equivalent:

$$
\begin{gathered}
R_n\overline{W}_n \overset{?}{=} R_n\overline{W}'_n \\
\tfrac{(R_nL_{n1}-\overline{W}_oL_{o1})+(R_nL_{n2}-\overline{W}_oL_{o2})}{2} \overset{?}{=} \tfrac{(R_nL_{n1}-W_{o1}L_{o1})+(R_nL_{n2}-W_{o2}L_{o2})}{2} \\
\overline{W}_oL_{o1} + \overline{W}_oL_{o2} \overset{?}{=} W_{o1}L_{o1} + W_{o2}L_{o2} \\
(R_oL_{o1} - R_o\overline{W}_o) + (R_oL_{o2} - R_o\overline{W}_o) \overset{?}{=} (R_oL_{o1} - R_oW_{o1}) + (R_oL_{o2} - R_oW_{o2}) \\
-2R_o\overline{W}_o \overset{?}{=} -R_oW_{o1} - R_oW_{o2} \\
R_o\overline{W}_o \overset{!}{=} \tfrac{R_oW_{o1}+R_oW_{o2}}{2}
\end{gathered}
\tag{6}
$$

#### 1.3 Assessing the variance of the optimal lambda estimate

The uncertainty around the maximum-likelihood estimate (MLE) $\lambda_{max}$ obtained from the observed gene count data $X$ for a given GF was computed by means of the Cramér-Rao bound, which gives a lower bound for the variance of an unbiased estimator. The MLE achieves this lower bound and is hence as efficient as possible. The Cramér-Rao bound for the variance of $\lambda_{max}$ is given as the reciprocal of the Fisher information $I(\lambda_{max})$ multiplied by the number of data points $n$ (= 37, the number of gene counts in the 37-species tree)(Nielsen, 2013):

$$
Var(\lambda_{max}) \geq \frac{1}{nI(\lambda_{max})} \tag{7}
$$

The Fisher information $I(\lambda_{max})$ is related to the curvature of the log-likelihood function evaluated at its maximum:

$$
I(\lambda_{max}) = -E\Big[\frac{\partial^2 \log f(X;\lambda)}{\partial\lambda^2}\Big|\lambda_{max}\Big] \tag{8}
$$

where $\log f(X;\lambda)$ is the log-likelihood function of $X$ given $\lambda$. To compute the curvature of $\log f$, we generated for each GF $j$ a range of 100 $\lambda_j$ values around the previously obtained $\lambda_j^*$ estimates with a step interval of 0.001. The log-likelihood of the BD model on the observed gene counts was evaluated for each $\lambda_j$ conditioned on the previously estimated optimal root size $r_j^*$. The resulting log-likelihood series revealed that the $\lambda_j^*$ values were not always fully optimized (see e.g. Suppl. Fig. M3). The $\lambda_{max}$ estimate for use in Eq. 7 and 8 was therefore defined as the sampled $\lambda_j$ value that yielded the highest log-likelihood. The log-likelihood function $f(X|\lambda)$ was approximated by fitting a third degree polynomial with ordinary least squares through the evaluated $\lambda_j$–log-likelihood pairs. The second derivative of this polynomial at $\lambda_{max}$ was then used to compute the Fisher information (Eq. 8) and the variance on the $\lambda_{max}$ estimate (Eq. 7). The standard error associated with $\lambda_{max}$ was obtained by taking the square root of the variance. The standard error linearly increases with $\lambda_{max}$ (Pearson's r = 0.908, see Suppl. Fig. M4A), but does not surpass 0.05 for any of the candidate WGM marker GFs. The

$\lambda^*$ estimates inferred from the BD model are generally close to the corresponding $\lambda_{max}$ values, and in all cases well within the standard error on $\lambda_{max}$ (see Suppl. Fig. M4B).

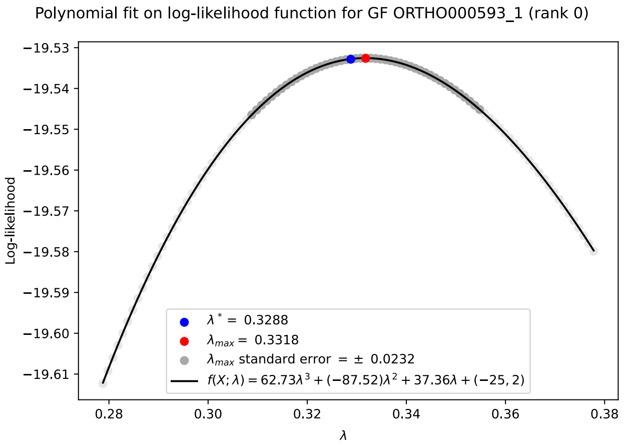


Figure M3: Analysis of the uncertainty on the $\lambda$ maximum likelihood estimate ($\lambda_{max}$) for gene family ORTHO000593_1 (rank 0). 100 $\lambda$ values were sampled around the $\lambda^*$ estimate produced by the BD model (blue dot) to approximate the log-likelihood function (black line). $\lambda_{max}$ (red dot) was defined as the sampled $\lambda$ value producing the highest likelihood. Sampled $\lambda$ values falling within the standard error range of $\lambda_{max}$ are highlighted in dark gray.

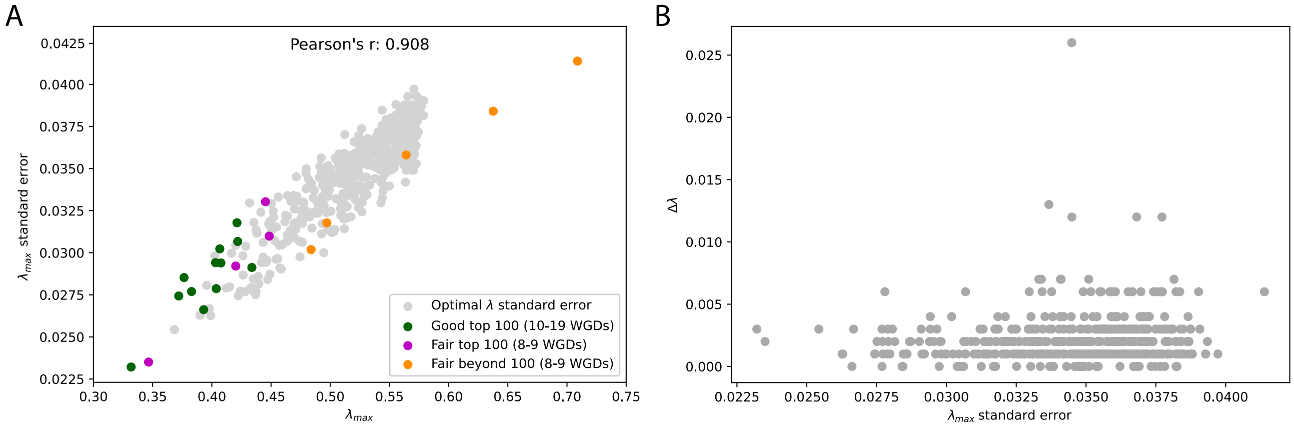


Figure M4: Standard error on $\lambda_{max}$ values. (A) The standard error of $\lambda_{max}$ as a function of $\lambda_{max}$. Dots indicate data points for the top 500 GFs plus two candidate marker GFs ranked higher. Green dots correspond to candidate marker GFs detecting $\geq$ 10 WGMs in the 37-species dataset, while magenta and orange dots correspond to candidate marker GFs detecting 8-9 WGDs and belonging to the top 100 or top 101-2000 group, respectively. (B) Absolute deviation of the lambda estimate from the BD model ($\lambda^*$) from the $\lambda_{max}$ estimate ($|\Delta\lambda|$), as a function of the $\lambda_{max}$ standard error.

### 2 Supplementary Figures

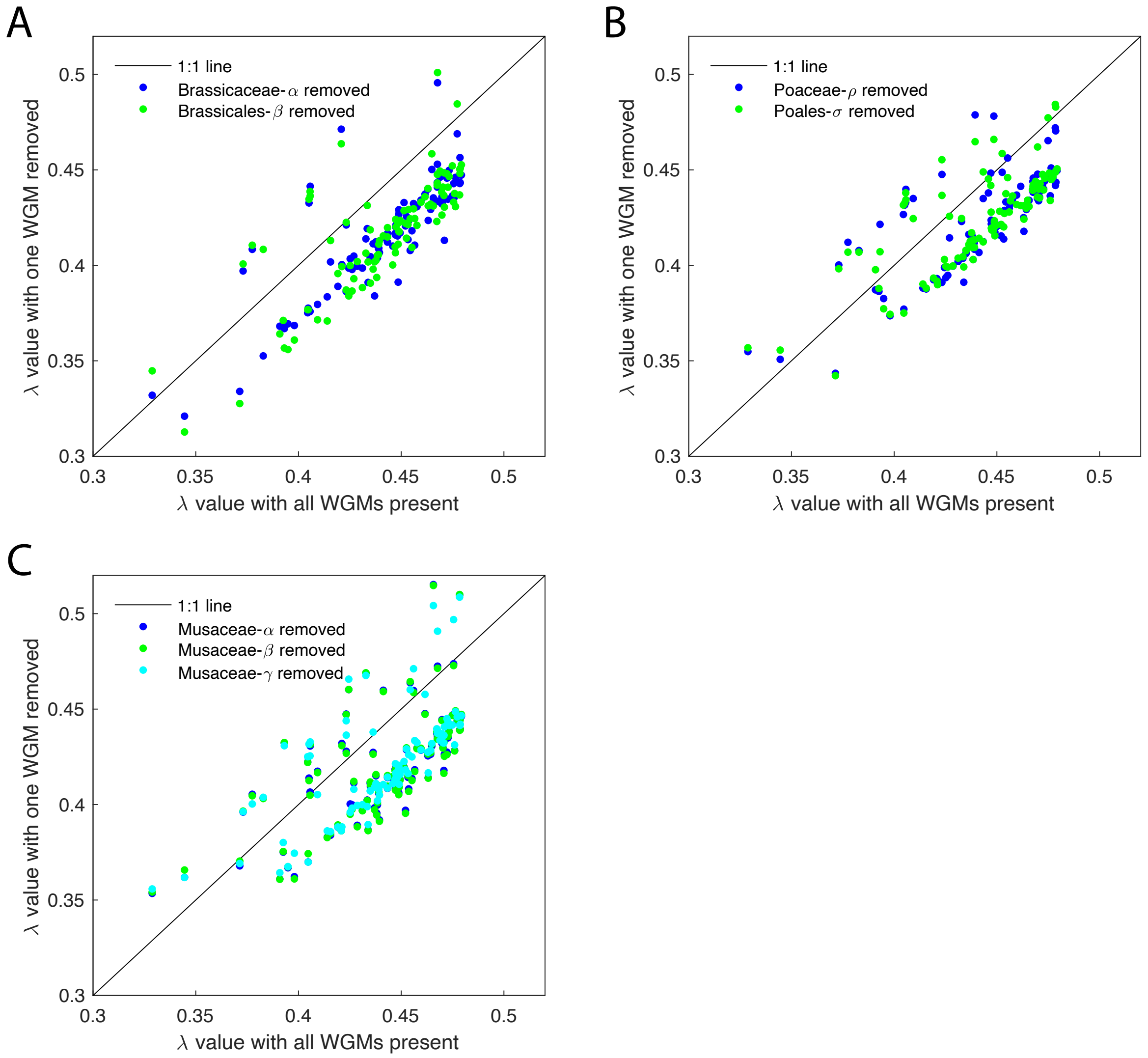


Figure S1: Effect of removing one of several WGMs on the same branch on $\lambda$ estimates. (A) Effects on $\lambda$ of modeling the absence of Brassicaceae-$\alpha$ or Brassicales-$\beta$ WGDs. (B) Effects on $\lambda$ of modeling the absence of the Poaceae-$\rho$ or Poales-$\sigma$ WGDs. (C) Effects on $\lambda$ of modeling the absence of the $\alpha$, $\beta$ or $\gamma$ WGD in the Musaceae. Each dot represents one of the top-100 GFs.

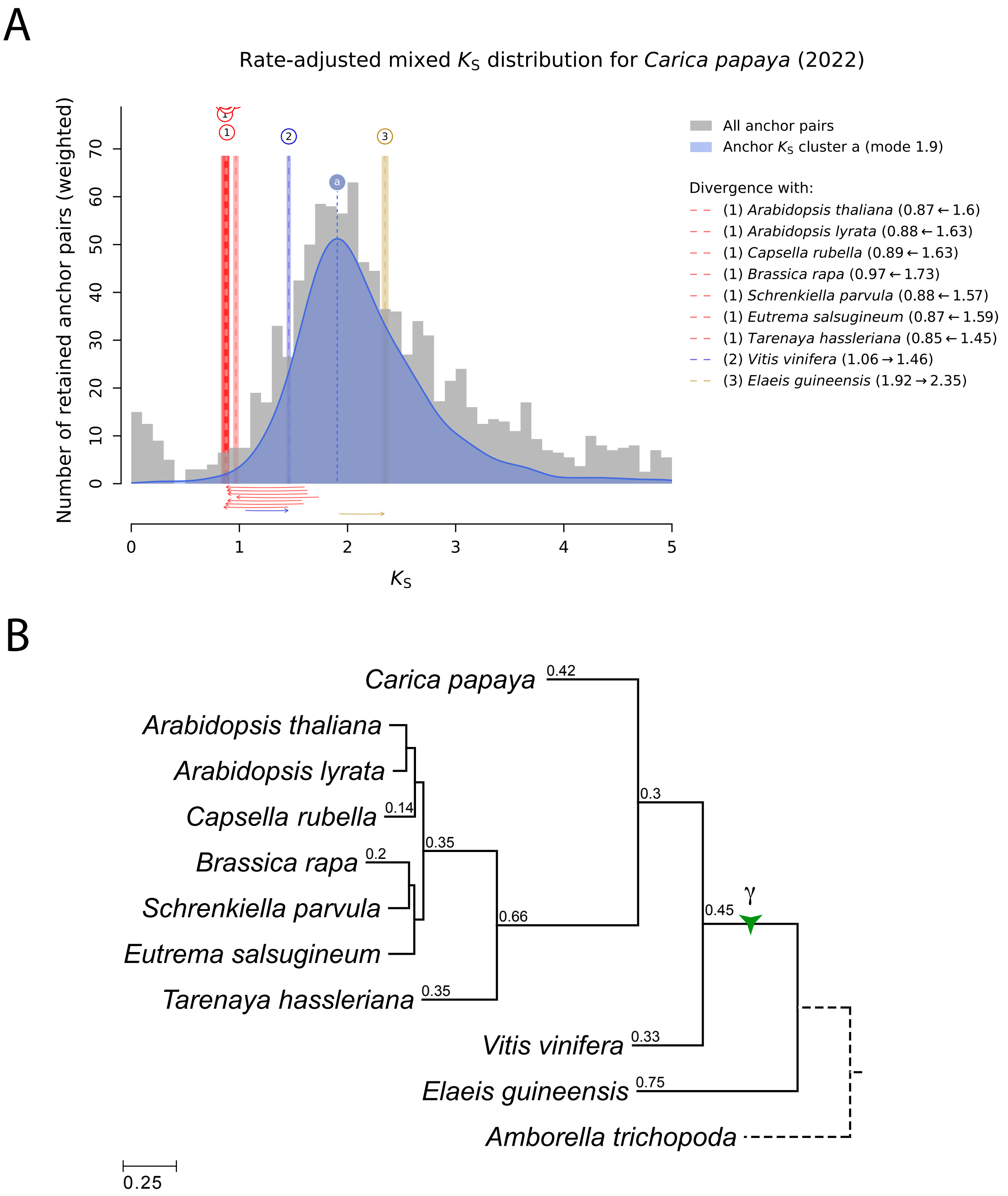


Figure S2: WGM inference in *Carica papaya* by means of substitution rate-adjusted mixed $K_S$ distribution analysis using `ksrates`. (A) Anchor pair $K_S$ clustering supporting the presence of a single WGM (blue cluster `a`): the core-eudicot $\gamma$ WGT shared by all the eudicot species included in the analysis, but not by the monocot *E. guineensis* (oil palm). Vertical lines labeled with numbers indicate the modes of ortholog $K_S$ distributions, which mark the divergence between *Carica papaya* and other species. Colored boxes range from one standard deviation (SD) below to one SD above the mean mode estimate. The $K_S$ ages of ortholog $K_S$ modes have been adjusted on the basis of substitution rate differences between species, as illustrated by the horizontal arrows above the x-axis. (B) Phylogenetic tree generated by `ksrates`, with the $\gamma$ WGT manually positioned on the tree as a green three-pointed star. Branch lengths are given in $K_S$ units.

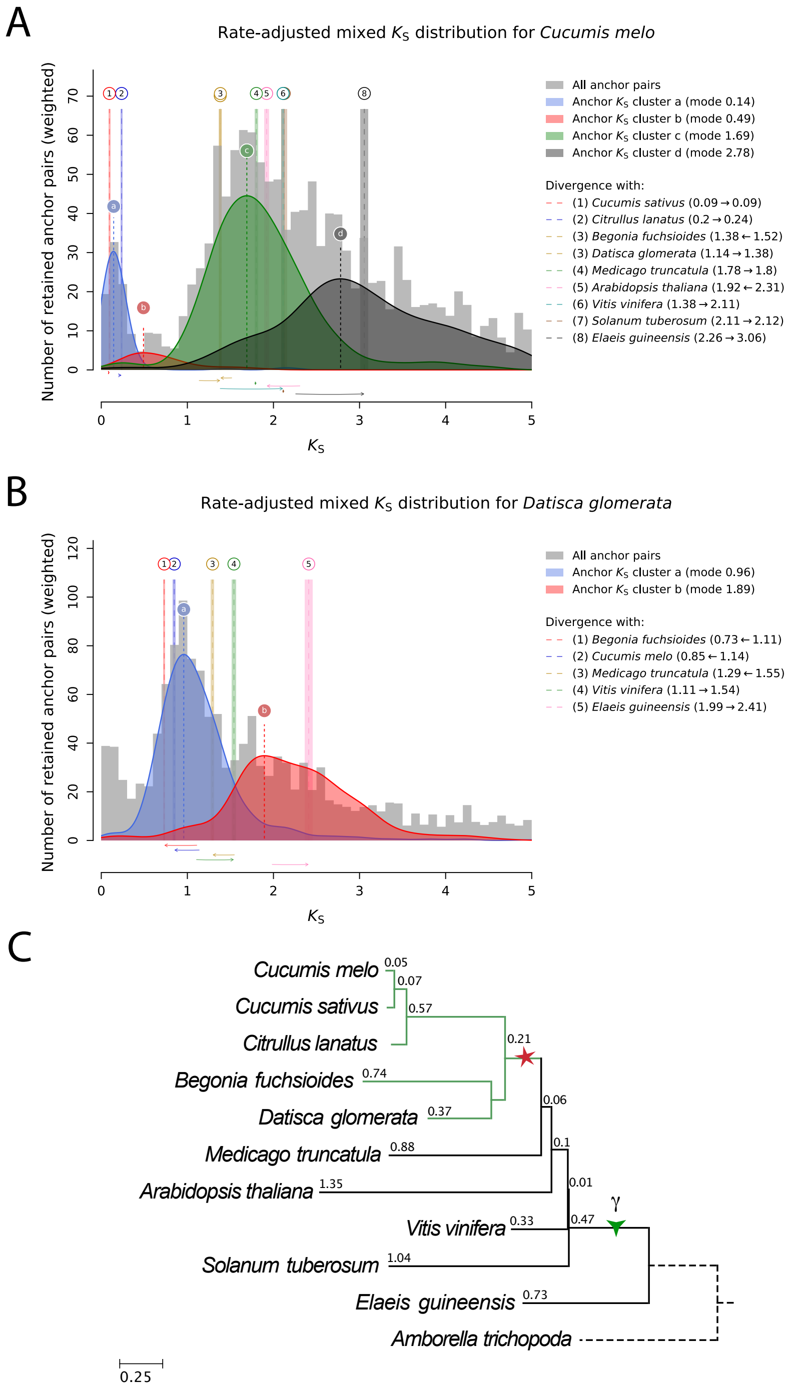


Figure S3: WGM inference in the Cucurbitales by means of substitution rate-adjusted mixed $K_S$ distribution analysis using `ksrates`. (A) Anchor pair $K_S$ clustering for *Cucumis melo* (melon), supporting the presence of a Cucurbitales-specific WGM (green cluster `c`). (B) Anchor pair $K_S$ clustering for *Datisca glomerata*, also supporting the presence of a Cucurbitales-specific WGM (blue cluster `a`). In both (A) and (B), there is also evidence for a WGM shared by all eudicot species analyzed but not by the monocot *Elaeis guineensis* (black cluster `d` in (A) and red cluster `b` in (B)), which corresponds to the core-eudicot $\gamma$ WGT. Vertical lines labeled with numbers in (A) and (B) indicate the modes of ortholog $K_S$ distributions, which mark the divergence between the focal species and other species. Colored boxes range from one standard deviation (SD) below to one SD above the mean mode estimate. The $K_S$ ages of ortholog $K_S$ modes have been adjusted on the basis of substitution rate differences between species, as illustrated by the horizontal arrows above the x-axis. (C) Phylogenetic tree generated by `ksrates`, with the two WGMs manually positioned and marked with red and green stars. The Cucurbitales clade has been highlighted in green. Branch lengths are given in $K_S$ units.

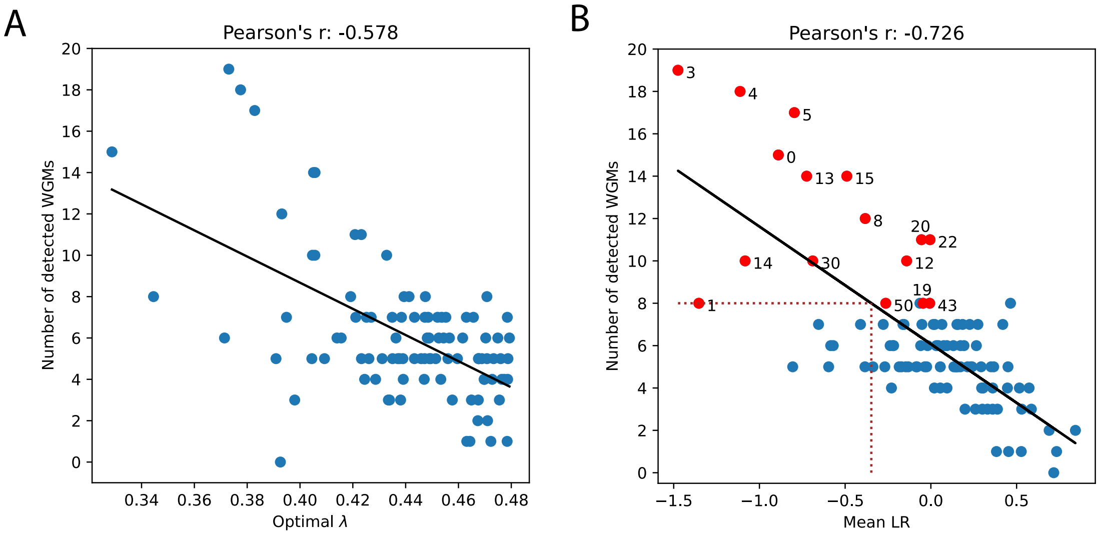


Figure S4: (A) Number of detected WGMs as a function of the estimated optimal $\lambda$ for the top-100 GFs. (B) Number of detected WGMs as a function of the mean observed likelihood ratio (mean LR) computed for the top-100 GFs, with means calculated over the LR values for $H_0$ = all WGMs present versus $H_1$ = one WGM absent for all 20 true WGMs in the 37-species tree. The dotted line in (B) marks the expected mean LR of GFs detecting at least 8 WGMs (-0.3472), which was used as the LR threshold to select additional GFs in the top 101-2000 range. The 16 best-performing GFs in the top-100 (at least 8/20 true WGMs detected, maximum 1 true WGM rejected and maximum 2 conflicting outcomes) are highlighted in red. The black lines in (A) and (B) are linear regression lines.

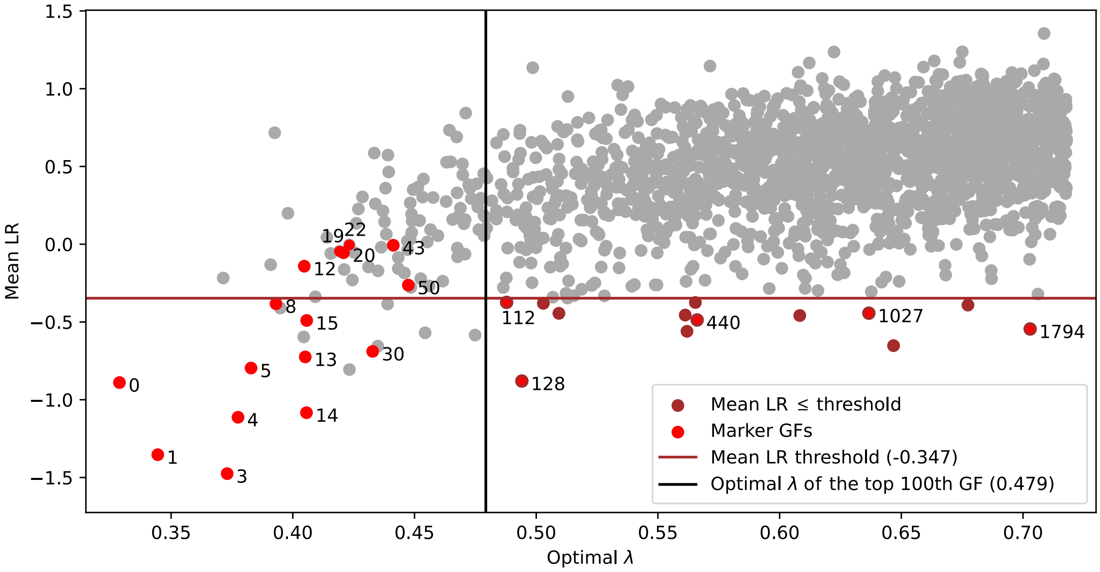


Figure S5: Scatter plot of the mean observed LR (mean LR) as a function of the optimal $\lambda$ for the top 2000 GFs, with means calculated over the LR values for the 20 true WGMs in the 37-species tree. The black vertical line separates the top 100 GFs from the top 101-2000 GFs. The dark red horizontal line indicates the LR threshold ($-0.3472$) for selecting potentially interesting GFs in the top 101-2000, as derived from Fig. S4B. Dark red dots and bright red dots with dark red borders in the bottom-right quadrant indicate the 13 GFs in the top 101-2000 with mean LR values below this threshold, for which full cLRTs were performed (GFs with ranks 112, 128, 155, 175, 402, 410, 435, 440, 756, 1027, 1126, 1466, 1794). Bright red dots indicate the 16 best-performing GFs from the top-100 and the additional 5 GFs in the top 101-2000 with comparable performance (at least 8/20 true WGMs detected, maximum 1 true WGM rejected and maximum 2 conflicting outcomes).

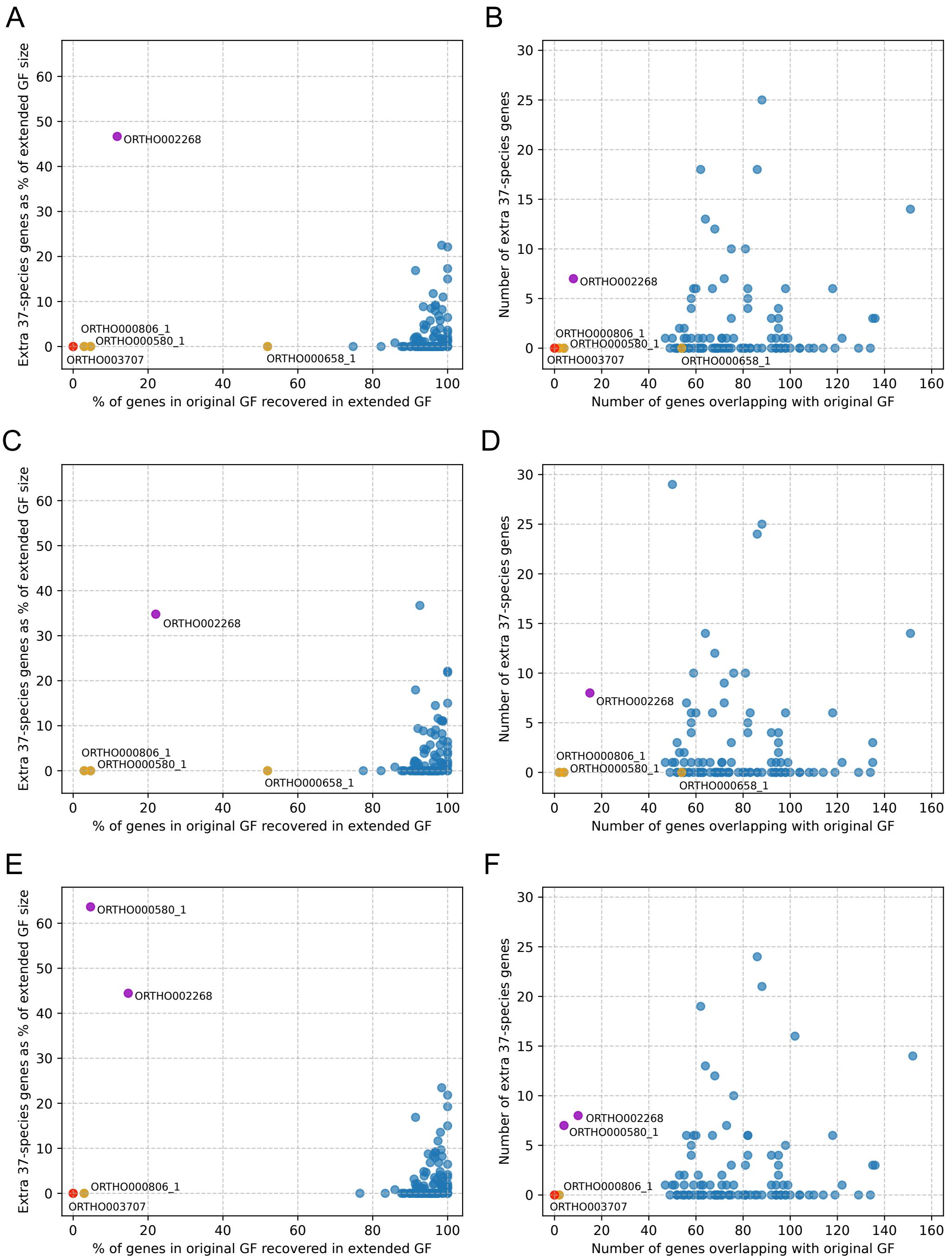


Figure S6: Overlap between the extended and original GFs and extra 37-species genes in the extended GFs. Each dot represents one of the top-100+5 GFs. Left plots show the percentage of genes in an original GF that are recovered in the extended GF on the x-axis and the extra 37-species genes as a percentage of the extended GF size on the y-axis, for the extended datasets incorporating Campanulids (A), Fagales (C) and Lamiales (E). Right plots show the corresponding numbers of overlapping genes and extra genes for Campanulids (B), Fagales (D) and Lamiales (F). GFs that could not be extended (no extended orthogroups mapping to original GF after filtering) are indicated with red dots. Extended GFs recovering ≤ 60% of the original GF's genes but not including additional 37-species genes are indicated with orange dots. Extended GFs recovering ≤ 25% of the original GF's genes and including substantial amounts of extra 37-species genes are indicated with purple dots. These three categories of extended GFs are the most dissimilar to the corresponding original 37-species GFs and hence likely also do not reliably represent the targeted RR GF in the added clade (Campanulids, Fagales or Lamiales).

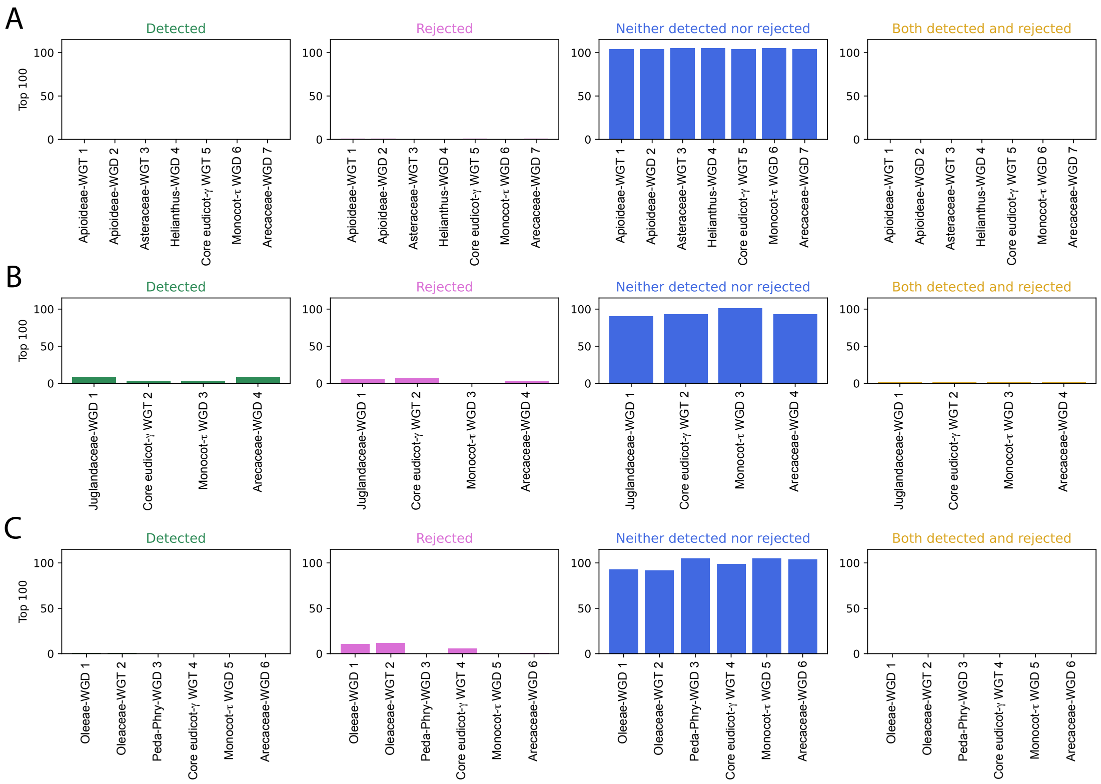


Figure S7: Per-WGM inference performance of the top 100+5 marker GFs on Campanulids (A) Fagales (B) and Lamiales (C).

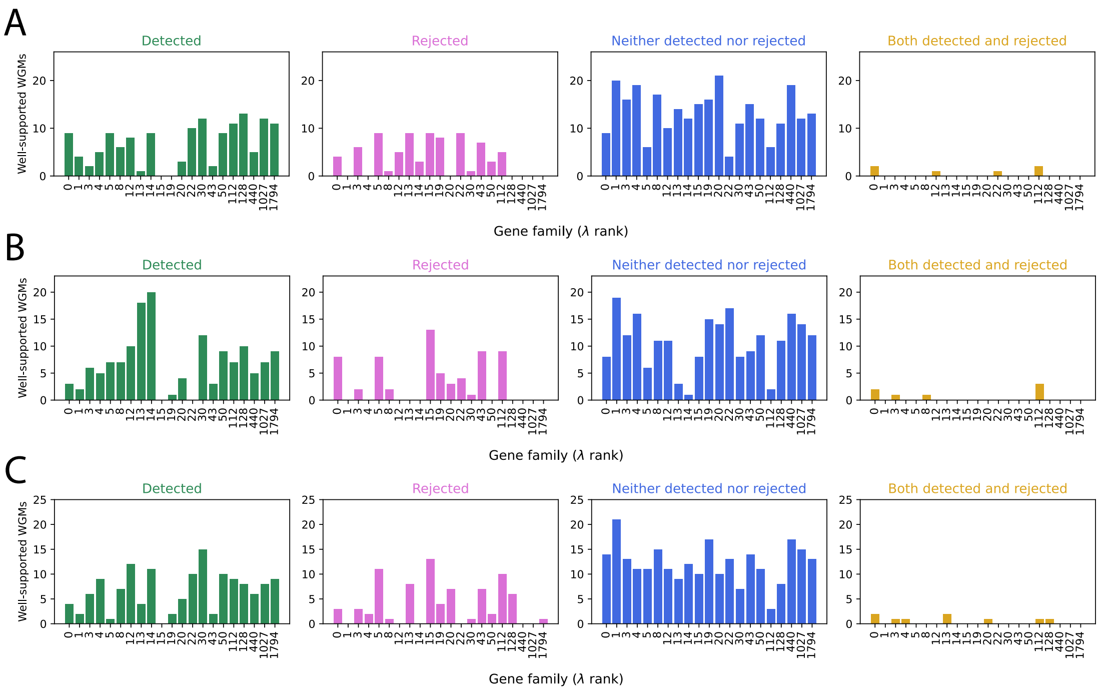


Figure S8: Per-GF inference performance of the 21 marker GFs on the three extended datasets, encompassing the original 37 species plus Campanulids (A), Fagales (B) or Lamiales (C).

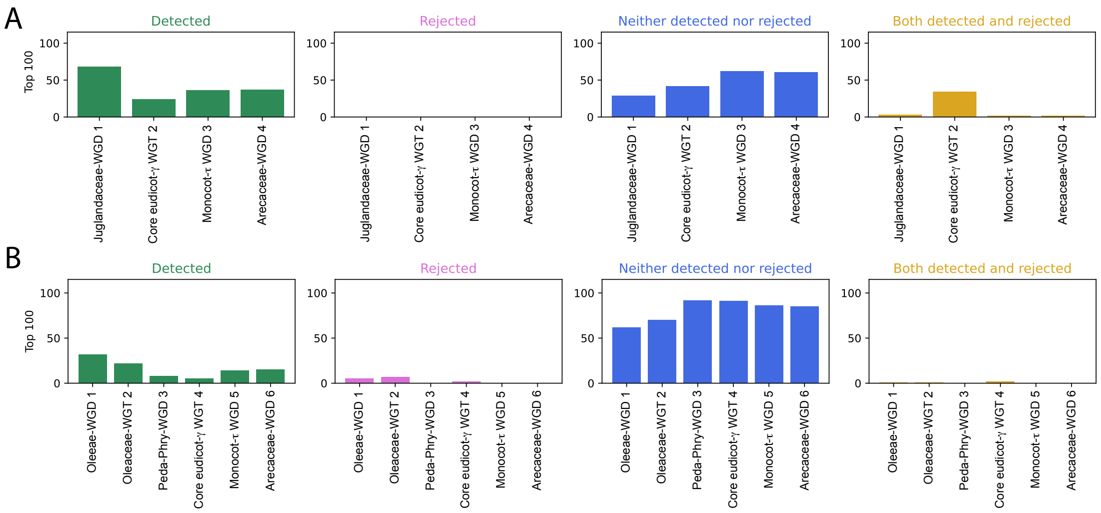


Figure S9: Per-WGM inference performance of the updated top-100 reconstructed gene families in Fagales (A) and in Lamiales (B), after having sorted the 9178 gene families according to their optimal $\lambda$ estimated from the respective clade dataset.

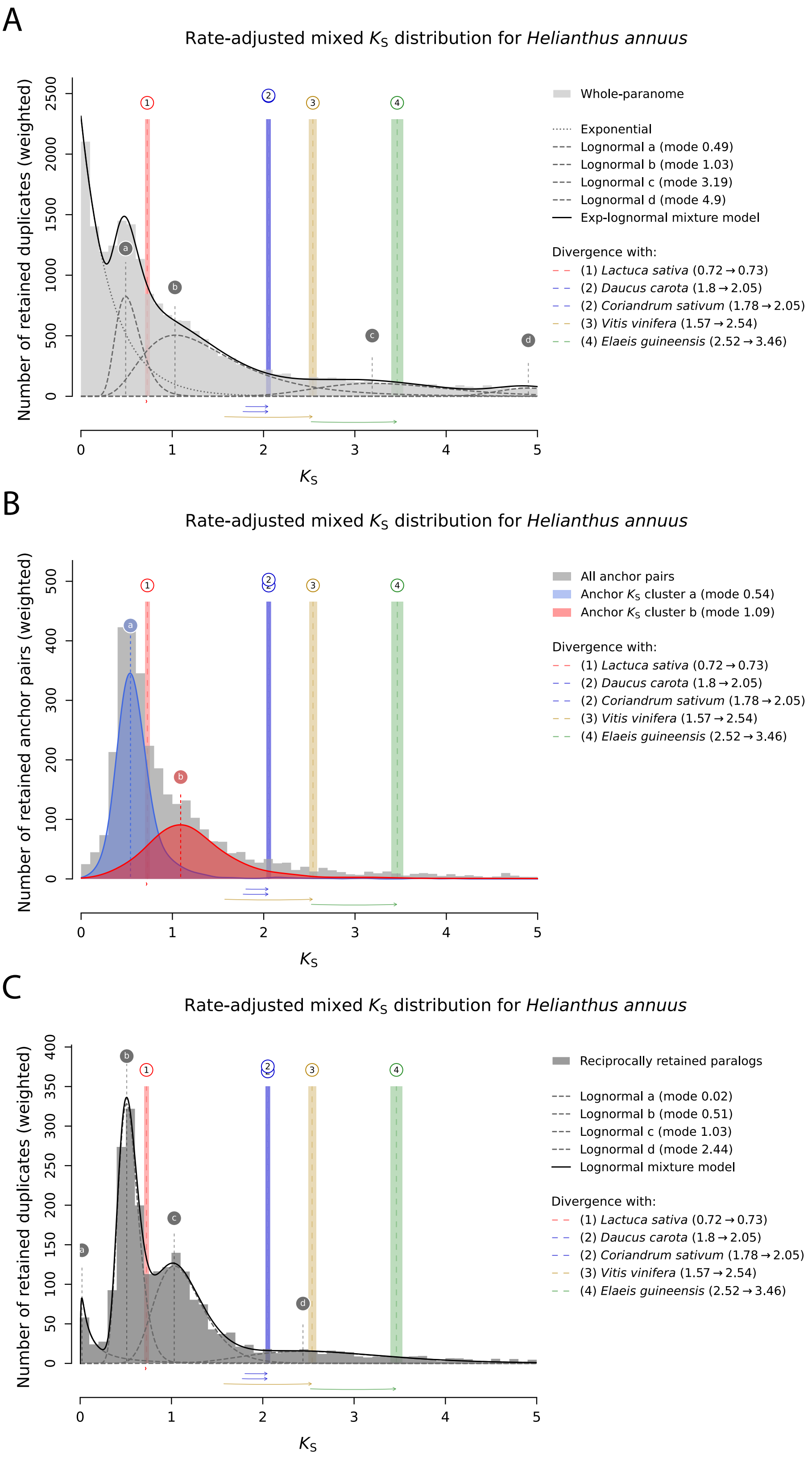


Figure S10: `ksrates`-based WGM peak inference in *Helianthus annuus* (sunflower, Campanulids). (A) Exponential-lognormal mixture model of the whole-paranome $K_S$ distribution. (B) Anchor pair clustering. (C) Lognormal mixture model of the $K_S$ distribution for the top-2000 RR GFs.

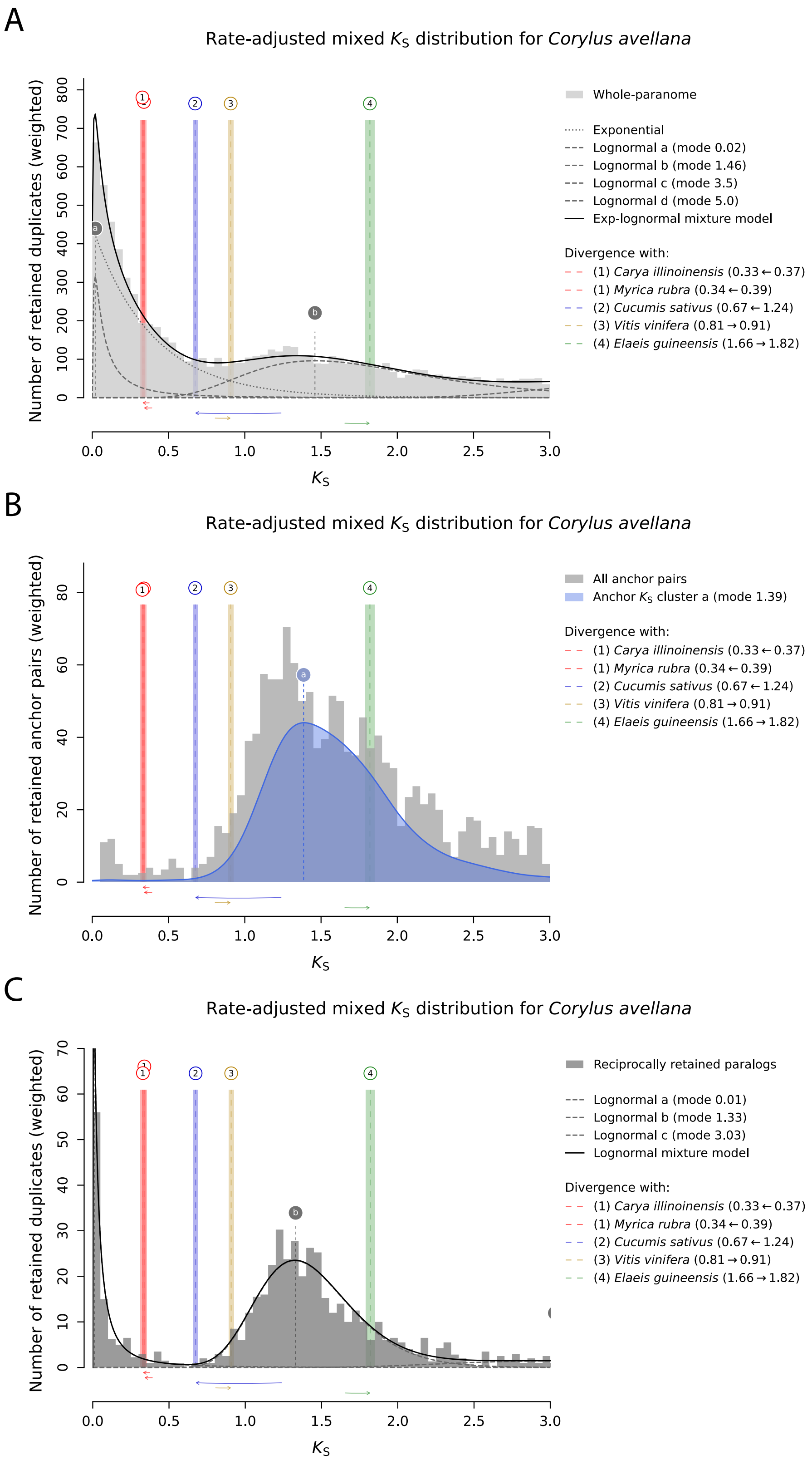


Figure S11: `ksrates`-based WGM peak inference in *Corylus avellana* (hazelnut, Fagales). (A) Exponential-lognormal mixture model of the whole-paranome $K_\mathrm{S}$ distribution. (B) Anchor pair clustering. (C) Lognormal mixture model of the $K_\mathrm{S}$ distribution for the top-2000 RR GFs.

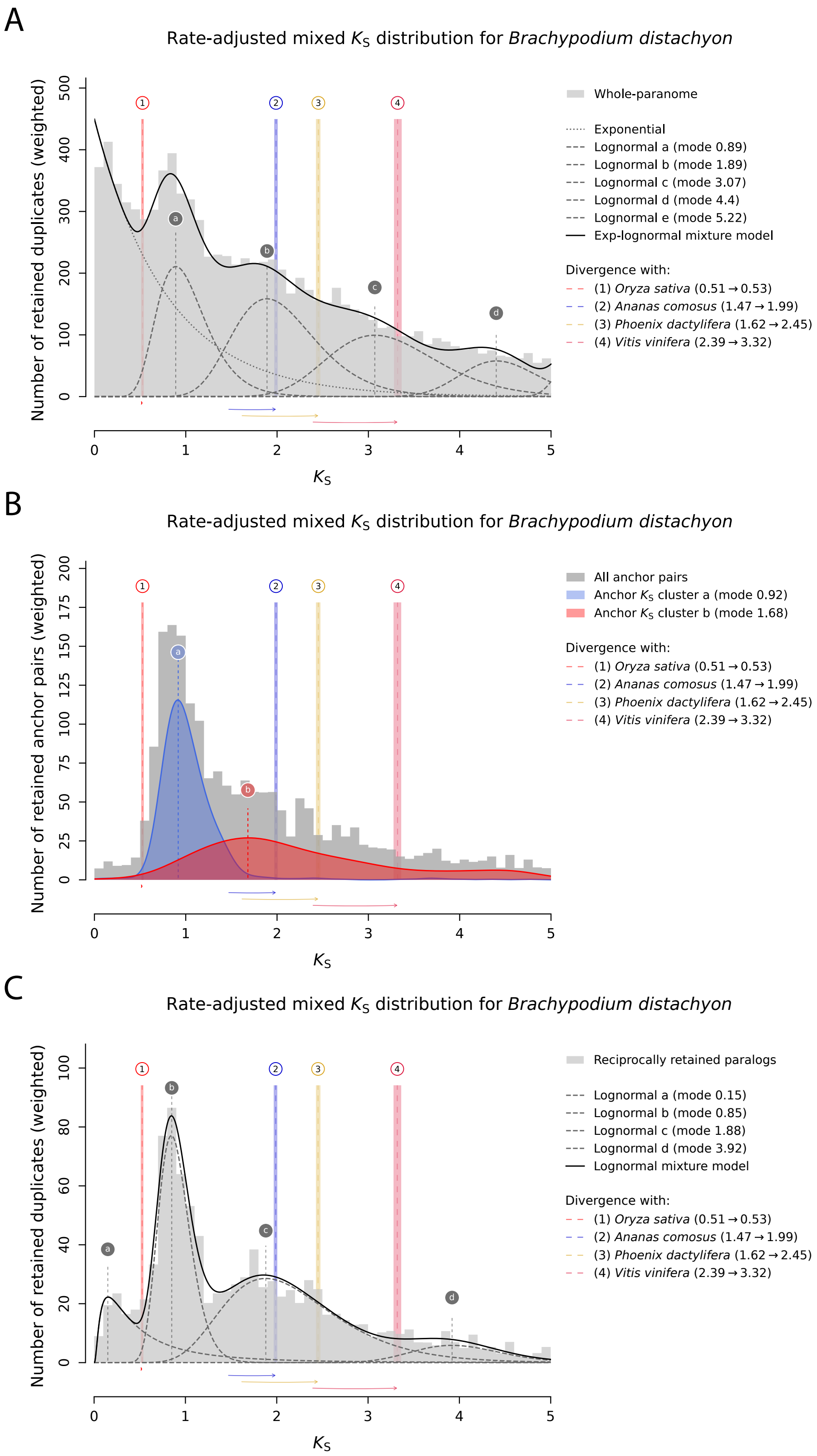


Figure S12: `ksrates`-based WGM peak inference in *Brachypodium distachyon* (Poaceae). (A) Exponential-lognormal mixture model of the whole-paranome $K_S$ distribution. (B) Anchor pair clustering. (C) Lognormal mixture model of the $K_S$ distribution for the top-2000 RR GFs. The top-2000 RR GFs are better able to discern the Poaceae $\rho$ and Poales $\sigma$ WGD peaks ($K_S \sim 0.9$ and ($K_S \sim 1.8$, respectively). The $\sigma$ and $\tau$ events cannot be separated out by either method, however.

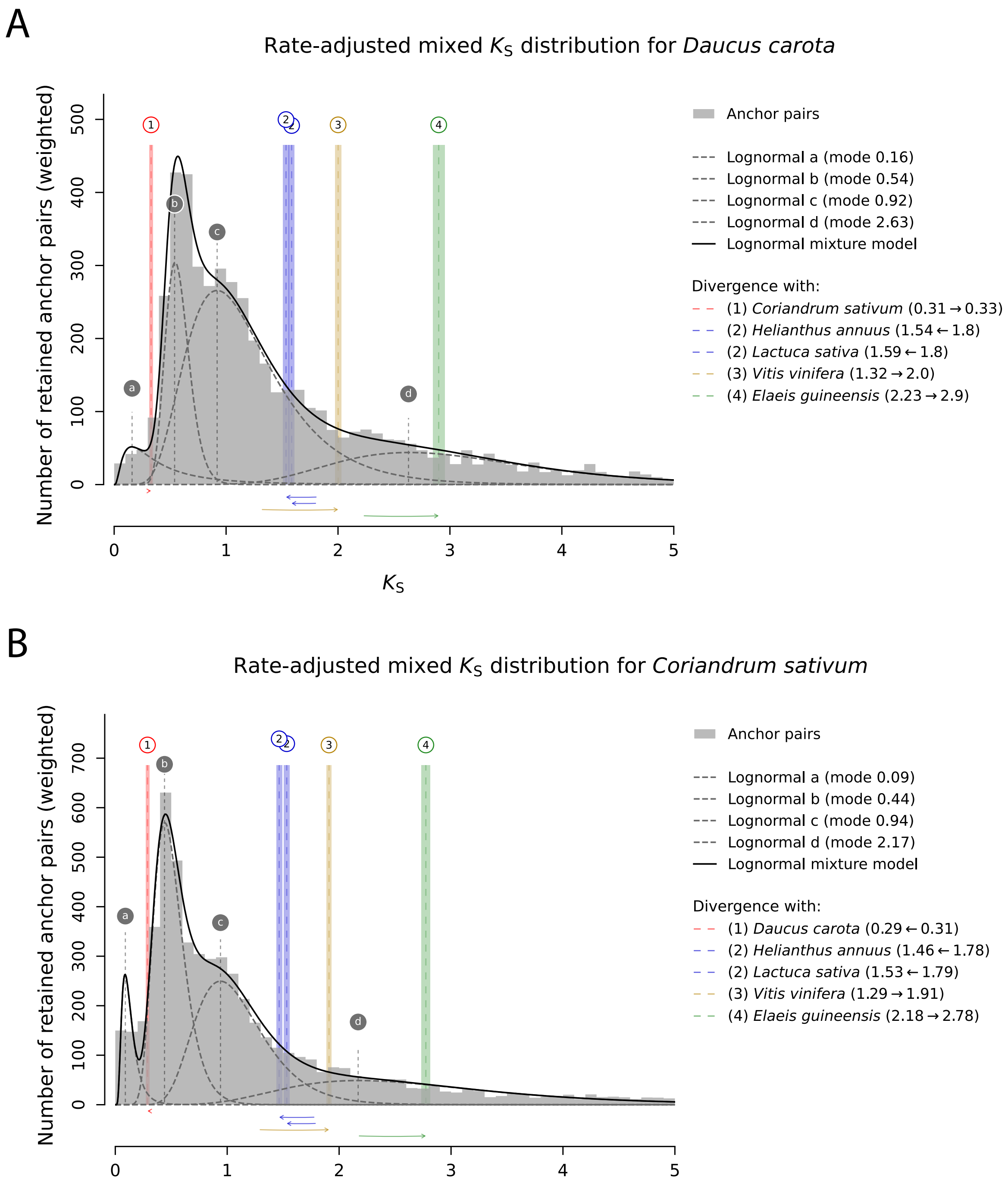


Figure S13: Substitution rate-adjusted mixed anchor pair $K_S$ distributions generated with `ksrates` for WGM inference and positioning in the Campanulids. The modes of the WGM peaks are shown as vertical lines labeled with letters. Vertical lines labeled with numbers indicate the modes of ortholog $K_S$ distributions, which mark the divergence between the focal species and other species. Colored boxes range from one standard deviation (SD) below to one SD above the mean mode estimate. The $K_S$ ages of ortholog $K_S$ modes have been adjusted on the basis of substitution rate differences between species, as illustrated by the horizontal arrows above the x-axis. (A) Lognormal mixture model of the anchor pair $K_S$ distribution for *Daucus carota*. (B) Lognormal mixture model of the anchor pair $K_S$ distribution for *Coriandrum sativum*. *D. carota* and *C. sativum* share the recent Apioideae WGD (peak `b` in (A) and (B)) and the older Apioideae WGT (peak `c` in (A) and (B)). Peak `d` in (A) and (B) indicates the core-eudicot $\gamma$ WGT.

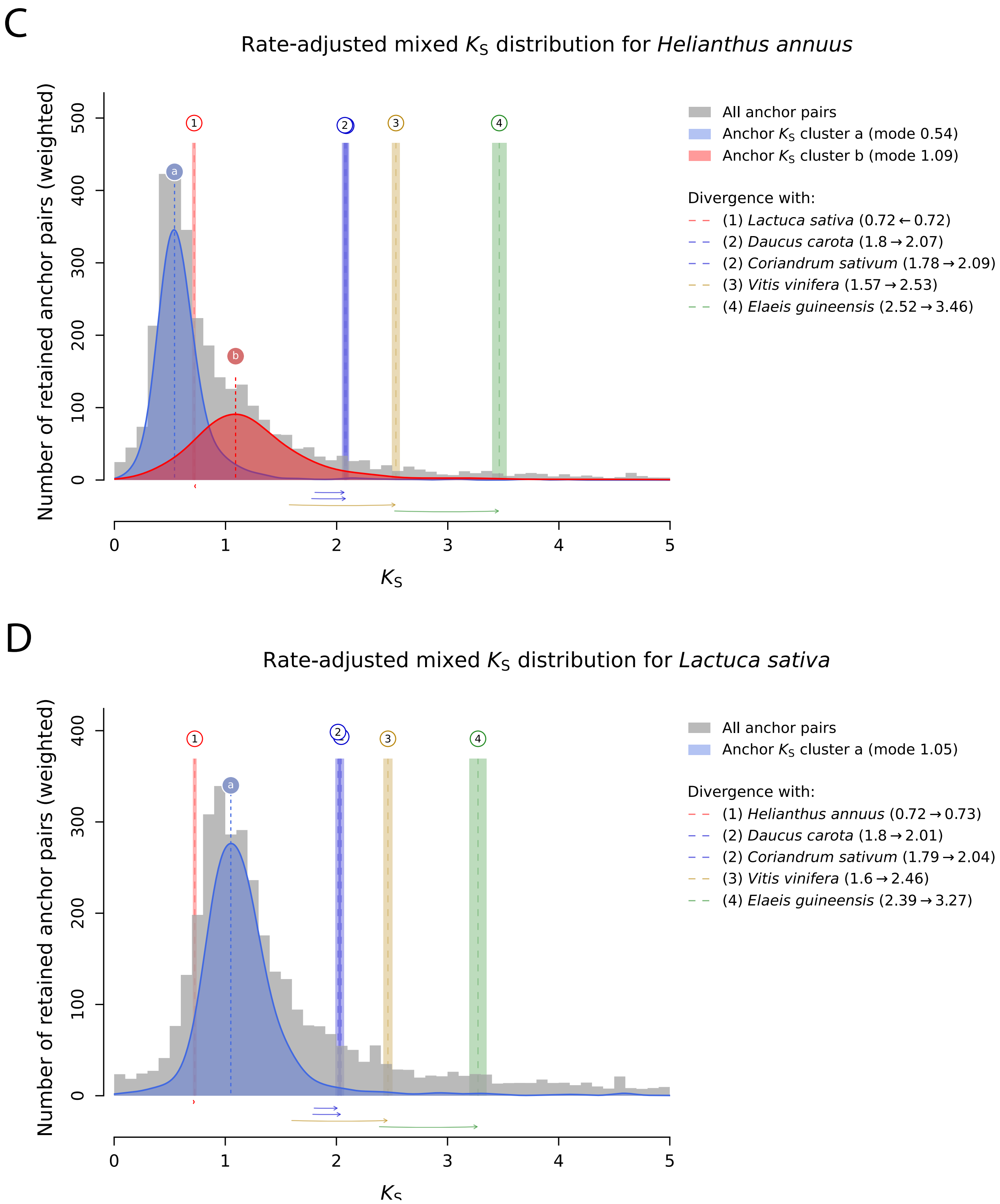


Figure S13: (continued) (C) Clustered anchor pair $K_S$ distribution for *Helianthus annuus*. (D) Clustered anchor pair $K_S$ distribution for *Lactuca sativa*. A recent WGD not shared with other Campanulids occurred in the Helianthus lineage (peak `a` in (A)). Remnants of an Asteraceae-specific WGT are present in the paranomes of both *Helianthus annuus* (peak `b` in (A)) and *Lactuca sativa* (peak `a` in (B)).

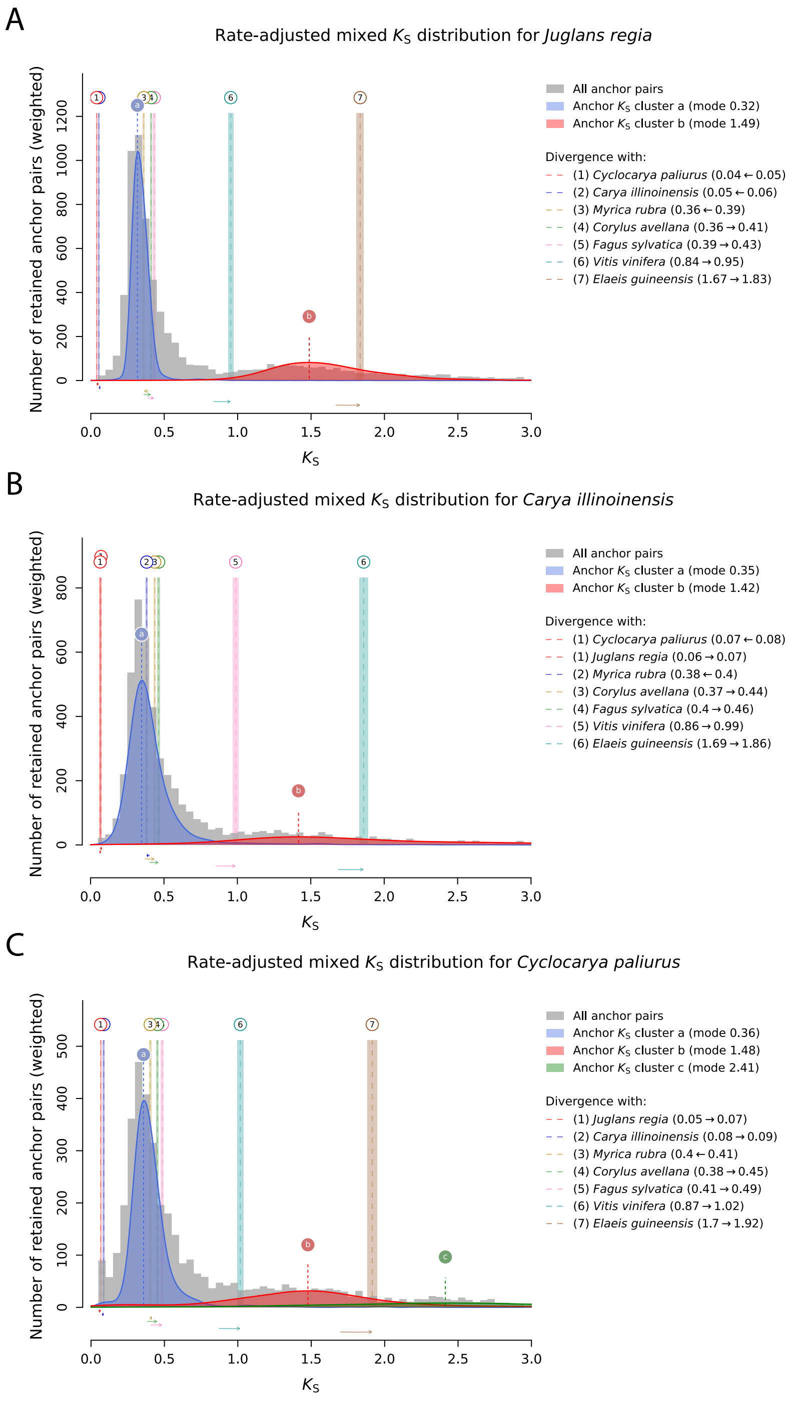


Figure S14: Substitution rate-adjusted mixed anchor pair $K_S$ distributions generated with `ksrates` for WGM inference and positioning in the Fagales. Anchor pair clusters delimiting WGM peaks are displayed as filled coloured areas. The modes of the WGM peaks are shown as vertical lines labeled with letters. Vertical lines labeled with numbers indicate the modes of ortholog $K_S$ distributions, which mark the divergence between the focal species and other species. Colored boxes range from one standard deviation (SD) below to one SD above the mean mode estimate. The $K_S$ ages of ortholog $K_S$ modes have been adjusted on the basis of substitution rate differences between species, as illustrated by the horizontal arrows above the x-axis. (A) Clustered anchor pair $K_S$ distribution for *Juglans regia*. (B) Clustered anchor pair $K_S$ distribution for *Carya illinoinensis*. (C) Clustered anchor pair $K_S$ distribution for *Cyclocarya paliurus*. All three species share the recent Juglandaceae WGD (peak `a` in (A), (B), (C)) and the eudicot $\gamma$ WGT (peak `b` in (A), (B), (C))

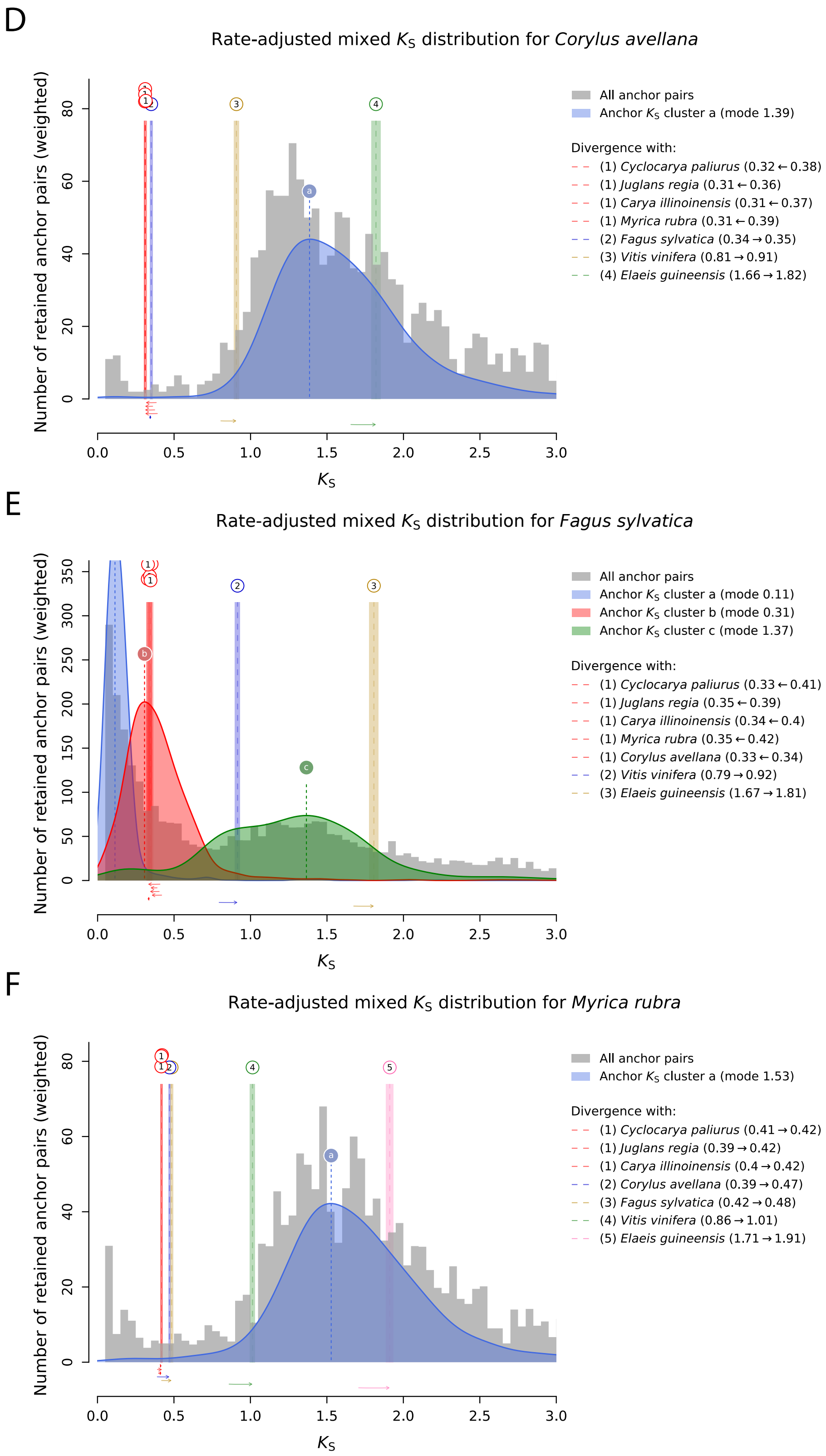


Figure S14: (continued) (D) Clustered anchor pair $K_S$ distribution for *Corylus avellana*. (E) Clustered anchor pair $K_S$ distribution for *Fagus sylvatica*. (F) Clustered anchor pair $K_S$ distribution for *Myrica rubra*. There is only evidence for the eudicot $\gamma$ WGT in these three species (peaks `a` in (D) and (F), peak `c` in (E)). Clusters `a` and `b` in (E) are likely SSD noise in the anchor pair distribution for *Fagus sylvatica*.

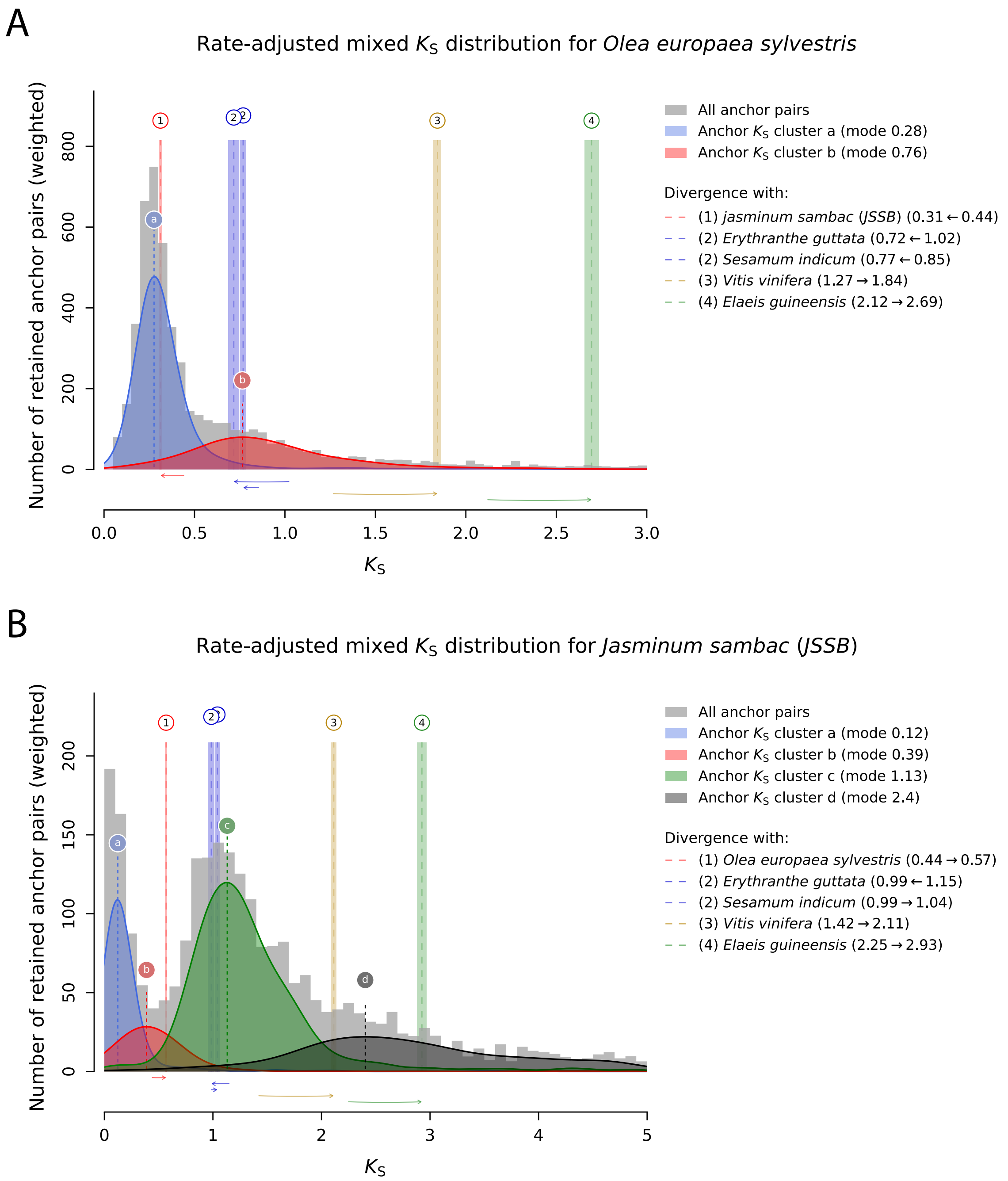


Figure S15: Substitution rate-adjusted mixed anchor pair $K_S$ distributions generated with `ksrates` for WGM inference and positioning in the Lamiales. (A) Clustered anchor pair $K_S$ distribution for *Olea europea sylvestris*. (B) Clustered anchor pair $K_S$ distribution for *Jasminum sambac (JSSB)*. Anchor pair clusters delimiting WGM peaks are displayed as filled coloured areas. The modes of the WGM peaks are shown as vertical lines labeled with letters. Vertical lines labeled with numbers indicate the modes of ortholog $K_S$ distributions, which mark the divergence between the focal species and other species. Colored boxes range from one standard deviation (SD) below to one SD above the mean mode estimate. The $K_S$ ages of ortholog $K_S$ modes have been adjusted on the basis of substitution rate differences between species, as illustrated by the horizontal arrows above the x-axis. Peak `a` in (A) indicates a recent WGD specific to Oleeae, while peak `b` in (A) and peak `c` in (B) indicate an older WGT in the Oleaceae lineage. This WGT appears to have happened around (A) or just before (B) the split with non-Oleaceae species *S. indicum* and *E. guttata*, but the lack of a contemporaneous WGM peak in the $K_S$ distributions of the latter species (see (C) and (D) below) indicates that the Oleaceae WGT happened right after this split instead. Peaks `a` and `b` in (B) are likely SSD noise or assembly errors leaking through to the anchor pair distribution. Peak `d` in (B) corresponds to the core eudicot $\gamma$ WGT.

C

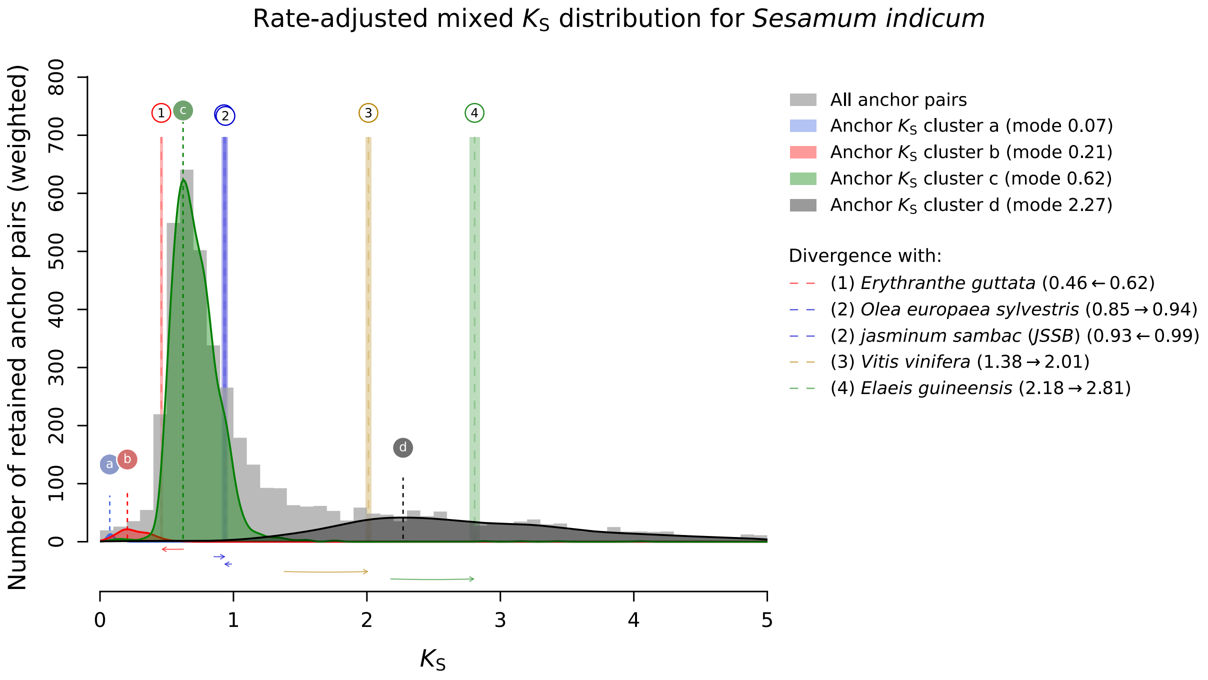


D

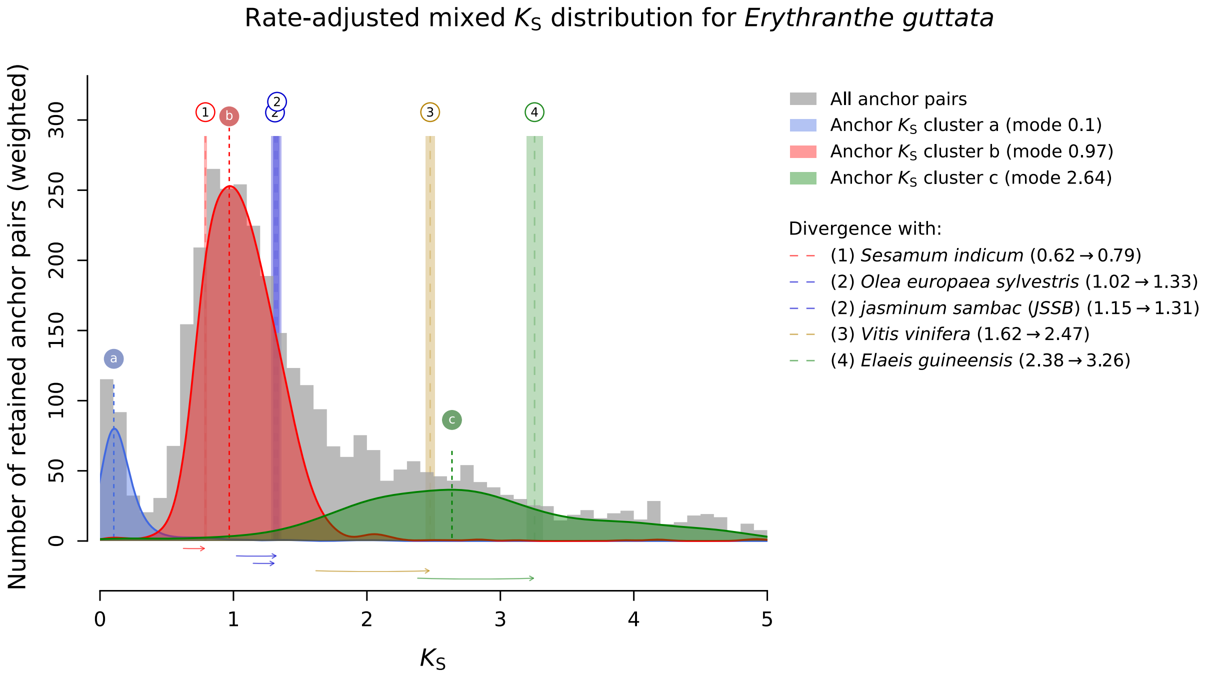


Figure S15: (continued) (C) Clustered anchor pair $K_S$ distribution for *Sesamum indicum*. (D) Clustered anchor pair $K_S$ distribution for *Erythranthe guttata*. A recent WGM peak shared by *S. indicum* (peak `c`) and *E. guttata* (peak `b`) dated after the divergence of *O. europea* and *J. sambac* corresponds to the Pedaliaceae-Phrymaceae WGD. Peaks `d` in (C) and `c` in (D) correspond to the core eudicot $\gamma$ WGT.

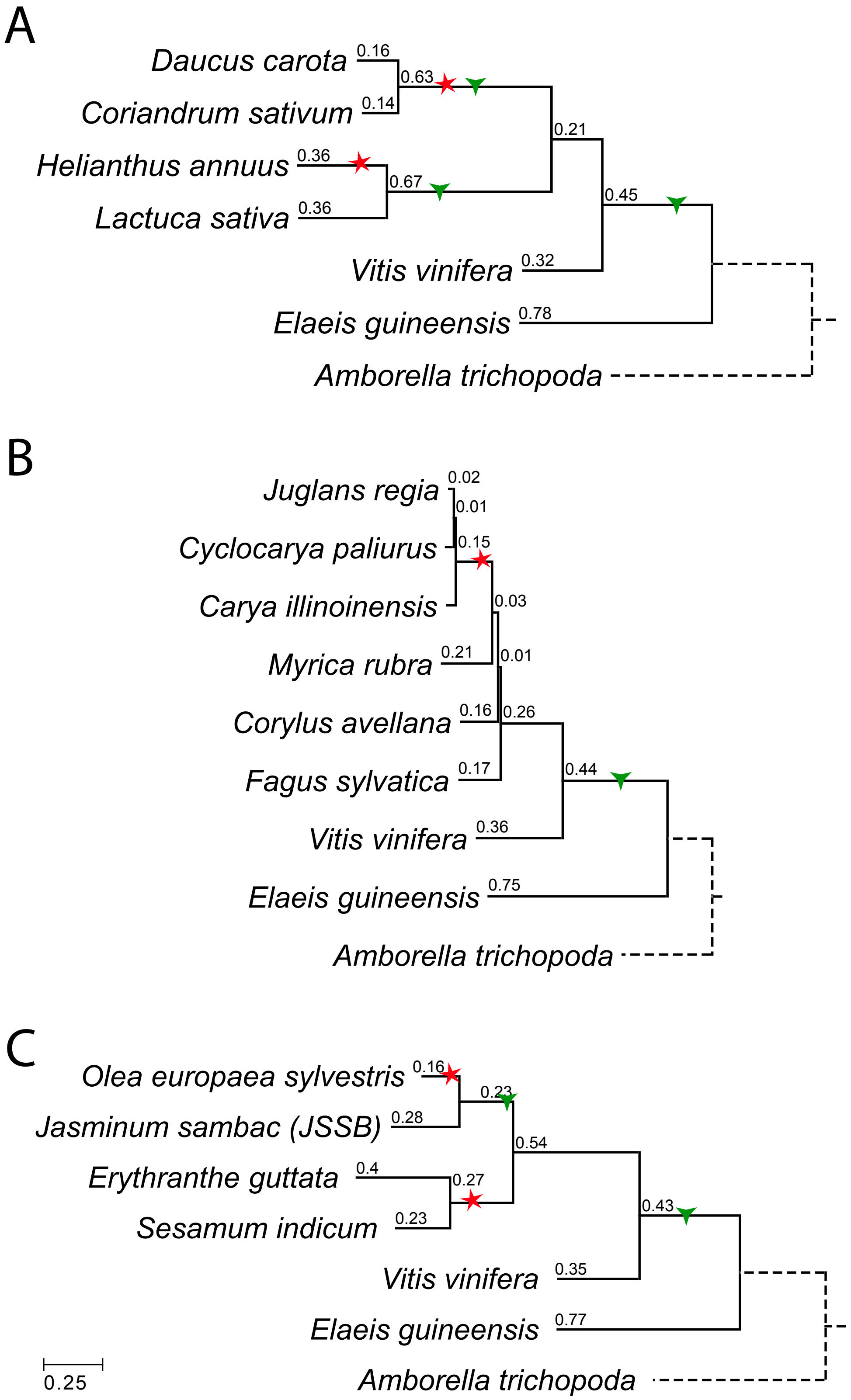


Figure S16: Phylogenetic trees produced by *ksrates* for Campanulids (A), Fagales (B) and Lamiales (C). Branch lengths expressed in $K_S$ units are indicated on the branches. WGMs are manually positioned on the trees, with WGDs marked as red five-pointed stars and WGTs as green three-pointed stars.

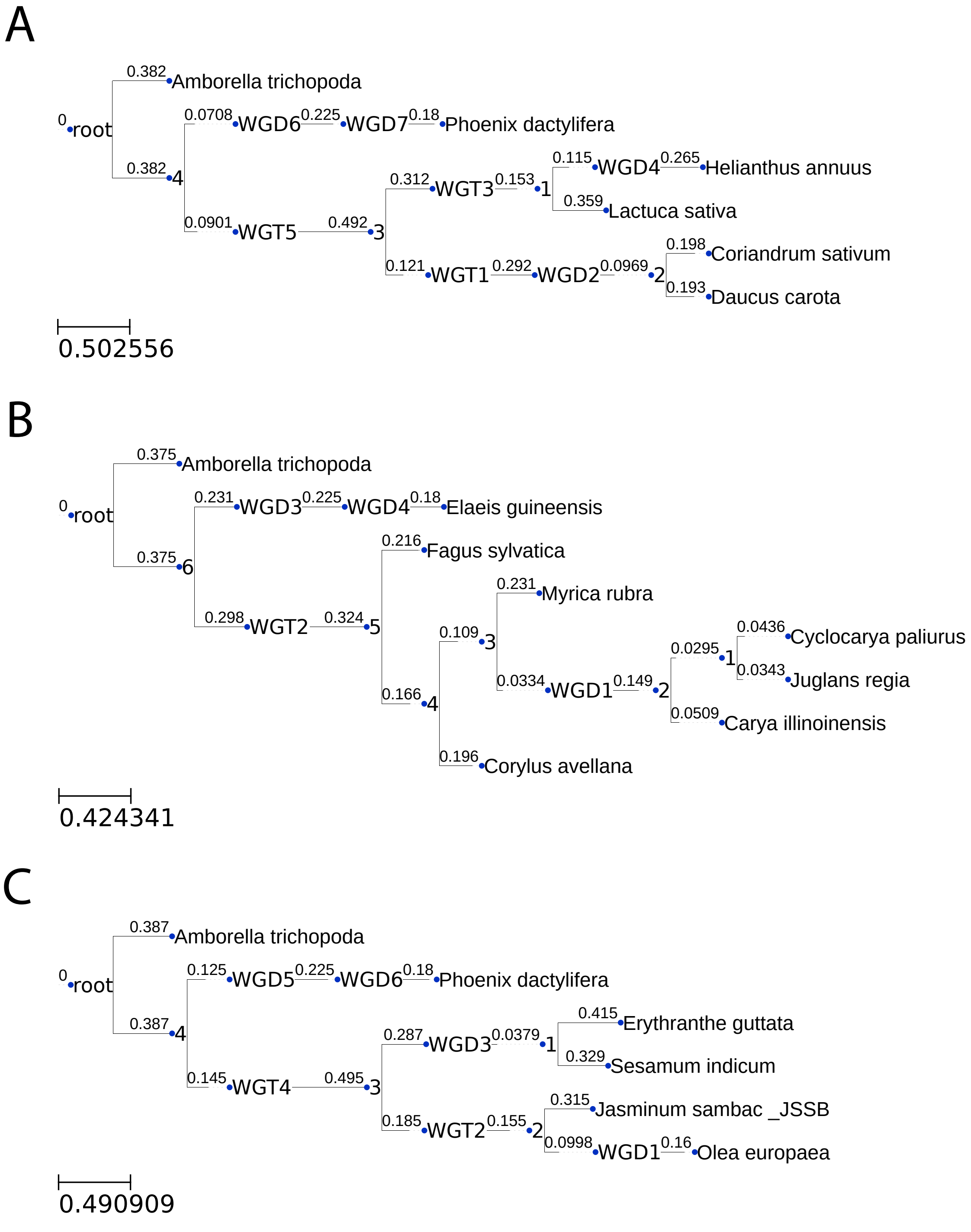


Figure S17: Phylogenetic trees of additional clades provided to the cLRT pipeline for gene count modeling. (A) Campanulids clade. WGT1 is the Apioideae WGT, WGD2 is the Apioideae WGD, WGT3 is the Asteraceae WGT, WGD4 is the Helianthus WGD, WGT5 is the core-eudicot $\gamma$ WGT, WGD6 is the monocot $\tau$ WGD, and WGD7 is the Arecaceae WGD. (B) Fagales clade. WGD1 is the Juglandaceae WGD, WGT2 is the core-eudicot $\gamma$ WGT, WGD3 is the monocot $\tau$ WGD and WGD4 is the Arecaceae WGD. (C) Lamiales clade. WGD1 is the Oleeae WGD, WGT2 is the Oleaceae WGT, WGD3 is the Pedaliaceae-Phrymaceae WGD, WGT4 is the core-eudicot $\gamma$ WGT, WGD5 is the monocot $\tau$ WGD and WGD6 is the Arecaceae WGD. Branch lengths are given in terms of numbers of substitutions per codon (t).

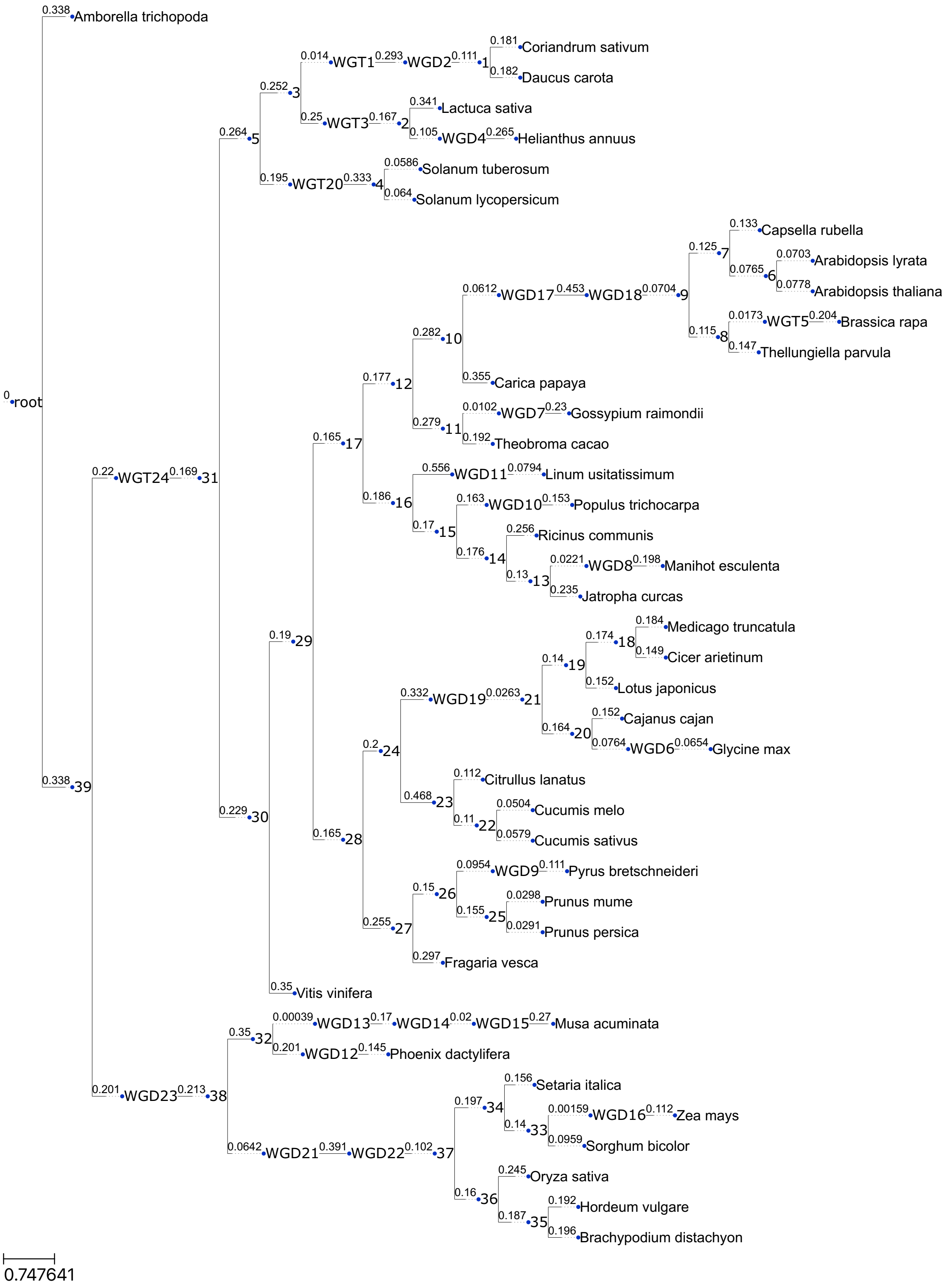


Figure S18: Phylogenetic tree obtained with IQ-TREE2 for the 37-species dataset plus Campanulids. Branch lengths in terms of numbers of substitutions per codon (t) are reported in black. WGM positions are marked with numbered WGD or WGT nodes, see Suppl. Data S5D for the correspondence between WGM numbers and WGM names.

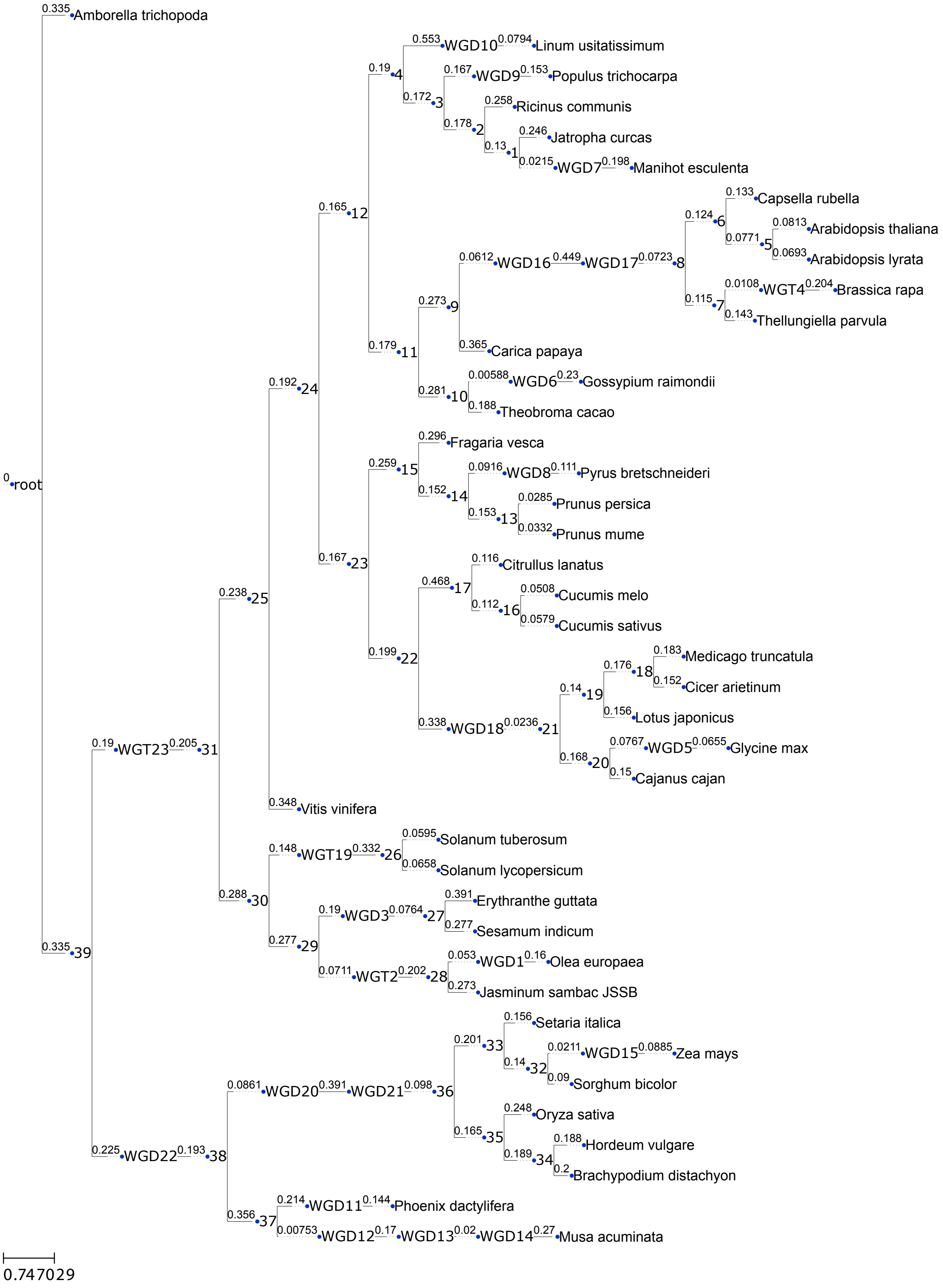


Figure S19: Phylogenetic tree obtained with IQ-T REE2 for the 37-species dataset plus Lamiales. Branch lengths in terms of numbers of substitutions per codon (t) are reported in black. WGM positions are marked with numbered WGD or WGT nodes, see Suppl. Data S5H for the correspondence between WGM numbers and WGM names.

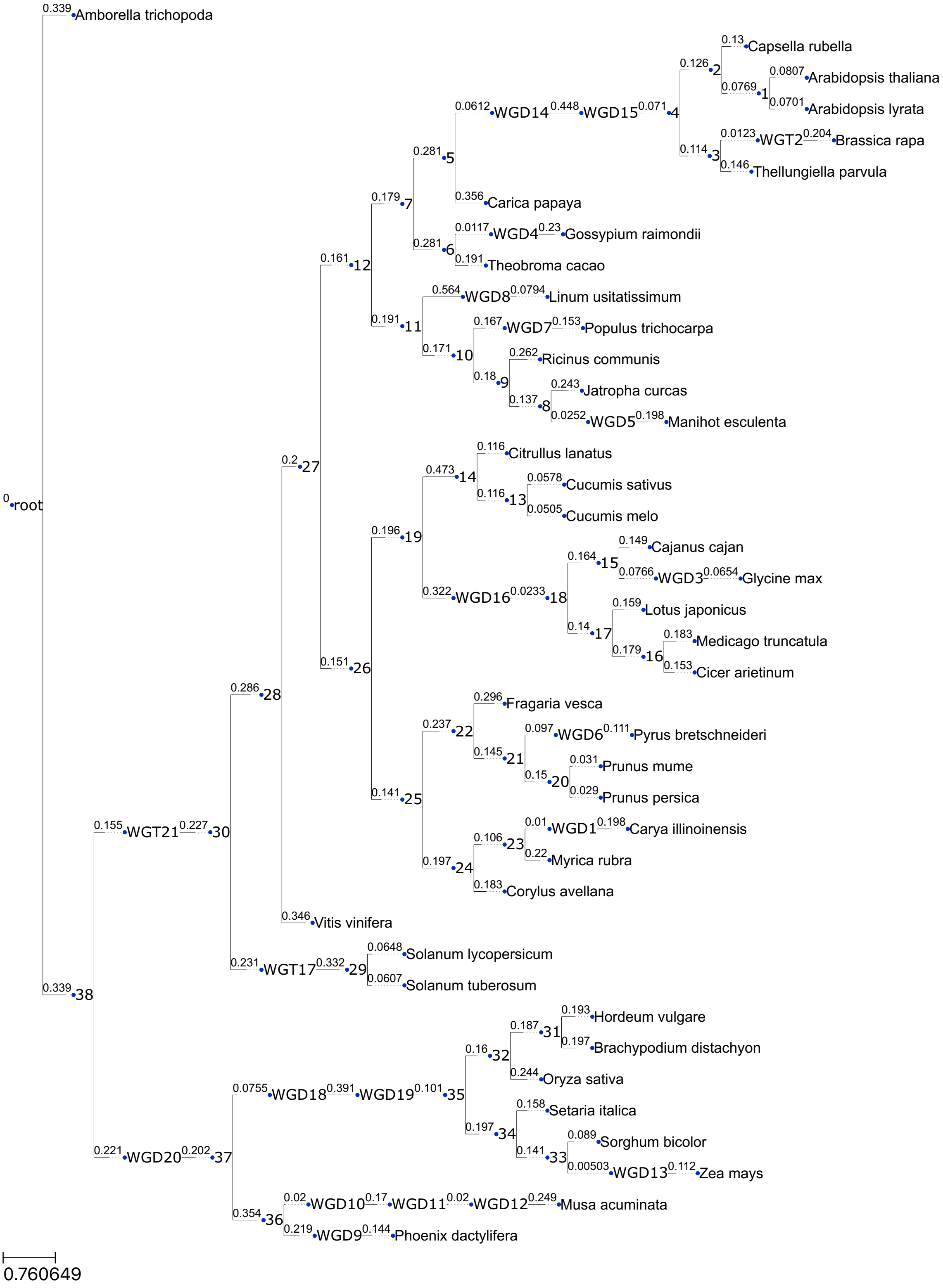


Figure S20: Phylogenetic tree obtained with IQ-TREE2 for the 37-species dataset plus Fagales. Branch lengths in terms of numbers of substitutions per codon (t) are reported in black. WGM positions are marked with numbered WGD or WGT nodes, see Suppl. Data S5F for the correspondence between WGM numbers and WGM names.

### 3 Supplementary Tables

| Full name | Source |
|---|---|
| Brassiceae WGT | Cheng *et al.* (2014) |
| Glycine WGD | Schmutz *et al.* (2010) |
| Gossypium WGD | Paterson *et al.* (2012) |
| Manihot-Hevea WGD | Bredeson *et al.* (2016) |
| Maleae WGD | Li *et al.* (2019) |
| salicoid WGD | Yates *et al.* (2021) |
| Linum WGD | Yuan *et al.* (2021) |
| Arecaceae WGD | Wang *et al.* (2022a) |
| Musaceae $\alpha$ WGD | Wang *et al.* (2023) |
| Musaceae $\beta$ WGD | Wang *et al.* (2023) |
| Musaceae $\gamma$ WGD | Wang *et al.* (2023) |
| Tripsacinae WGD | Gault *et al.* (2018) |
| Brassicaceae $\alpha$ WGD | Mabry *et al.* (2020) |
| (core) Brassicales $\beta$ WGD | Mabry *et al.* (2020) |
| Faboideae WGD | Young *et al.* (2011) |
| Solanaceae WGT | Tomato Genome Consortium (2012) |
| Poaceae-$\rho$ WGD | McKain *et al.* (2016) |
| Poales-$\sigma$ WGD | McKain *et al.* (2016) |
| monocot-$\tau$ WGD | Jiao *et al.* (2014) |
| core eudicot $\gamma$ WGT | Jiao *et al.* (2012) |

Table S1: List of the 20 well-supported WGMs. The full name of a WGM is composed of the name of the clade (most likely) sharing the WGM based on literature, followed by the WGM type (WGD or WGT). The 'WGD' in the Gossypium lineage was most likely a pentaploidization (2.5-fold increase of haploid gene counts, Paterson *et al.* (2012)), but as our modeling framework can only handle integer-fold increases of haploid gene counts, it was treated as a tetraploidization (2-fold increase of haploid gene counts).

| Abbreviation | Full name | Distance from parent node (t) |
|---|---|---|
| Cpap | *Carica papaya* WGD | 0.20 |
| Euphorbiaceae | Euphorbiaceae WGD | 0.05 |
| Rcom | *Ricinus communis* WGD | 0.21 |
| Eurosids | Eurosids WGD | 0.02 |
| Ljap | *Lotus japonicus* WGD | 0.13 |
| Fabaceae-Cucurbitales | Fabaceae-Cucurbitales WGD | 0.01 |
| Cucurbitales | Cucurbitales WGD | 0.02 |
| Rosales | Rosales WGD | 0.11 |
| Vvin | *Vitis vinifera* WGD | 0.22 |
| Pooideae | Pooideae WGD | 0.07 |

Table S2: List of the 10 unsupported WGDs used to counter-test the top and bottom gene families. Their positioning is expressed as the t distance from the parent node of their branch.

| Clade | Species | Version | Source | Publication |
|---|---|---|---|---|
| Campanulids | *Daucus carota* | JGI v2.0 | Plaza Dicots 5.0 | (Iorizzo *et al.*, 2016) |
| | *Coriandrum sativum* | – | Coriander Genome DB | (Song *et al.*, 2020) |
| | *Helianthus annuus* | r1.2 | Plaza Dicots 5.0 | (Badouin *et al.*, 2017) |
| | *Lactuca sativa* | v8.0 | Plaza Dicots 5.0 | (Reyes-Chin-Wo *et al.*, 2017) |
| Fagales | *Juglans regia* | Chandler v2.0 | GigaDB | (Marrano *et al.*, 2020) |
| | *Carya illinoinensis* | – | GigaDB | (Huang *et al.*, 2019) |
| | *Cyclocarya paliurus* | – | Genome Warehouse | (Zheng *et al.*, 2021) |
| | *Fagus sylvatica* | v2 | Beech genome | (Mishra *et al.*, 2022) |
| | *Corylus avellana* | v1.0 | Plaza Dicots 5.0 | (Pavese *et al.*, 2021) |
| | *Myrica rubra* | – | *Myrica rubra* Database | (Ren *et al.*, 2019) |
| Lamiales | *Olea europaea* var. *sylvestris* | – | Olive genome | (Unver *et al.*, 2017) |
| | *Jasminum sambac (JSSB)* | – | – | (Wang *et al.*, 2022c) |
| | *Sesamum indicum* | v3 | – | (Wang *et al.*, 2022b) |
| | *Erythranthe guttata* | v2.0 | Plaza Dicots 5.0 | (Ibarra-Laclette *et al.*, 2013) |
| Outgroup | *Elaeis guineensis* | EG5.1 | Plaza Monocots 5.0 | (Singh *et al.*, 2013) |

Table S3: Genome assemblies and related publications used to generate the input for the cLRTs for the additional clades Campanulids, Fagales and Lamiales. For the outgroups *A. trichopoda* and *P. dactylifera*, the genome versions were the same as reported in Li *et al.* (2016).

| Clade | WGM name | Type | mya | Species |
|---|---|---|---|---|
| Campanulids | Apioideae | WGT | 54–61 | *D. carota*, *C. sativum* |
| | Apioideae | WGD | 45–52 | *D. carota*, *C. sativum* |
| | Asteraceae | WGT | 40–45 | *H. annuus*, *L. sativa* |
| | Helianthus | WGD | 29 | *H. annuus* |
| Fagales | Juglandaceae | WGD | 66 | *J. regia*, *C. illinoinensis*, *C. paliurus* |
| Lamiales | Oleaceae | WGT | 57–63 | *O. europaea*, *J. sambac* |
| | Oleeae | WGD | 26–30 | *O. europaea* |
| | Pedaliaceae-Phrymaceae | WGD | 50–55 | *E. guttata*, *S. indicum* |

Table S4: WGMs previously found in the Campanulids, Fagales and Lamiales clades. For related publications, see Table S3. mya: million years ago.
